# The open-source Masala software suite: Facilitating rapid methods development for synthetic heteropolymer design

**DOI:** 10.1101/2025.07.02.662756

**Authors:** Tristan Zaborniak, Noora Azadvari, Qiyao Zhu, S.M. Bargeen A. Turzo, Parisa Hosseinzadeh, P. Douglas Renfrew, Vikram Khipple Mulligan

## Abstract

Although canonical protein design has benefited from machine learning methods trained on databases of protein sequences and structures, synthetic heteropolymer design still relies heavily on physics-based methods. The Rosetta software, which provides diverse physics-based methods for designing sequences, exploring conformations, docking molecules, and performing analysis, has proven invaluable to this field. Nevertheless, Rosetta’s aging architecture, monolithic structure, non-open source code, and steep development learning curve are beginning to hinder new methods development. Here, we introduce the Masala software suite, a free, open-source set of C++ libraries intended to extend Rosetta and other software, and ultimately to be a successor to Rosetta. Masala is structured for modern computing hardware, and its build system automates the creation of application programming interface (API) layers, permitting Masala’s use as an extension library for existing software, including Rosetta. Masala features modular architecture in which it is easy for novice developers to add new plugin modules, which can be independently compiled and loaded at runtime, extending functionality of software linking Masala without source code alteration. Here, we describe implementation of Masala modules that accelerate protein and synthetic peptide design. We describe the implementation of Masala real-valued local optimizers and cost function network optimizers that can be used as drop-in replacements for Rosetta’s `minimizer` and `packer` when designing heteropolymers. We explore design-centric guidance terms for promoting desirable features, such as hydrogen bond networks, or discouraging undesirable features, such as unsatisfied buried hydrogen bond donors and acceptors, which we have re-implemented far more efficiently in Masala, providing up to two orders of magnitude of speedup in benchmarks. Finally, we discuss development goals for future versions of Masala.

## 1 Introduction

Computational design methods have revolutionized heteropolymer engineering. Since the early 2000s, proteins have been designed with physics-based software tools like the Rosetta software suite [1, 2] to fold into new structures not found in nature [3, 4], to self-assemble into oligomers [5], nanosheets [6], and nanocages [7–9], to bind to target proteins [10–12] and small molecules [13], and even to catalyze chemical reactions that no natural enzyme can catalyze [14–17]. Around 2016, the generalization of the Rosetta energy function and the introduction of unbiased sampling methods, such as generalized kinematic closure (GeneralizedKIC) and parametric methods [18–20], carried computational design into the realm of synthetic heteropolymers made from natural and non-natural chemical building-blocks. There are now many examples of computationally-designed mixed-chirality peptide macrocycles, ranging from 7 to 60 amino acids, which fold into rigid structures verified by x-ray crystallography or nuclear magnetic resonance spectroscopy (NMR) [18, 19, 21, 22]. These have been engineered to have diverse functions, with some binding metals and acting as metal-triggered conformational switches [22], others diffusing across cell membranes [23], and others binding to targets of therapeutic interest, such as the New Delhi metallo-*β*-lactamase 1 (NDM-1) or various histone deacetylases (HDACs) [24, 25]. Synthetic peptide macrocycles are also structurally fascinating, since they can posses internal symmetry, and, when synthesized from mixtures of `L`- and `D`-amino acids, can access symmetries involving mirror operations inaccessible to natural proteins made exclusively from `L`-amino acids.

Thanks to the vast databases of protein sequences and structures, the advent of machine learning has revolutionized the design and validation of proteins built from natural amino acids [26–30]. Although notable attempts have been made to use ML models to design peptide macrocycles [31, 32], these are hindered by the paucity of training data for exotic chemical building-blocks, and tend to produce peptides that closely resemble natural proteins in their chemical composition (such as all-`L` peptide macrocycles), or which fail to capture the unique conformational preferences of more exotic building-blocks [33]. ML models are best at interpolating and finding variations near to their training data, meaning that to go beyond the canonical 20 amino acids, general algorithms and physics-based methods are still needed, as is experimentation with new computational approaches.

Sadly, the Rosetta software suite, which for over two decades has been at the forefront of protein design [34], and which permitted pioneering work in non-canonical peptide macrocycle design (reviewed in [35]), is starting to show its age. Rosetta’s protein-centric assumptions hinder development of general methods for non-canonical heteropolymer design. Its poor support for modern multi-core CPUs, and its lack of support for GPUs, FPGAs, or other modern computing hardware, prevent users from making efficient use of modern computing clusters. The complexity of Rosetta’s library architecture also hinders new methods development, particularly by biologists and biochemists who may have greater scientific expertise than software engineering expertise, limiting its ability to be the platform in which one may implement and experiment with new noncanonical peptide modelling approaches. Using Rosetta in conjunction with other software is also difficult: the lack of a Rosetta application programming interface (API) prevents the powerful inner modules of Rosetta from being used directly by other software, and the closed-source licence hinders the free flow of code to and from other codebases, preventing scientific advancements from cross-pollinating. Although a Rosetta replacement is badly needed, the fact that Rosetta is actively being used for modelling cyclic peptides and proteins, both in many academic labs and in industry settings, is an obstacle to the needed rewrite. The danger is a forking of the Rosetta project: during a lengthy rewrite, developers’ efforts would be divided between maintaining the existing Rosetta code and implementing the replacement. New features added to Rosetta during this time may be lost once the replacement was introduced; alternatively, new features added to the replacement might not be usable for a long time.

In this chapter, we introduce the Masala software suite, which is intended to be a successor to Rosetta for designing, docking, and predicting structures of canonical proteins and exotic non-canonical heteropolymers, including mixed-chirality peptide macrocycles. We address the problem of developing a replacement for Rosetta while Rosetta is being actively used by building a general scientific software platform that can serve both as standalone software, or as an extension library for other software. By linking Rosetta against Masala, we can focus on adding new features to Masala, getting the benefit of them in Rosetta during Masala’s development, while retaining access to them once Masala matures into a standalone software package. Masala also addresses a number of Rosetta’s shortcomings. Unlike Rosetta, which has protein-centric assumptions hard-coded in its core, Masala’s Core library is deliberately written to be completely agnostic to the type of chemistry being described, with methods intended to apply to only one type of heteropolymer encapsulated in specialized plugin libraries. Unlike Rosetta, which was structured with low-memory compute nodes with a single CPU core in mind, Masala is constructed around the single instruction, multiple data (SIMD) paradigm, with all modules operating on vectors of input structures or chemical data to most efficiently take advantage of modern computing hardware (multi-core CPUs, GPUs, FPGAs, *etc.*). Perhaps the most innovative new feature, intended to address the difficulty of learning to contribute code to Rosetta, is Masala’s plugin-based architecture, which offers a low barrier to entry into Masala development by permitting developers to write standalone Masala modules in independently-compiled Masala plugin libraries that can be linked at runtime. This means that a novice developer can write and compile a small, self-contained module and have it integrate itself into a larger software project without need for any modifications to the larger project. Related to this, where Rosetta had no means of defining an API, Masala makes it easy for relatively inexperienced developers to define stable APIs for their modules with just a few lines of code. And in contrast to Rosetta’s closed licence, Masala’s free, open-source AGPL 3.0 licence maximizes its scientific utility and promotes the sharing of techniques across projects and disciplines.

This chapter is organized as follows. Section 2 gives an overview of the strengths and weaknesses of Rosetta, and the features of Masala that reflect what we learned from decades of Rosetta development. Section 3 is intended primarily for users, and it describes how external software may link Masala’s Core library at compilation time and load Masala plugin libraries (such as the Standard Masala Plugins library) at runtime. Included are examples of modifications to Rosetta modules, such as the `PackRotamersMover` and the `MinMover`, to permit Masala plugin optimizers not known at compilation time to be used, as well as an overview of the user experience, including examples of the Masala API definitions being used to automatically generate elements for the RosettaScripts user interface. Implementation details for developers are provided in the Appendices. Section 4 provides Masala computational benchmarks showing speed enhancements on a single CPU core, as well as efficient parallelization across many CPU cores for Rosetta design tasks. In section 5, we discuss the planned work to turn Masala from a support library for Rosetta and other software into fully-fledged standalone software. And finally, we summarize and conclude in section 6. Developers interested in writing their own Masala plugins or in modifying other software to use Masala plugins are directed to the Appendices for details of coding conventions (Appendix A, section 7), the contents of the Masala Core library (Appendix B, section 8), the contents of the Standard Masala Plugins library (Appendix C, section 9), and the integration of Masala with external software like Rosetta (Appendix D, section 10).

Version 1.0 of Masala’s Core library is made available in the `masala public` repository on Github (https://github.com/flatironinstitute/masala_public). The Standard Masala Plugins library, version 1.0, is also available from the `masala public standard plugins` Github repository (https://github.com/flatironinstitute/masala_public_standard_plugins). We host Doxygen source code documentation for the Masala Core library and for the Standard Masala Plugins library online at https://users.flatironinstitute.org/~vmulligan/. We have also publicly released our changes to Rosetta as a free and open-source BSD 3-clause licensed patch (https://github.com/flatironinstitute/masala_rosetta_patch_public).

## 2 Software overview

This section is divided into two subsections. First, in section 2.1, we review the strengths and weaknesses of the existing state of the art for general heteropolymer modelling (including cyclic peptide design): the Rosetta software suite. Then, in section 2.2, we introduce the features of the new Masala software suite that are intended to address the current shortcomings of Rosetta, while still permitting its best features to be preserved.

### 2.1 The Rosetta software suite

The Rosetta software suite is a set of C++ libraries and software applications for heteropolymer structural modelling. Originally developed in the mid 1990s as a set of FORTRAN libraries for protein structure prediction, Rosetta was subsequently adapted for protein design. The original FORTRAN code, though efficient, was not particularly adaptable, so various attempts to modernize the codebase were made. Automated translation to C++ gave rise to Rosetta 2 (then called Rosetta++) in 2005; however, this proved to be an unwieldy codebase with FORTRAN-style organization but C++ syntax, which did not take proper advantage of the object-oriented nature of C++ [34]. In 2009, Rosetta 3 was released. This version had been manually rewritten from scratch in C++ and given a more modular architecture [1], and this permitted a period of rapid expansion in the capabilities of the software. New methods for design and molecular docking, tools for building symmetric quaternary structures, specialized protocols for antibodies and for enzymes, and advanced analysis modules were added by an ever-growing community of Rosetta developers, in laboratories scattered across the world [34]. Over time, a rigorous system of code review, testing, and benchmarking was developed to ensure that the software remained stable and reliable (reviewed in [36]). Additionally, much work was done to begin to generalize Rosetta to permit it to model and compute energies of heteropolymers other than proteins, including nucleic acids, branched glycan chains, synthetic peptides built from mixtures of `D`- and `L`-amino acids, peptide macrocycles, and oligopeptoids [18–20, 37–40].

At its core, Rosetta consists of a kinematic engine which permits long chains of atoms to be manipulated by moving internal degrees of freedom (mainly dihedral angles, but also bond angles and bond lengths) [1]. This is in contrast to molecular dynamics (MD) simulators (reviewed in [41]), which typically permit atoms to move independently in Cartesian space, constrained by potentials that maintain internal degrees of freedom. Built atop Rosetta’s kinematic engine are various algorithms for exploring the vast conformational and sequence spaces of heteropolymers, and a scoring engine for computing energies given a particular sequence in a particular conformation. And built atop this are even more complex protocols that recombine simpler structure-manipulation operations to achieve particular design, structure prediction, or analysis tasks [1, 2, 20]. Because Rosetta uses primarily physics-based sampling and scoring approaches, it is extremely generalizable to exotic heteropolymers. Rosetta remains an important tool for modelling these molecules even in an age of machine learning (ML) methods trained on natural protein structures, which generalize more poorly to classes of molecules on which they were not trained [33]. For detailed chapters describing how to use Rosetta for cyclic peptide design, see references [42–44].

#### 2.1.1 Rosetta’s strengths

Due to its power and versatility, Rosetta is widely used for protein, peptide, nucleic acid, glycan, and general synthetic heteropolymer modelling. Dozens of academic laboratories are part of the Rosetta Commons [34], which actively develops and maintains Rosetta, and many more use Rosetta through the free non-commercial use licence. Additionally, dozens of corporations licence Rosetta for commercial use. Given the large user base, any project aiming to produce a successor to Rosetta must preserve Rosetta’s strengths. These strengths include its scientific performance (particular for non-canonical heteropolymer modelling), its core combinatorial optimizer for sequence design, its kinematic model, and its general conformational sampling tools for non-canonical heteropolymers and macrocycles. Atop all of this, Rosetta’s scripting languages permit the many powerful tools in Rosetta to be recombined for new problems. All of these features are discussed in greater detail below.

##### Scientific performance for non-canonical design and validation

For many decades, until recent advancements in ML-based protein structure prediction, Rosetta had been one of the top tools for protein structure prediction, consistently ranking among the top methods in the Critical Assessment of protein Structure Prediction (CASP) competitions [45–49]. It was also unsurpassed for protein design, producing the first fully *de novo* designed protein with a novel fold [3] and proteins with many novel functions [5, 7–9, 12–17]. The `ref2015` energy function in particular remains one of the most accurate force fields for protein modelling [20, 39]. Even in the ML era, Rosetta continues to play an essential role in many protein design projects, permitting high-accuracy structural modelling, analysis, and refinement of ML-designed sequences [28, 50].

Despite the encroachment of ML methods like Alphafold [27] for canonical proteins, in the realm of synthetic heteropolymer design, Rosetta remains unsurpassed. Rosetta has been used to design peptide macrocycles built from mixtures of natural `L`-amino acids, synthetic `D`-amino acids, and exotic chemical cross-linkers, with sizes ranging from 7 to 60 amino acid residues, that fold into rigid structures, bind metals, switch conformation, or diffuse across cell membranes [18, 19, 21–23, 35] – tasks that ML methods are unlikely to be well-suited to doing, for want of training data. Perhaps most importantly, Rosetta has been used to design peptide macrocycles that can bind to targets of therapeutic interest, such as the New Delhi metallo-*β*-lactamase 1 (NDM-1) and various histone deacetylases (HDACs) [24, 25]. Unlike peptide macrocycles produced by blind screening approaches, Rosetta design methods produced macrocycles that could bind to a desired site on the target, and in a desired binding conformation and relative orientation to the target. In the case of HDACs, selectivity for HDAC2 and not for its close homologue HDAC6, or for HDAC6 over HDAC2, was achieved [25]. Moreover, Rosetta-based estimates of peptide macrocycles’ Δ*G* of folding, based on conformational sampling simulations using Rosetta’s `simple cycpep predict` application, were shown to correlate strongly with experimentally-measured *IC*_50_ values, making Rosetta an invaluable tool for ranking designs to produce shortlists for costly experiments [24].

As new methods are developed, preserving Rosetta’s unmatched accuracy and robustness in the realm of synthetic heteropolymer design will be crucial.

##### Efficient, general combinatorial optimization for heteropolymer sequence design

Sequence design is a very difficult computational challenge. Given a chain of *N* amino acids or other chemical building-blocks, with *B* possibilities for the building-block identity at each position, there are *B^N^* possible sequences. Searching this exponentially-scaling search space has been proven to be an NP-hard problem [51]. For a 100-amino acid protein made only from the 20 canonical amino acids, there are 20^100^, or approximately 10^130^, possible sequences, an astronomically large number that cannot be exhaustively enumerated on any computing hardware that has been built, or which ever could be built. (Note that the universe is estimated to contain 10^80^ atoms [52, 53]. Even if a single atom were used as computer memory to store each sequence, and all matter in the universe were devoted to the purpose, the available memory for exhaustive enumeration would fall short by a factor of 10^50^.) Among these sequences, the vast majority would not fold at all, and the vast majority of the small subset that do fold would adopt folds different from the desired fold. Only an insignificantly tiny subset would fold into the desired structure compatible with the desired function. When non-canonical amino acids or other exotic chemical building blocks are used, *B* can grow from 20 to hundreds or thousands, vastly increasing the size of the sequence space that must be searched. For precise, physics-based methods with atomic detail, the challenge is complicated by the fact that one must typically assign a sidechain conformation in addition to a chemical building-block identity (together called the *rotamer* ), and given ⟨K⟩ discretized sidechain conformations on average per building-block identity, there are B ⟨K⟩ possible rotamers per position and (B ⟨K⟩)^N^ possible rotamer assignments from which to choose. And complicating this even further is the need for a means of recognizing a “good” sequence: to do this properly, one must compute the energy gap between the desired conformation and all alternative conformations given the candidate sequence, which requires a simultaneous search of an astronomically large sequence space and an astronomically large conformation space. To date, no good method has been proposed for doing this: all current methods *either* focus on very small subsets of sequence space while attempting to search conformation space, *or* focus on small subsets of conformation space while attempting to search sequence space.

Of the many approaches devised for sequence design, MD-based approaches, such as alchemical free-energy methods, permit very accurate estimation of how the free energy change on folding changes (ΔΔ*G_f_* ) when one makes a single point mutation [54]. However, these these come at enormous computational cost for each point mutation considered, and do not permit the consideration of more than a small handful of sequence variants. As such, alchemical methods typically serve only as refinement techniques, not as full *de novo* design techniques. To accelerate the sequence search to allow far more sequences to be considered for *de novo* design, a common approximation is to attempt to find the sequence that minimizes the energy of the folded state, rather than maximizing the energy gap between the folded state and all alternative states. When this approximation is made, dead-end elimination methods, such as the `DEE/A*` method and its variants implemented in the Osprey software suite [55], or the related branch-and-bound methods offered by Toulbar2 [56–58], can be good methods for small design problems. These permit many hundreds of thousands of potential sequences to be implicitly considered by rejecting entire branches of the decision tree, and focusing the search on the subset of regions that provably contain the rotamer selection that globally minimizes an objective function. However, these methods rapidly reach a tractability ceiling as sequence lengths or rotamer counts grow large. ML methods such as ProteinMPNN [29], which are trained on the hundreds of thousands of known protein structures and the hundreds of millions of known protein sequences, can even more rapidly choose small subsets of sequence space on which to focus, at the cost of losing guarantees of finding the global minimum. However, these generalize poorly (or, in many cases, not at all) to building-blocks outside of those found in the training data, and are ill-suited for designing non-canonical peptides or other exotic heteropolymers.

Rosetta’s `packer`, one of the core Rosetta algorithms, uses highly efficient simulated annealing-based methods to find low-energy solutions to NP-hard sequence design problems [1, 59], with no loss of generality when non-canonical building-blocks are used. This has proved to be one of the most versatile methods for sequence design, as well as one of the fastest, permitting much larger problems than can be carried out by `DEE/A*` or branch-and-bound deterministic methods, at the cost of losing strong guarantees of finding the global optimum in order to find a “good enough” local optimum quickly. For efficiency, the `packer` divides design problems into a pre-computation phase and a search phase.

During the pre-computation, energy terms that are dependent on single chemical building blocks (one-body terms) or on pairs of interacting chemical building blocks (two-body terms) can be pre-calculated and cached, given a rigid backbone conformation [1]. Rosetta’s current default energy function, `ref2015`, is entirely composed of one- and two-body terms, meaning that it can be fully pre-computed [20]. This frees Rosetta from having to consider molecular geometry during the search phase. The one-body pre-calculation is an 𝒪 (B ⟨K⟩ N) calculation (*i.e.* it scales linearly with the total number of rotamers in the problem). The two-body pre-calculation is in the worst case an 𝒪 (B ⟨K⟩ N)^2^ calculation; however, in practice, because only local two-body interactions are considered and because most positions have on average a constant number of neighbouring residues in a large protein or other heteropolymer, this is effectively an 𝒪 (B ⟨K⟩)^2^ N calculation (*i.e.* it scales quadratically with the number of rotamers per position, and linearly with the number of positions). Because the pre-calculation is polynomial in complexity, it can be efficiently parallelized, and it is one of the few parts of Rosetta that is currently able to take advantage of multi-threading.

The search phase is the NP-hard phase. Any known approach to find the energetically-optimal rotamer selection shows poor scaling (𝒪 (B ⟨K⟩)*^N^* or worse), and Rosetta’s `packer` is no exception: simulated annealing approaches do not change the time-complexity scaling of the search phase. However, by avoiding any consideration of molecular geometry during this phase, and by relying only on rapid lookups of pre-computed one- and two-body scoring terms to calculate energy changes, Rosetta’s `packer` is able to consider on the order of hundreds of thousands to millions of unique sequences per second on a single CPU core, offering a large constant factor speedup over alternative methods despite not changing the scaling. The `packer` uses a simulated annealing search that starts from a “null” rotamer selection. Each move consists of choosing a position at random and substituting a new rotamer at that position at random, then computing the change in the objective function that results using the pre-computed lookup tables [59]. Moves that lower the value of the scoring function are always accepted; those that raise it are accepted or rejected with conditional probability *P_accept_* given by the Metropolis criterion shown in Eq. (1), where Δ*S* is the change in the value of the scoring function and *k_B_T* is a temperature. The temperature *k_B_T* ramps down in a stair-stepped fashion that approximates an exponential decay, then jumps back up; the default temperature range is 100 kcal/mol to 0.3 kcal/mol.

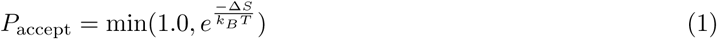

Note that Rosetta uses a slightly modified version of the general Metropolis criterion if the previous score, *S*_prev_, was greater than 1.0 kcal/mol. This promotes more acceptance of energy-raising moves at high scores, at the expense of assigning significance to the zero point of the score scale, reducing the generality somewhat. This is shown in Eq. (2)

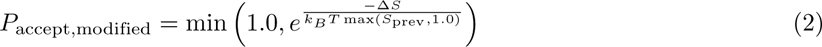

The details of the cooling schedule are set adaptively. Typically, at each “step” down in the stair-step, the temperature is set using Eq. (3):

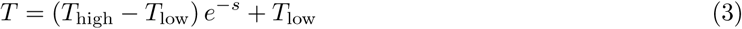

In the above, *s* is a counter incremented from zero with each “step”. If, after three such “steps”, the score hasn’t dropped by at least 1 kcal/mol compared to the recent average, the algorithm treats this as a stall, resets *s* to 0, and “re-heats” to *T*_high_. On the final “step” down, the temperature is automatically set to *T*_low_. For a given trajectory, the `packer` stores 1-dimensional integer arrays storing both the current state and the best state, allowing for return of the best state at the end of the trajectory.

Also note that the `packer`’s default behaviour is to omit the pre-calculation and to re-compute scores on the fly during the search phase (which requires explicit consideration of molecular geometry and slows the search by a factor of ten or more). For small problems, this is more efficient (*i.e.* the amount by which the search phase is lengthened is less than the cost of the pre-calculation), but as problems grow large, it becomes less efficient. Also note that additional design-centric guidance terms that are often not decomposable into one- and two-body terms can also be included [42, 43]. These must be computed on the fly during the search phase. These are further discussed in section 10.4 and in Appendix D, section 10.5.

Although the Rosetta `packer` is one of the most efficient and versatile means of designing sequences of exotic heteropolymers, it has its limitations. These include the fact that it is hard-coded to operate on chemical building block rotamers, preventing its use for more general problems with the same functional form, and that it is hard-coded to use simulated annealing, limiting developers’ ability to experiment with alternative means of solving design problems. These limitations are discussed further in section 2.1.2.

##### Effective propagation of derivatives for minimization using internal degrees of freedom

One of Rosetta’s strengths has been its kinematic representation of the protein backbone, which permits protein structures to be manipulated by rotating dihedral angles or altering other internal degrees of freedom (bond lengths, bond angles, or rigid-body transforms between chains) [1] Rosetta’s `AtomTree` class establishes a directed acyclic graph (DAG) describing the kinematic relationships between atoms, so that changes to degrees of freedom propagate to downstream atoms. This has enabled Rosetta’s backbone conformational sampling approaches, which permit much larger movements by directly altering dihedral values rather than relying on simple Cartesian-space translations of atoms [45]. Furthermore, Rosetta has the ability to propagate derivatives of energies with respect to Cartesian atom coordinates to compute derivatives of energies with respect to internal degrees of freedom, allowing gradient-descent minimization trajectories which, for instance, hold bond lengths, bond angles, and torsion angles for rigid bonds fixed while only allowing rotatable bonds to change. Rosetta’s `minimizer` module uses this to relax protein structures, permitting larger changes leading to deeper minima than could be found were gradient-descent minimization carried out in the much more rugged, higher-dimensional space of energy as a function of Cartesian coordinates of atoms [1].

##### General tools for heteropolymer and macrocycle conformational sampling

Heteropolymer conformational space is incredibly vast, necessitating clever algorithms for finding the rare regions that give rise to functional folds: the “needles” in the “haystack”. Historically, the prediction of protein structures and the rational design of new protein folds was achieved by fragment assembly methods [45], in which pieces of proteins of known structure were used to guide the search for the native state of a protein of known sequence but unknown structure, or to find “designable” conformations compatible with a desired function when crafting new proteins. These tools remain available in Rosetta; however, more recently, ML methods trained on all known protein structures have supplanted fragment-based approaches as the method of choice for exploring natural protein conformations. These include Alphafold, trRosetta, and RFDiffusion [27, 60–62]. Both fragment-based method and ML methods fall under the umbrella of prior knowledge-based approaches, which make the conformational sampling problem tractable by ruling out the vast majority of conformation space that does not resemble one’s prior expectation, informed by past observations, of what a protein “ought” to look like. Unfortunately, prior knowledge-based approaches are incompatible with the exploration of new synthetic heteropolymer conformations when few experimentally-determined structures exist for synthetic heteropolymers in the class of interest. For these, approaches that bias the search based on other forms of prior knowledge, such as the secondary structures accessible to a synthetic heteropolymer, or the geometric constraints of a mixed-chirality cyclic peptide, are needed.

Rosetta offers tools like the `MakeBundle`, `PerturbBundle`, and `BundleGridSampler` movers, which permit coiled-coil conformations of heteropolymers to be parametrically generated and sampled [19]. These are written to be general enough to support any helix type, not just the right-handed *α*-helix accessible to `L`-amino acids. Synthetic peptides and other heteropolymers can form diverse helix types, such as the left-handed *α*-helices accessible to `D`-amino acids, the 3_14_ helices formed by *β*-amino acids, or the type I and II polyproline helices formed by peptoids, and any of these could conceivably be designed to self-assemble into coiled-coil tertiary folds. Since the *β*-strand and other non-canonical strands represent a special case of a helix, in which the turn per residue is close to 180*^◦^*, the tertiary folds that can be sampled include folds like *β*-barrels. Rosetta’s parametric tools use coiled-coil generating equations developed by Francis Crick [63], which have user-controllable parameters such as bundle radius *r*_0_, bundle twist *ω*_0_, and individual helix roll Δ*ω*_1_ which can be sampled to continuously vary the coiled-coil tertiary fold. Parametric tools provide an essential means of exploring the tertiary structures of exotic peptides and other heteropolymers; however, these do have their limitations as well (see further discussion in Appendix B, section 8.3.1). One limitation is that, although in theory derivative propagation to construct gradients with respect to parametric degrees of freedom should be straightforward, the Rosetta implementation hinders this: since the parametric model does not directly replace Rosetta’s kinematic model, the Rosetta `minimizer` cannot be used for parameter-space relaxation. A Rosetta replacement should ideally provide a more versatile kinematic system, permitting experimentation with different parametric representations of conformation.

In addition to having means of sampling assemblies of secondary structures into tertiary structures, the design of synthetic peptides and other non-canonical heteropolymers depends on having means of sampling loop conformations; in the case of cyclic peptides, these can be used to sample macrocycle conformations. For this, Rosetta offers tools like `GeneralizedKIC` (short for “generalized kinematic closure”) [18], which uses kinematic methods borrowed from the robotics field to sample conformations of a chain of atoms with fixed start and end points subject to the constraint that the chain remains closed. Most degrees of freedom of the chain can be sampled in user-controllable ways (for instance, biased by the conformational preferences of particular canonical or non-canonical building-block types), after which a remaining six degrees of freedom (typically, six dihedral angles adjacent to three “pivot” atoms) are assigned values analytically in order to preserve the rigid-body transformation from the start of the chain to the end. `GeneralizedKIC` was a generalization for arbitrary chemical building blocks that was introduced to Rosetta in 2014, building on earlier kinematic closure approaches developed for chains of canonical amino acids [64, 65]. It has been used extensively to design cyclic peptides that fold [18, 19, 21], bind metal ions [22], or bind to targets [24, 25], and can also be used for sampling loop regions of linear heteropolymers that fold [66]. Unfortunately, `GeneralizedKIC`’s CPU implementation is less efficient than it could be given Rosetta’s one-structure-in, one-structure-out paradigm: the kinematic closure algorithm by its nature finds multiple closure solutions from each closure attempt, and often makes many closure attempts with different random perturbations to the non-pivot degrees of freedom resulting in a potentially lengthy candidate conformation list, but a single “best” solution must be returned prior to proceeding into subsequent steps of a protocol. In many structure prediction and design protocols, this means some unnecessary computational expense since any given sample is the product of independent (and often redundant) conformational sampling. A Rosetta successor better able to operate on batches of structures in tandem could make much more efficient use of kinematic closure approaches.

##### RosettaScripts and PyRosetta scripting languages

Early versions of Rosetta were controlled entirely at the command-line, or by providing specialized input files that configured options for particular protocols. Unfortunately, this meant that Rosetta modules could only be recombined in new ways to solve new problems by writing new C++ protocols. To address this, two Rosetta scripting interfaces were introduced. The first, PyRosetta, was introduced in 2009, and it provides Python bindings for all classes, their public member functions, and the non-class functions in Rosetta [67]. Since Python is a familiar language for many scientists, with a lower entry barrier and shallower learning curve than C++, this has permitted far more Rosetta users to develop their own protocols to solve new problems using existing Rosetta modules.

PyRosetta introduced some problems of its own, however. One was that it defined no application programming interface (API). Since the *entire* Rosetta C++ codebase was now a Python user interface, any change to any function signature, anywhere, could potentially result in an unknown number of users’ Python-scripted protocols breaking. The other was that certain performance-optimized features of Rosetta, such as its Message Passing Interface (MPI) based job distribution system, could not easily be used through PyRosetta.

In 2011, the complementary RosettaScripts scripting language was introduced [68]. RosettaScripts, unlike PyRosetta, is not intended to be Turing-complete. Instead, it provides a simple means of stringing together a series of Rosetta `Mover` objects, which alter structures in some way, and `Filter` objects, which analyse a structure and make decisions about whether to continue or abort a pass through the script, in order to assemble a linear protocol producing one sample for a design, structure prediction, docking, or other modelling task. Additional classes of scriptable objects, such as `SimpleMetrics` for analysis and `ResidueSelectors` for selecting parts of a structure when configuring other modules, were added later [2]. RosettaScripts permits samples to be efficiently parallelized using MPI-based job distribution, with each MPI process executing embarrassingly parallel instances of the scripted protocol to produce independent samples. Although, like PyRosetta, RosettaScripts initially lacked a definition of its interface, refactoring efforts in 2016 led to Rosetta being able to produce an XML schema definition (XSD) for the entire RosettaScripts language. This allowed better unit and regression testing of the XML interface, and greater stability from version to version of Rosetta. It also permitted automatic generation of documentation, and online help by running RosettaScripts with the `-info <MODULE name>` flag to output a human-readable description of the XML interface for a particular module. Rosetta’s ability to report its own interface in an XSD file also allows better interactivity with IDEs such as VSCode, which can read the RosettaScripts XSD to provide tab completion and mouse-over help to users writing RosettaScripts XML scripts. And because RosettaScripts tags can be provided to Rosetta modules in a PyRosetta context, the RosettaScripts XSD has provided a stable interface for *PyRosetta* as well. Indeed, this is now the preferred way to configure Rosetta modules when invoking them from PyRosetta.

The notion of having software that can produce a human- or machine-readable description of its own interface has proved to be revolutionary for Rosetta. As discussed in section 2.2.3 and in Appendix B, section 8.1.1, this is an idea that can be taken one step further in a Rosetta successor, if it can be used not only to describe a user interface for humans, but a machine interface for other code: an API allowing a Rosetta successor to be used as a library.

#### 2.1.2 Rosetta’s weaknesses and limitations

Rosetta has proven itself to be an invaluable tool for general heteropolymer design, structure prediction, docking, and analysis, and is central to design pipelines intended to make mixed-chirality cyclic peptides for medicine or other applications. Nevertheless, many weaknesses and limitations have become apparent over its lifetime. These include hard-coded assumptions about canonical proteins, poor support for modern parallel computing hardware such as multi-core CPUs, GPUs, or FPGAs, software architecture that is difficult to extend (particularly for scientists whose expertise may not lie in software engineering), the lack of an API, and the closed-source codebase. These are discussed here as important points that should be addressed in a possible successor to Rosetta.

##### Canonical protein-centric assumptions

Rosetta was originally developed to predict structures of (and later to design) proteins built from the 20 canonical amino acids [3, 45]. This has meant that Rosetta’s codebase is littered with protein-centric assumptions, not only in protocols specific for proteins, but in core infrastructure. In many places in the codebase, it was assumed that the number of building-block types would be known at compilation time, and that all cases could be covered exhaustively by conditional logic — a fine assumption given only 20 building-block types, but one which breaks down when working with hundreds or thousands of exotic non-canonical building blocks that can be incorporated into synthetic peptides. Extensive refactoring was needed to make Rosetta compatible with `D`-amino acids, to remove hard-coded assumptions about the number of backbone atoms in a building-block (fixed in *α*-amino acids, but not when one can potentially design with *β*- or *γ*-amino acids), or to permit non-canonical rotamer libraries [2, 18, 37, 38, 42, 43]. The energy function in particular had to be generalized to ensure that mirror-image structures score identically, that non-canonical mainchain and sidechain potentials (of possibly different dimensionality than those of the canonical amino acids) could be loaded from disk on demand, and that both potentials and their derivatives were properly interpolated when using non-canonical lookup tables [2, 20]. Rosetta’s symmetry code also had to be refactored to add support for mirror symmetry operations, which do not occur in canonical proteins made only of `L`-amino acids [22]. While enormous progress has been made towards fully general heteropolymer modelling software, even today, protein-specific assumptions do turn up as causes of bugs affecting non-canonical modelling tasks, and protein-centric architecture continues to be an obstacle to expansion of Rosetta to support more exotic heteropolymers.

##### Poor support for modern computing hardware

Rosetta’s development began in the mid-1990s, at a time when the dominant paradigm in computing was the single instruction, single datum (SISD) workflow, and when the memory on a compute node was very limited. Much of Rosetta is therefore based around operations performed sequentially on a single input structure (a `Pose` object, in Rosetta nomenclature), resulting in a single output structure. Protocols based on large-scale sampling clear memory and repeat the process with a second `Pose` object (with freshly-drawn random numbers for stochastic sampling protocols), keeping only one `Pose` in memory at any given time. Limited support was later added for parallelism using the message passing interface (MPI), but this still relied on each process (typically mapped to a single CPU core) operating on one `Pose` at a time for an entire multi-operation sampling protocol, before moving on to the next sample, with samples carried out asynchronously in different processes [69]. The one-`Pose`-at-a-time requirement is baked into Rosetta’s architecture both at the level of job distribution and in the operations available. Rosetta `Mover` subclasses, for instance, are modules which alter `Pose` objects, of which 539 have been implemented and made available to the RosettaScripts scripting language at the time of this writing. All `Mover` subclasses take one `Pose` as input and produce one `Pose` as output. The same is true of `Filter` subclasses, which analyse a single `Pose` to produce a pass or fail result, and of which there are 190 in Rosetta that are available in the RosettaScripts scripting language at the time of this writing.

Although the SISD approach minimizes demands on memory, it is inefficient on modern hardware. Modern compute nodes can have terabytes of memory, and modern processor architecture very efficiently supports single instruction, multiple data (SIMD) pipelines [70], meaning that sampling protocols would be better structured to operate on many `Pose` objects at once, performing the same operation on all before moving on to the next operation. Additionally, modern CPUs feature many levels of cache memory permitting faster lookups of small amounts of frequently-used information, meaning that if code architecture were constructed cleverly, it could be possible to minimize cache misses by holding a representation of only the salient parts of many structures in L1 cache, closest to the CPU core, and operating on those, rather than operating on many large `Pose` objects in their entirity requiring frequent retrievals of disparate data from main memory and producing frequent cache misses.

The rise of GPU computing (reviewed in [71]) has further revealed the limits of Rosetta’s current architecture. GPUs operate best when large amounts of work can be expressed as vectors of operations to perform concurrently, but this too is incompatible with Rosetta’s one-`Pose`-at-a-time approach: any given operation on a single Pose (even evaluation of the Rosetta energy function or computation of its gradient vector) involves too little vectorizable work to justify moving data to and from the GPU. The cost of moving the operation onto the GPU is greater than the time saved when the operation executes. Again, it would be far more efficient to operate on many `Pose` objects at once.

Finally, although Rosetta’s greatest strengths include its general, physics-based approach to molecular modelling, it cannot be denied that for particular problems (including sub-problems that must be solved in a heteropolymer design or analysis pipeline), ML methods are the best approach. A truly robust heteropolymer modelling platform would permit combination of traditional algorithmic steps and ML steps into larger pipelines [62]. Limited forays have been made into incorporating neural networks into Rosetta protocols (such as the Rosetta `PTMPredictionMetric`, used to predict sites of post-translational modification in natural proteins [30]). While inference for trained models can be carried out on GPU, smaller models show little benefit over CPU evaluation unless multiple inputs are provided. This is yet another argument for an architecture that operates on many `Pose` objects at a time; unfortunately, this would require complete refactoring of Rosetta.

##### Inflexible architecture, difficult development workflow, and steep learning curve

The introduction of Rosetta 3 in 2009 made certain development tasks easier. With more modular, object-oriented architecture, Rosetta 3 permitted implementation of new `Mover` and `Filter` subclasses reasonably easily [1], and this led to an explosion of both types of module [2]. Unfortunately, Rosetta was in part the victim of its own success: because `Mover` and `Filter` modules could be implemented easily, many with subtly-different, overlapping functionalities were implemented, making it difficult for users to discern which was most appropriate for any given task. The proliferation of Rosetta protocols, all of which were contributed back to a common codebase, led Rosetta 3 to swell to enormous size: at the time of this writing, the source code is 3,295,072 lines of C++ (and the unit test suite is another 268,913). The size of the codebase has further contributed to the problems of code duplication (since it is harder to determine whether the functionality that one aims to implement is already present somewhere), and has increased the learning curve (since understanding the organization of so much code is very difficult for new developers).

While `Mover` and `Filter` modules in Rosetta’s `protocols` sub-libraries proliferated, the complexity of the codebase hindered other tasks. In particular, energy function development has languished. The energy function is defined at a lower level, in Rosetta’s `core` sub-libraries, and the fear of introducing bugs affecting everyone else’s code serves to deter new developers from attempting to modify these layers. Additionally, the addition of new energy terms requires intimate understanding of the scoring system, correct sub-classing of one of many types of `EnergyMethod`, and knowledge of the various places in the Rosetta `core` sub-library in which one must register new energy methods. These requirements have also discouraged many novice developers. The consequence is that there have been no new energy terms added to Rosetta in the last two years, and only a handful (all of which are specialized) added in the last five years. Rosetta’s default energy function, `ref2015` [20, 39], has remained largely unchanged (but for some bug-fixes) since it was introduced ten years ago.

Rosetta’s architecture has also slowed scientific development in other ways. The question of how best to solve the NP-hard problems of exploring heteropolymer conformational and sequence space is an important one, but unfortunately, there is little experimentation with alternative approaches to the solvers found in Rosetta. Rosetta’s optimizers are quite powerful, but with the advent of multi-core CPUs and of GPUs for scientific computing, it makes sense to revisit the algorithms that were settled upon decades ago. Our recent demonstration that quantum annealers are useful for peptide sequence design [72] and that they can be used for *N*-body docking [73] indicates that new approaches can be fruitful, but there needs to exist a software platform with sufficiently modular architecture that new algorithms can be swapped out for older ones on demand. The recent implementation of a Covariance Matrix Adaptation Evolution Strategy optimizer (CMA-ES) as an alterative to Rosetta’s L-BFGS gradient descent optimizer (the Rosetta `minimizer`) was an exciting development [74], but unfortunately, some of Rosetta’s most commonly-used modules, such as the `MinMover`, `FastRelax`, and `FastDesign` are still hard-coded to use the Rosetta `minimizer`, and provide no option for using CMA-ES as an alternative. An ideal platform would provide easy means of integrating new modules into existing pipelines, something that Rosetta currently fails to do.

##### Lack of an application programming interface (API)

The 2009 rewrite of Rosetta in modern C++ created Rosetta 3 library organization that was in part intended to permit the use of Rosetta, not just as standalone software, but as a set of support libraries for other software [1]. Unfortunately, this goal never materialized, in large part because there was no easy way to define an application programming interface (API) for any of Rosetta’s modules or libraries. An API is a set of functions or methods by which one piece of software may interface with another. A strong API definition provides guarantees to external developers that particular interface functions will retain their function signatures from revision to revision of the software, allowing other software to be written to access these particular functions. It also serves to minimally restrict internal developers, who are free to refactor any code as needed that is not part of the external interface. Because Rosetta lacks an API, any change to any public function in any class throughout the codebase potentially breaks the ability for other software to link Rosetta – that is, all three million lines of Rosetta’s code represent potential external interface for want of a defined API. This is an intractably large interface to keep stable.

As mentioned in section 2.1.1, the introduction of PyRosetta, which provides Python bindings for almost all public functions in the C++ codebase [67], compounded this problem. Given the lower barrier to entry to Python programming, this led to an explosion in development, with thousands of user scripts accessing countless pieces of the internal workings of Rosetta. Unfortunately, the lack of a defined API has meant that any change to the Rosetta C++ code — even the simplest bug-fix that changes an internal function signature — could break any number of user scripts.

Interestingly, the RosettaScripts scripting language [68] provides the closest thing to an API definition, insofar as it provides an XML schema definition (XSD). Although this is a description of a *user* interface (the allowed XML commands that can be used to configure Rosetta modules) rather than a *programming* interface, it resembles an API definition in that the RosettaScripts XSD contains a clearly defined list of the allowed commands, types accepted, and syntax. This has served as inspiration for the Masala API definitions described in section 2.2.3.

##### Closed-source codebase

Rosetta is not an open-source project. Code added to the Rosetta repository becomes the intellectual property of the Rosetta Commons (the consortium that owns and manages the Rosetta software), and, under Rosetta’s proprietary software licence, may not be incorporated into other software projects. Although Rosetta’s source code is made available to users, and may be modified for personal use, modified versions of the software cannot be redistributed. Additionally, Rosetta is only free for noncommercial users: for-profit companies must pay a licensing fee to access the software, further limiting access to new scientific methods implemented in the Rosetta codebase.

In addition to the obstacle that Rosetta’s closed-source codebase creates for ideas to spread to other software, Rosetta’s proprietary software licence creates obstacles to absorbing good ideas from other software. The Rosetta licence is incompatible with many commonly-used strong copyleft licences, such as the GNU Public Licence (GPL), which prevents the incorporation of many pieces of open-source scientific code into Rosetta. This has particularly hindered the cross-pollination of ideas and techniques between Rosetta and other similar molecular modelling software, such as the open-source GROMACS [75] and OpenMM [76] packages used by the molecular dynamics (MD) community. Advancements in MD force fields are rarely benchmarked for Rosetta-style molecular modelling, and enhancements to Rosetta’s energy function are rarely tested in molecular dynamics simulations, largely due to licensing obstacles that prevent incorporation of scoring code from Rosetta into MD software or force field code from MD software into Rosetta.

Additionally, the lack of an open-source licence deters code contributions for some. Graduate students and postdoctoral scholars who leave Rosetta labs for industry lose access to their own code unless they are willing to pay a licensing fee, and industry scientists who are authorized to contribute code to open-source projects under standard licences can have much more trouble convincing their companies’ legal teams to approve contributions to Rosetta given the non-standard and closed-source Rosetta licence. The Rosetta Commons has been discussing options to open the Rosetta source code, but at the time of this writing, Rosetta remains closed-source.

### 2.2 The Masala software suite

We have introduced the Masala software suite as an eventual successor to Rosetta. Because Rosetta’s many strengths (see section 2.1.1) have produced a large user base, and because Rosetta is particularly necessary for cyclic peptide design pipelines that cannot plausibly be replaced with ML-based pipelines, it has been important to devise a strategy for developing Masala while still permitting new Masala features to be used in existing Rosetta pipelines. Since this goal was in close alignment with the goal of producing software that would have a stable API that would permit it to be used, not only as a standalone package, but as a set of support libraries for other software, we first prioritized development of the API system (section 2.2.3) so that, as we gradually added features to Masala intended to update or replace elements of Rosetta’s functionality, we could continue to use those pieces in Rosetta cyclic peptide design and validation pipelines, or in other contexts in which Rosetta is the current state of the art. This “Ship of Theseus” approach is intended to minimize disruption to Rosetta users while simultaneously avoiding bifurcation of cyclic peptide design and validation pipelines, as might happen if some new features were being added to Rosetta while others were being added to Rosetta’s eventual replacement.

This section provides an overview of Masala’s features that should be useful to both users and developers, while Appendices B and C (sections 8 and 9) provide a more detailed, developer-oriented tour of the Masala codebase. Section 2.2.1 introduces the Masala plugin system. When another software package has been adapted once to use Masala plugins, novice developers can add new features to the other software simply by implementing them in Masala plugin libraries that can be linked by the other software at runtime, without ever having to alter the other software again. Section 2.2.2 describes the Masala concepts of *engines* and *data representations*, used for making hard problems efficient to solve on a given type of hardware. section 2.2.3 describes the Masala build system, and the means by which Masala facilitates the definition of a stable

API, even by relatively inexperienced developers. Since a stable API depends on being able not only to add functionality, but to deprecate old functionality, section 2.2.4 describes the simple process for deprecating API functions. And finally, section 2.2.5 describes the open-source Masala licence. Additional details about Masala coding conventions for developers are provided in Appendix A (section 7).

#### 2.2.1 The Masala plugin system

Masala’s Core library, described in detail in Appendix B (section 8), provides full functionality for loading other Masala libraries at runtime, and for making the modules defined in these libraries available to other code linking Masala. We refer to runtime-loaded Masala libraries as Masala *plugins* or *plugin libraries*, and the classes defined within as Masala *plugin modules*.

In order for a plugin system to be functional, three elements are needed. First, there must be a central means of keeping track of what plugin modules are available, and for permitting code to request modules by name. The `MasalaPluginLibraryManager` and `MasalaPluginModuleManager`, both defined in the `base` sub-library of the Masala Core library (discussed further in section 8.1.5 in Appendix B), provide this functionality, with the former keeping track of what Masala plugin libraries have been loaded, and the latter maintaining a database of the individual modules within each library and the categories in which they fall. Second, there must be a standard means by which plugin modules register themselves with the central registry. To this end, all Masala plugins must define a C-style function, `extern "C" void register_library()`. This function is called by the `MasalaPluginLibraryManager` when a new plugin library is first loaded. And third, once a piece of calling code interrogates the list of available modules and requests a particular module, there must be a means by which the code can determine the interface for the module so that it can be used (since this was not available at compilation time). This is provided by the automatic API definition system described in section 2.2.3.

**Fig. 1** shows the operation of the Masala plugin system. At compilation time, other software need only be linked against Masala’s Core library, which defines the `MasalaPluginLibraryManager` and `MasalaPlug-inModuleManager` (**Fig. 1A**). At runtime, other software can submit one or more Masala plugin library paths (possibly provided by the user, loaded from a configuration file, or read from the system environment) to the `MasalaPluginLibraryManager` (**Fig. 1B**). The library manager checks the path and runtime links any Masala plugin libraries found. It then calls the `register_library()` function defined in each plugin library. This function is expected to do two things. First, it registers the plugin library with the `MasalaPlug-inLibraryManager`, providing details about version and requirements. (Version compatibility checks are automatically performed by the `MasalaPluginLibraryManager` with the assistance of another module, the `MasalaVersionManager`, described in section 8.1.10 in Appendix B.) Second, it calls registration functions for each sub-library, which register each module that has a defined API with the `MasalaPluginModuleM-anager`. These registration functions provide an instance of a creator class for each plugin module to the `MasalaPluginModuleManager`. Since the generation of these functions is fully automated and handled by the build system, developer error is minimized, and most developers need not concern themselves with the mechanics of this process.

**Fig 1.**
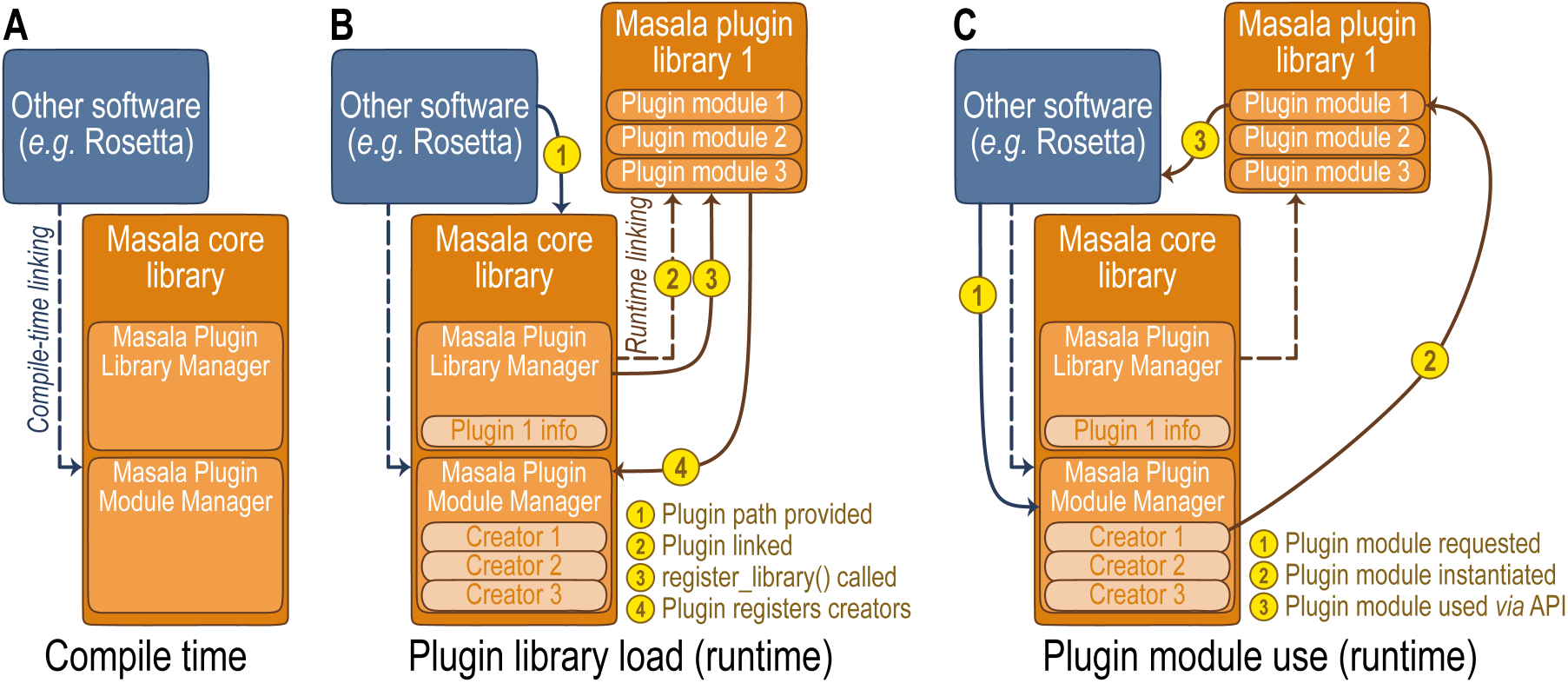
Overview schematic of Masala’s plugin system. Masala modules are shown in orange, and external code in blue. (**A**) External code may be linked at compilation time (dashed blue line) against the Masala Core library, which defines the `MasalaPluginLibraryManager` and the `MasalaPluginModuleManager`. (**B**) At runtime, external code may provide paths to Masala plugin libraries (possibly based on user inputs) to the `MasalaPluginLibraryManager` (**1**). The `MasalaPluginLibraryManager` searches each provided path for dynamic link libraries (`.dll`, `.so`, or `.dylib` files, depending on one’s operating system), dynamically links each (brown dashed line, **2**), and attempts to call `register_library()` in each (**3**). The plugin library’s `register_library()` function registers each of the library’s API-accessible classes with the `MasalaPluginModuleManager`, passing to it an instance of a creator for each such class (**4**). Creators provide information to the `MasalaPluginModuleManager` specifying the name and categories for each plugin module. (**C**) Once plugins have been registered with the plugin managers in Masala’s Core library, external code may interrogate the `MasalaPluginModuleManager` for lists of available plugin modules, and may request these by name (**1**), possibly based on user inputs. The `MasalaPluginModuleManager` invokes the creator for the requested plugin module to instantiate it (**2**), and provides it to the calling code, which may then use the module through the API definition described in section 2.2.3 (**3**).

Once the plugin library has been registered, any code that was linked against Masala’s Core library at compilation time may submit requests to the `MasalaPluginModuleManager` for lists of available plugins in hierarchical categories (expressed as vectors of strings, **Fig. 1C**). Calling code may then request a particular plugin module by name (string), which triggers instantiation through the module’s creator. Calling code may then use that module by calling its API functions through its API description, as described in section 2.2.3. Note that while queries for plugin modules and the process of instantiating requested modules may be somewhat slow (on the order of microseconds), due to the need to generate and parse strings, once a plugin class is instantiated, it can be used efficiently through its API definitions with native input and output of data, with no string parsing.

#### 2.2.2 Engines and data representations

While speed is important in macromolecular modelling software, it is not necessary to give equal priority to making all parts of the code efficient. Typical heteropolymer design and validation pipelines involve a series of steps, only some of which involve sub-problems that are NP-hard, generally poorly-scaling, or otherwise computationally expensive to solve. These steps are the ones which consume most of the computing time and which need to be the most efficient. Rosetta strove to make these steps efficient by maintaining a general-purpose data structure, called the `Pose`, that was intended to present any required structural data in the format needed by modules that would operate on it. Sub-problems in heteropolymer modelling, such as sampling chain conformations and updating atom coordinates as internal degrees of freedom change, carrying out combinatorial optimization to predict sidechain conformations or to design sequences, or relaxing structures to nearby minima in the conformational free energy landscape were handled by dedicated Rosetta modules (in these examples, the kinematic machinery built into the `Conformation` object contained within a `Pose`, the Rosetta simulated annealing based `packer`, and the Rosetta gradient-descent `minimizer`, respectively), each of which was hard-coded to use a particular algorithm, and each of which expected data to be represented in a convenient form for that algorithm. While this did make these operations efficient, it was also an obstacle to experimentation with new approaches. This also gave rise to disputes about the single “best” means of representing molecular data, such as the extended debate at a Rosetta developers’ meeting in 2013 over whether Rosetta’s `Residue` class, which stores geometry for an instance of a particular amino acid residue, should internally represent its data as a graph or as a vector. The reality was that both sides of the debate were correct: under different circumstances or for different tasks, each representation could be the better choice.

Masala uses a different paradigm. The Masala `MolecularSystem` (section 8.3.2 in Appendix B), used to represent structures, is intended for generality over speed. The subset of steps in a protoocol that are most expensive are handled by *engines*. Each engine solves a particular problem, and operates on one or more *data representations*. A protocol must take the general data structures used to move information from step to step and convert the relevant data within to a suitable data representation for the selected engine. The engine then operates on the data representation, producing an output that is converted back to the general representation used to move on to the next step. A concrete example of this is the rotamer optimization problem described in section 10.3 in Appendix D. Since this maps to a type of problem called a cost function network optimization (CFN) problem, a protocol may request a Masala CFN optimization engine, prepare a suitable CFN data representation from the all-atom molecular geometry data, pass this to the optimization engine, and receive back a CFN solution which it can interpret as a rotamer assignment. Since engines and data representations have common external interfaces but are “black boxes” on the inside, with their own algorithms and means of organizing information, respectively, the engine/data representation paradigm means that molecular modelling modules like Rosetta `Mover` subclasses may be written to request a user-defined engine of a given type, and may prepare a compatible data representation and run the engine without having to be written for a *particular* engine or data representation. This means that, should someone implement, say, some clever new means of solving CFN problems, a new CFN solver engine and CFN data representation could be implemented in a new Masala plugin library, loaded at runtime, and used for rotamer optimization without having to modify the code that handles the molecular geometry or sets up the problem for the new solver. The cost of performing data conversion once at the start or end of an expensive step is dwarfed by the potential time savings from efficiently performing the step itself.

#### 2.2.3 The build system and automatic API sub-library generation in Masala

One of the challenges in developing modular, extensible scientific software is ensuring that modules may be linked and called from other software, and that their application programming interface (API) remains stable from version to version, or at least follows a well-controlled process for the modification and deprecation of interface elements. Masala solves this by separating hand-written sub-libraries, which are considered to be mutable from version to version, from auto-generated API sub-libraries, which are intended to be stable and safe for external code to access. A given Masala plugin library will contain both hand-written sub-libraries and auto-generated API sub-libraries.

Masala classes that are intended to be invoked *only* by other modules within a Masala library may have arbitrarily-defined public member functions, since changes to these will result in compilation errors when the library is built if all calling code within the library is not updated as well. Masala classes intended to be accessible to other libraries, however, must define their own public-facing application programming interface (API) by implementing a public `get_api_definition()` function override, returning a constant weak pointer to a `MasalaObjectAPIDefinition` object. The `MasalaObjectAPIDefinition` object provides a full description of all getters, setters, and work functions that should be exposed to outside code. Contained within this object are `MasalaObjectAPISetterDefinition`, `MasalaObjectAPIGetterDefinition`, and `MasalaObjectAPIWorkFunctionDefinition` objects, each of which in turn contains a function name, its description, descriptions of all inputs and outputs in the function signature, and a pointer to the function for the current instance of the class. (The `MasalaObjectAPIDefinition` is returned by weak pointer instead of by shared pointer since the instance of the class providing it retains ownership of the `MasalaObjectAPIDefinition` object, and on destruction will ensure that it too is destroyed; calling code must therefore check the existence of the object by first obtaining a shared pointer to the `MasalaObjectAPIDefinition` prior to directly using the function pointers.) Within each sub-library, objects that define `get_api_definition()` functions must also be registered in an `api/generate_api_classes.cc` file needed by the build system.

At compilation time, the CMake build script first builds the hand-written sub-library. It then runs Masala code that it has just built, which calls the `get_api_definition()` function for each class that implements this function, and uses the API definitions to generate a JSON description of the API for that sub-library. This in turn is used by a Python script, `generate_library_api.py`, which is part of the Masala build system, to auto-generate C++ code for a new API sub-library, containing one API container class for each class in the hand-written sub-library that has an API. Each API container is designed to hold an instance of the corresponding hand-written class. The API container class has pass-through public member functions for each function in the hand-written class that is included in the API definition; these pass-through functions simply call the equivalent function of the inner, hand-written class. Each API container also has its own object mutex, and each auto-generated pass-through function in the container class locks the mutex before calling the inner, hand-written class function, providing automatic thread safety. (In performance-critical work function code in classes intended to be used only by a single thread, developers can annotate functions in the API definition to specify that no mutex locking should occur.) The sequences of events executed by Masala’s build system for the Core Masala library (described in Appendix B, section 8), and for the Standard Masala Plugins library (described in Appendix C, section 9) are shown in **Fig. 2A-B**.

**Fig 2.**
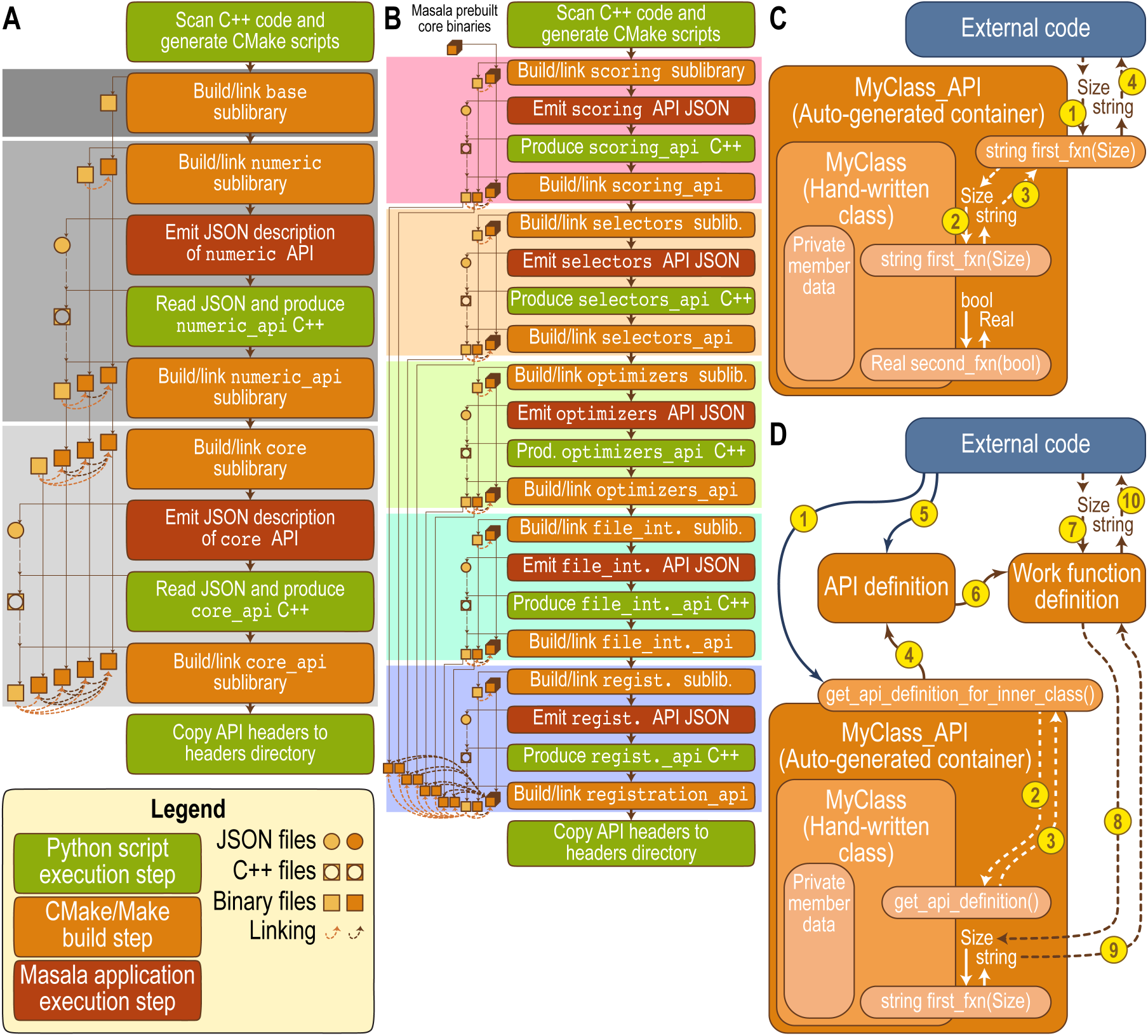
The Masala build and API generation systems. Steps executed by Python scripts are coloured green, build steps are orange, and steps involving the execution of Masala application code are red. (**A**) The sequence of automated events that occurs when building the Masala Core library. Dark, medium, and light grey regions represent the `base`, `numeric`, and `core` sub-libraries, respectively. With the exception of the `base` sub-library, each sub-library’s build process involves building the hand-written code (orange), running a Masala application to emit a JSON description of the API (red), running a Python script to generate the C++ code for the corresponding API sub-library (green), and building the API sub-library (orange). (**B**) The sequence of automated events that occurs when building the Standard Masala Plugins library. The `scoring`, `selectors`, `optimizers`, `file interpreters`, and `registration` sub-libraries are shaded in pastel red, orange, green, cyan, and blue, respectively. The same four steps occur for each sub-library. (**C**) Accessing a Masala module’s interface through the API container’s public functions, if the library containing the module has been linked at compilation time. In this case, external code can safely call public member functions of the API container (**1**), which triggers a call of the equivalent function in the inner class (**2**) with input parameters passed accordingly. The return value for the inner function is passed to the outer function (**3**) and returned (**4**). In this example, `first_fxn(·)` is part of the API, but `second_fxn(·)` is not, so the API container has no `second_fxn(·)`. By only calling API container functions, a developer is guaranteed that the interface will be stable in the future. (**D**) Accessing a Masala module’s interface through the API definition object, if the library containing the module was linked at runtime. External code must first call the API container’s `get_api_definition for inner class()` function (**1**), which calls `get_api_definition()` of the contained class (**2**). This returns an API definition weak pointer (**3**) which is returned by the outer function (**4**). External code can then convert this to a shared pointer and interrogate the API definition object (**5**) to receive API function definitions (**6**). It can then pass inputs to the function pointer contained in a given function definition (**7**) to call that function on the inner class (**8**) and receive the return value (**9**, **10**). Inner class functions that are not part of the API definition are not accessible through this scheme. Concrete examples of this are shown in sections 10.2 and 10.3 in Appendix D.

An example is shown in **Fig. 2C-D**: here, a developer has written a class called `MyClass`, with public member functions `std::string MyClass::first_fxn( Size input_int )` that is to be part of the API. Other functions may exist, such as `Real MyClass::second_fxn( bool input_bool )`, shown in **Fig. 2C**, that are *not* part of the API; here, the developer has only included `std::string MyClass::first_fxn( Size input_int )` in the API definition provided by `MyClass::get_api_definition()`. At compilation time, an API container class called `MyClass_API` is automatically generated, which contains an object of class `MyClass`. The container provides a function, `Size MyClass_API::first_fxn( Size const input int )`, which locks the `MyClass_API` mutex, then calls `MyClass::first_fxn(·)` on the contained object and returns its return value. The public member functions of `MyClass_API` therefore serve as the stable API for external code that links the plugin library at compilation time (**Fig. 2C**). Alternatively, if only Masala’s Core library is linked by the external code and Masala plugin libraries are loaded at runtime, external code may interrogate the `MyClass_API` object to receive a weak pointer to an API definition object, convert this to a shared pointer, and call iterators in the API definition object to access API function definitions that include function pointers to the public API functions of the inner class (**Fig. 2D**). Functions not part of the API definition, although publicly accessible to any other code within the plugin library in which `MyClass` is defined, are inaccessible to other Masala libraries or to codebases that link Masala. Note that class inheritance relationships for API container classes matches the class inheritance relationships for the classes that they contain. That is, if `MySecondClass` derives from `MyClass`, then the Masala build scripts ensure that `MySecondClass API` derives from `MyClass_API`.

While all of this may seem complicated, the bulk of the complexity is handled by the automated build system, minimizing developer burden. *Some* understanding of the API definition system is needed to adapt external code to use Masala plugin libraries, but this need only be done once; concrete examples of this are shown in sections 10.2 and 10.3 of Appendix D, with the API definition system’s class architecture described in section 8.1.1 of Appendix B. Novice developers need understand very little of this in order to write functional Masala plugins that provide `get_api_definition()` functions and which can be used by external code adapted to use Masala plugins: the build system fully automates the creation of the API containers.

Masala conventions specify that hand-written sub-libraries are found in a `src` directory within each plugin library repository. Auto-generated code for a given sub-library will be produced in an adjacent directory with `api` appended, in an `auto generated api` sub-directory. Within this directory, the directory structure will match the hand-written sub-library. The convention of namespaces matching directory structure will be maintained. Thus, if `MyClass` were defined in namespace `my plugin library::my sub - library::inner namespace` and located in file `src/my sub library/inner namespace/MyClass.hh`, then the `MyClass_API` container class will be auto-generated in namespace `my plugin library::my sub library -api::auto generated api::inner namespace`, and located in `src/my sub library api/auto generated -api/inner namespace/MyClass_API.hh`. In addition to the `MyClass_API` container, the build system will also auto-generate a creator class, `MyClassCreator`, in the same namespace and directory. The creator class’s sole *raison d’^etre* is instantiating `MyClass_API` objects when requested; it is the class stored by the `MasalaPluginModuleManager` and registered by the `register_library()` function. Most developers need not concern themselves directly with creator classes, since they are both created automatically by the build system and used by auto-generated registration code that is also created by the build system.

Note that the API definition system also makes it easy for code linking Masala’s Core library to auto-generate user interfaces (and even user documentation) for Masala modules contained in plugin libraries loaded at runtime. This is discussed further in Section 3.

#### 2.2.4 Function annotations and deprecation

An API definition must provide guarantees that the functions in the API will be stable from one revision of the software to the next. Nevertheless, it is inevitable that *some* changes to the API must occur over time, as the software gains functionality, bugs are fixed, or code is refactored for greater efficiency or generality. In Masala, the following three rules are expected to be followed for both Masala’s Core library and for its plugin libraries:

1. The API may grow from revision to revision. Addition of new classes or functions is always permitted.
2. The function signatures of functions that are part of the API may not change. The number and types of the input parameters, the type of the return value, and the name and constness of the function must all remain the same.
3. Functions may only be deleted by using deprecation annotations.

Note that although function signatures cannot change, it is always permissible to create a new function with a revised signature. However, the old function signature must continue to be maintained until it is deprecated.

Functions can be annotated with particular types of *function annotations*. Among these, *deprecation annotations* are notes added to function definitions that indicate that a particular function’s days are numbered. They include a library version at which the function will be deleted, and an optional library version at which warnings will be emitted if the function is called. Please see Appendix B (section 8.1.1) for an example of the code that developers may use to mark functions as being deprecated.

Masala’s build system automatically adds runtime warnings once the warning version is reached, and automatically removes the function when the deprecation version is reached. For indirect access to the inner class function by way of pass-through functions in the API container class (*i.e.* for when the plugin library is linked by external code at compilation time, as shown in **Fig. 2C**), this is accomplished by adding tracer output to the pass-through function for warnings, and by removing the pass-through function on deprecation. For direct calls of functions in the inner class by way of the function pointers provided by the API definition (*i.e.* for when the plugin library is linked by external code at runtime, as shown in **Fig. 2D**), this is accomplished by replacing the bound function with one which writes to the tracer and then calls the relevant function if the function should warn, and which throws an informative error if the function has been deprecated. The compiler variables `MASALA_DISABLE_DEPRECATION_WARNINGS` or `MASALA_ENABLE_DEPRECATED_FUNCTIONS` may be added at compilation time to disable deprecation warnings or to restore deprecated functions, respectively, though this is not recommended.

The net effect of this deprecation scheme is that developers needing to deprecate functions may annotate them as to-be-deprecated once, and then all subsequent steps of producing warnings and of actually removing the function are handled automatically by the build system. This reduces the reliance on human discipline to go though with deprecations, and frees developer time to focus on scientific code rather than on maintenance tasks like function deprecation.

#### 2.2.5 Open-source software licence

Masala and its plugin libraries are released under an AGPL 3.0 licence. This was chosen to maximize the benefit to science: the AGPL 3.0 licence is a strong copyleft licence that ensures that Masala may be used for any purpose and modified by anyone, provided that any redistribution of Masala or its derivative works occurs under the same free, open-source licence. Scientific ideas are best able to circulate and cross-pollinate if they are never made proprietary or placed under usage restrictions. This also maximally protects the interests of graduate students and postdocs who may contribute to the software, and who should have the right to continue to use their own code for their own purposes in their future careers. At the same time it minimally deters industry use, since the requirement that derivative works be *redistributed* under an AGPL 3.0 licence only applies if those works leave the hands of those individuals or organizations that develop them.

## 3 Extending existing software with Masala: Masala for Rosetta

As described in section 2.2, while Masala is ultimately intended to be usable as standalone software, it is also built to be usable as a support library for other software. Obviously, it made sense to modify Rosetta to permit Masala to extend its functionality. This section gives an brief overview of the development efforts needed to modify existing software, like Rosetta, to take advantage of Masala plugin modules, as well as a detailed explanation for users of the user experience once software like Rosetta has been modified in this way. Full details of the code for developers are in Appendix D (section 10).

Modifying existing software to make use of Masala plugin modules (including modules not known at compilation time) is somewhat more complicated than writing new Masala plugin modules, since the developer must necessarily deal with abstract base classes and with function pointers from API definitions, rather than with concrete classes with callable functions known at compilation time. Nevertheless, it is reasonably straightforward, and once this has been done once, Masala’s API definition system makes it extremely easy for even inexperienced developers to write new plugin modules that can be detected by the other software at runtime and used without any further modifications to the other software. Broadly, the steps are (1) linking the other software against Masala’s Core library, (2) modifying the other software’s user interface to permit user configuration of Masala modules unknown at compilation time, for which the user interfaces should be based on their setter API definitions, and (3) modifying the functionality of the other software to make use of work functions in Masala modules. Here, we illustrate these steps by showing how Rosetta can be linked against Masala (section 3.2), and by describing modifications that we made to Rosetta’s `PackRotamersMover`, design-centric guidance functions, and `MinMover` to allow Masala discrete cost function network (CFN) and continuous real-valued local (RVL) optimizers to be used in place of the Rosetta `packer` and `minimizer`, respectively (section 3.3, with details in Appendix D). We also provide an overview of the impact on the Rosetta user interface and user experience that comes from the added Masala support (section 3.4).

Our modifications to Rosetta are made publicly available in the `masala_rosetta_patch_public` Github repository (https://github.com/flatironinstitute/masala_rosetta_patch_public), as a patch for the Rosetta source code. This patchfile has been released under a permissive 3-clause BSD licence. Since we have set things up so that default compilation of Rosetta is unaffected, and all new C++ code is protected by `#ifdef USE_MASALA` blocks activated by compiling with the `extras=masala` compilation flag, this patch could be incorporated into public releases of Rosetta should the Rosetta community so wish.

### 3.1 Masala libraries and code organization

Masala is divided into a central Core library and any number of specialized plugin libraries. The Core library contains essential infrastructure for managing hardware resources, input and output, API definitions, and, of course, Masala plugin modules. This is intended to be the library that other software, such as Rosetta, links at compilation time, and it enables the loading of plugin libraries at runtime. Plugin libraries may define any number of Masala plugin modules, which at runtime the Masala Core library serves to software that has linked the Masala Core library at compilation time (*e.g.* Rosetta). Plugin libraries must be linked against Masala’s Core library, but can be compiled independently of Rosetta or other software linking Masala’s Core and only loaded by that software at runtime. This permits easy extension of existing software like Rosetta that links Masala’s Core, without the need to modify or recompile the existing software whenever new features are added to Masala plugin modules.

The Standard Masala Plugins library is intended to be a broadly useful plugin library for Masala users, offering CPU implementations of a range of optimization algorithms and other tools that can be used for many biomolecular modelling tasks. While any number of additional Masala plugin libraries may be introduced in the future and used alongside the Standard Masala Plugins library, we anticipate that most users will want to load this library for most tasks.

Users should be aware of this basic division between the Core library (the central infrastructure needed to enable runtime linking of Masala plugins) and Masala plugin libraries (optional components which can be loaded on demand by users who wish to take advantage of the particular plugin modules that a given plugin library may offer). Developers wishing to understand the inner workings of Masala are referred to Appendix B (section 8), which describes the Masala Core library in detail, and Appendix C (section 9), which describes the Standard Masala Plugins library in detail.

### 3.2 Linking Rosetta against Masala

At compilation time, Rosetta (or other software) need only be linked against the Masala core. To do this, first, Masala’s Core library must be compiled. Second, the system’s library path environment variables (`$LIBRARY_PATH` and `$LD LIBRARY_PATH` on Linux systems, `$LIBRARY_PATH` and `$DYLD LIBRARY_PATH` on MacOS) must be updated to include the path to the Masala Core library’s `build` directory. Third, Masala’s external headers, which are copied to the `headers` directory on compilation, must be symlinked to Rosetta’s `external/headers` directory (or otherwise placed somewhere where other software can find these headers). And fourth, the patched version of Rosetta must be compiled with `extras=masala` appended to the usual `scons` command used to compile Rosetta. The effects of `extras=masala` are in the patched version of Rosetta’s `basic.settings` file: this adds the compiler variables `USE_MASALA` and `MULTI_THREADED` (the latter triggering inclusion of certain thread safety features in the Rosetta codebase), sets the C++ language version to C++17, adds the headers directory to the include path, and adds `masala base`, `masala numeric`, `masala_-_numeric_api`, `masala_core`, `masala_core_api`, and `pthread` libraries to the linker commands. (This should be emulated to link other software against Masala’s Core library).

Once Rosetta has been compiled and linked against Masala’s Core library, Masala’s plugin libraries can be linked at runtime by passing the command-line flag `-masala_plugins <PATH1> <PATH2> <PATH3> …` to any Rosetta application. The provided paths are searched for plugins, and any encountered are loaded automatically. Section 10.1 in Appendix D shows modifications to Rosetta’s `core::init()` function to add support for loading Masala plugin libraries from user-provided paths when Rosetta initializes.

### 3.3 The developer experience: Invoking Masala modules from Rosetta code

A detailed description of the Rosetta modifications permitting Masala modules to be used may be found in Appendix D (section 10). Here, we provide a high-level overview suitable for users, to allow them to understand what the software is doing “under the hood”. Briefly, a developer must modify external software to pass a Masala plugin path to the `MasalaPluginLibraryManager`, which will load Masala plugins not known at compilation time; to this end, we modified Rosetta’s `core::init()` function as described in section 10.1 of Appendix D. The developer must also modify the external code to interrogate the `MasalaPluginModuleManager` to obtain lists of Masala plugin modules of a given type, and to use those lists to provide users with options to select the module that the user wishes to use. The developer must then ensure that the external code asks the `MasalaPluginManager` for an instance of the desired module, has the module produce its API definition, and uses the API definition to provide interface elements to the user to allow the module to be configured. Finally, the developer must ensure that the module’s work function is called, and must convert the module’s output to output usable by the external code.

As a concrete example of this, we modified Rosetta’s `PackRotamersMover`, which calls the Rosetta `packer` to perform either sequence design or side-chain conformational structure prediction, to permit a Masala CFN optimizer to be used instead, with the Masala CFN optimizers that are loaded at runtime (none of which was known at compilation time) made available to Rosetta users through the RosettaScripts interface. This involved modification of the `PackRotamersMover`’s RosettaScripts XML-parsing functions to provide an interface for these Masala modules based on their runtime-loaded API definitions, as described in Appendix D, section 10.2. We also altered the `PackRotamerMover::apply()` function to convert the rotamer optimization problem to a suitable CFN problem data representation on which the Masala optimizer can operate, to run the Masala optimizer, and to convert the CFN solution back to a rotamer selection. Section 10.3 of Appendix D describes these modifications. We also made additional modifications to Rosetta’s design-centric guidance scoring terms [42, 43] to configure suitable Masala `CostFunction` instances, to allow these also to be used seamlessly with Masala CFN optimizers. Appendix D, section 10.4 provides a detailed description of the modifications made to allow Rosetta’s `hbnet` term, which promotes hydrogen bond networks during design, to be used with Masala CFN optimizers, and section 10.5 summarizes the modifications for the `aa_composition`, `voids_penalty`, `buried unsatisfied penalty`, and `netcharge` terms.

We made similar modifications to Rosetta’s `MinMover`, permitting it to use Masala RVL optimizers in place of the Rosetta `minimizer` to relax heteropolymer structures. These modifications are described in Appendix D, section 10.7. Finally, we ensured that Rosetta’s `FastRelax` and `FastDesign` movers, which use the `PackRotamersMover` and `MinMover` to perform alternating rounds of discrete side-chain optimization and continuous relaxation, could also use Masala CFN and RVL optimizers for these tasks, respectively.

Having made these modifications, new Rosetta functionality can be added at runtime simply by providing new Masala plugins. Step-by-step instructions for writing new Masala plugin modules are found in section 10.8 in Appendix D.

### 3.4 The user experience: Controlling Masala modules from the Rosetta user interface

section 3.3 and Appendix D (section 10) describe the process by which developers can extend existing software, like Rosetta, to use Masala plugin modules, and, once this is done, how they can add new Masala modules that can be used in Rosetta protocols without altering Rosetta’s source code or recompiling Rosetta. In this section, we describe what this means for a Rosetta user (or for users of other software linking Masala). Section 3.4.1 explains how global defaults in Rosetta can be set at the command-line to specify that wherever a particular Rosetta optimization engine, such as the `packer` or the `minimizer`, would have been used, a particular Masala optimization engine should be used instead. Section 3.4.2 describes how Masala modules may be inserted selectively into particular steps of Rosetta protocols through the RosettaScripts scripting interface. Both of these interfaces automatically extend themselves as new Masala plugin modules are loaded; however, this relies on there existing some means by which a user can figure out what Masala modules are available and how to configure them. Section 3.4.3 describes how Rosetta’s online help can produce information for the user about configuring Masala plugin modules. Finally, section 3.4.4 describes how the user can configure an IDE like VSCode to allow tab-completion, syntax highlighting, and mouse-over help when editing RosettaScripts XML scripts that contain Masala elements.

#### 3.4.1 Global control from Rosetta command-line flags

Masala modules may be configured globally, to replace Rosetta modules in all instances in which the Rosetta modules would have been used, by passing command-line flags to any Rosetta application. The `-default_masala_packer_configuration <ONFIGURATION file name>` command-line flag allows the user to provide a RosettaScripts-formatted XML file configuring a `PackRotamersMover` instance, to serve as the default `PackRotamersMover` configuration in all Rosetta protocols. This provides the opportunity to specify that a Masala CFN optimization engine should be used in place of the Rosetta `packer`. Similarly, the `- default_masala_minmover_configuration <CONFIGURATION file name>` command-line flag allows the user to provide an XML file configuring a `MinMover` instance, to serve as the default `MinMover` configuration wherever relaxation is performed. This allow a Masala RVL optimization engine to be set as the default relaxation optimizer, rather than the Rosetta `minimizer`. The implementation details for this are described in sections 10.3 and 10.7 in Appendix D.

An example configuration file for the `PackRotamersMover`, for use with the `-default masala packer configuration` flag, is shown in Listing 1. Here, we specify that a `MonteCarloCostFunctionNetworkOptimizer` is to replace the Rosetta `packer`, that it is to perform Monte Carlo trajectories with as many steps as those that would be performed by the `packer`, and that it is to perform four trajectory replicates in four threads, using a `LogarithmicRepeatAnnealingSchedule` as the annealing schedule. This may be considered a current “best practices” script, suitable for many situations, though for large design tasks one may wish to increase the number of replicates or the length of the simulated annealing trajectories.

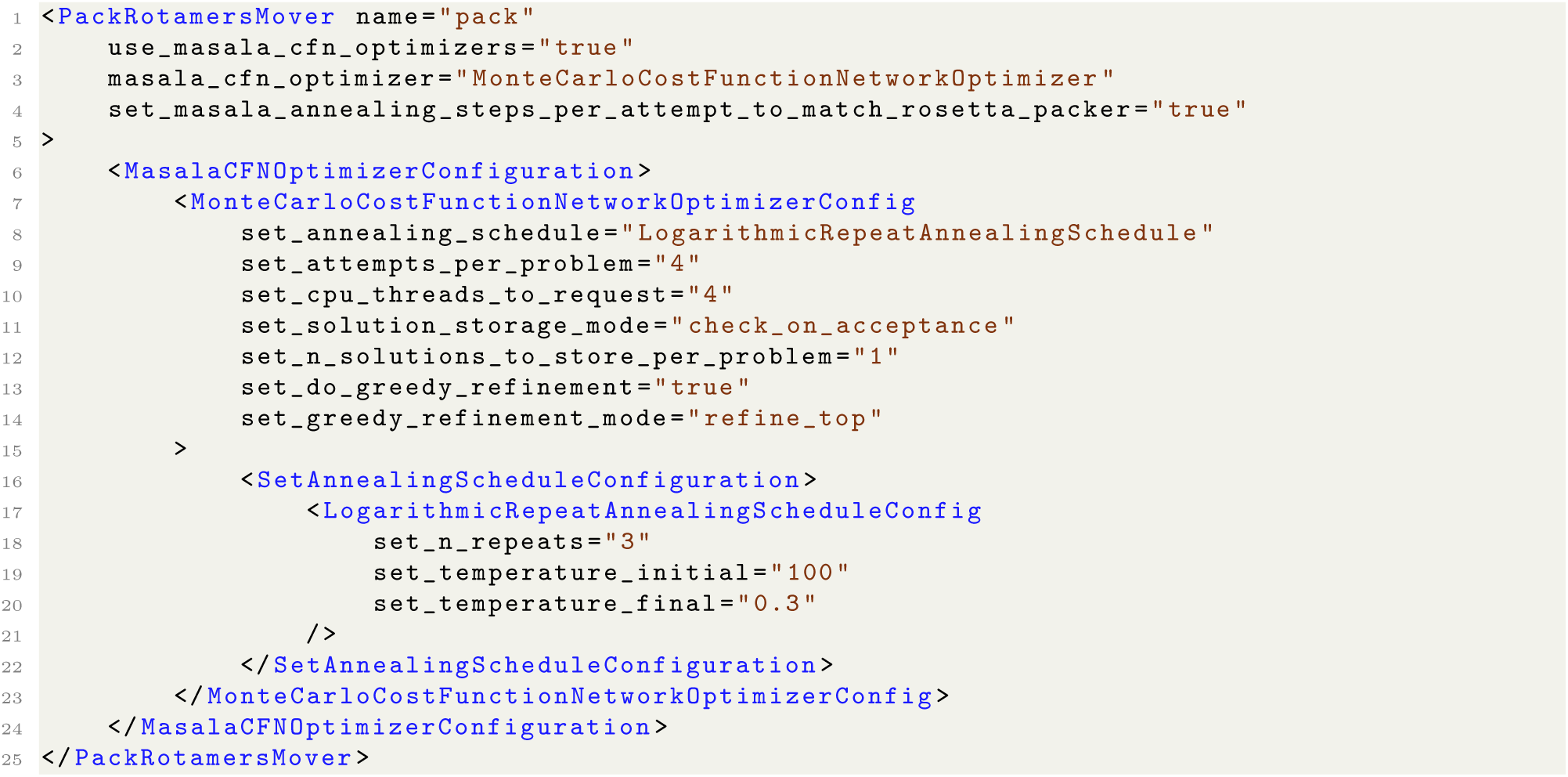

**Listing 1.** An example configuration script to set global defaults for the `PackRotamersMover`, to be passed to Rosetta with the `-default masala packer configuration` flag. This script specifies that in lieu of the `packer`, the `PackRotamersMover` should use the `MonteCarloCostFunctionNetworkOptimizer`, that the annealing schedule should be a `LogarithmicRepeatAnnealingSchedule`, that the optimizer engine should perform four trajectories in as many threads, and that each trajectory should be as many steps long as a default `packer` trajectory would be (by Rosetta convention, 100 times the total number of rotamers in the problem). This script also specifies that the top solution should be subjected to greedy refinement. As shown, this script represents current “best practices”, serving as a good starting point for rotamer optimization setup for many Rosetta protocols.

#### 3.4.2 Finely-grained control from the RosettaScripts scripting language

In addition to global control, Masala modules may be configured at particular stages of scripted protocols using the RosettaScripts language. The XML schema definition modifications and XML parsing functions described in section 10.2 of Appendix D allow automatic generation of RosettaScripts user interface elements (XML tags) for Masala plugin objects loaded at runtime, based on their API defintions.

Listing 2 shows an example RosettaScripts script that uses the `FastDesign` protocol to design a cyclic peptide, making use of Masala CFN and RVL optimizers for rotamer optimization and relaxation steps within `FastDesign`. **Fig. 3A** shows a cyclic peptide designed by this protocol. This protocol was executed using Protein Data Bank structure `5KX0`, a computationally-designed mixed-chirality peptide with right- and left-handed *α*-helices [18], as input. Note that full design protocols that sample backbone conformations and carry out design are of course possible. For demonstration purposes, since it was not our intention to illustrate backbone conformational sampling, we have provided an input cyclic peptide backbone.

**Fig 3.**
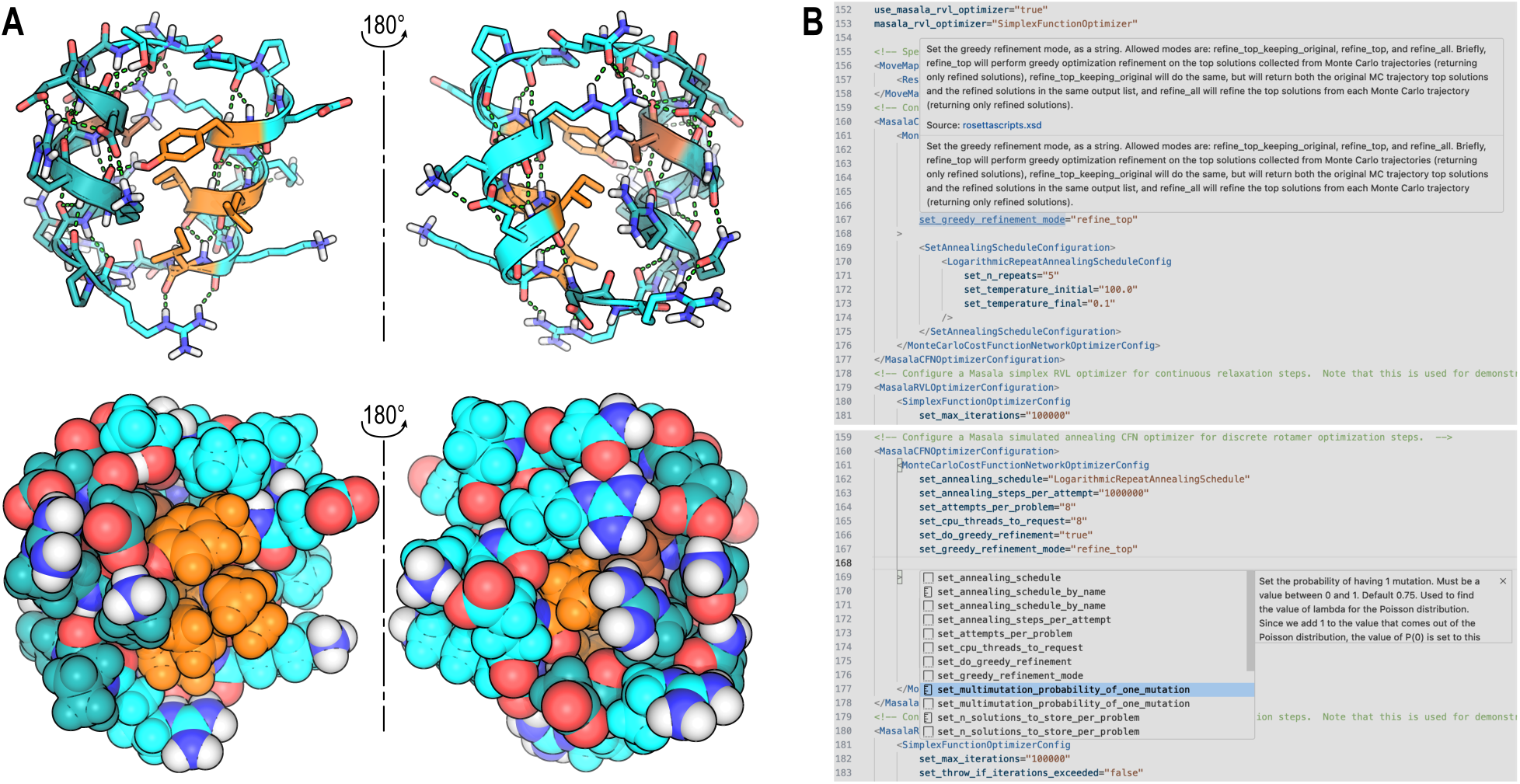
Use of Masala in a RosettaScripts context. (**A**) Example of a cyclic peptide designed with the RosettaScripts script shown in Listing 2, using the Masala `MonteCarloCostFunctionNetworkOptimizer` for rotamer optimization and the Masala `SimplexFunctionOptimizer` for relaxation in the context of the Rosetta `FastDesign` protocol. Hydrophobic `L` and `D`-amino acids are shown in orange and brown, respectively. Polar L- and D-amino acids are shown in cyan and dark teal, respectively. Top: sticks and cartoon view. Hydrogen bonds and polar contacts, detected by PyMOL, are shown as green dashed lines. Bottom: space-filling spheres view. (**B**) Use of Masala with the RosettaScripts XML schema definition (XSD) and with the VSCode IDE. When the RosettaScripts XSD is generated with `<PATH to Rosetta>/Rosetta/main/source/bin/rosetta scripts.masala.linuxgccrelease -output schema <XSD filename> -masala plugins$MASALA STANDARD PLUGINS` and loaded in VSCode (see section 3.4.4), then a user writing a RosettaScripts script enjoys mouse-over help (top), providing useful explanations of Masala modules and their options, and tab-completion (bottom), allowing available Masala modules and their options to be discovered.

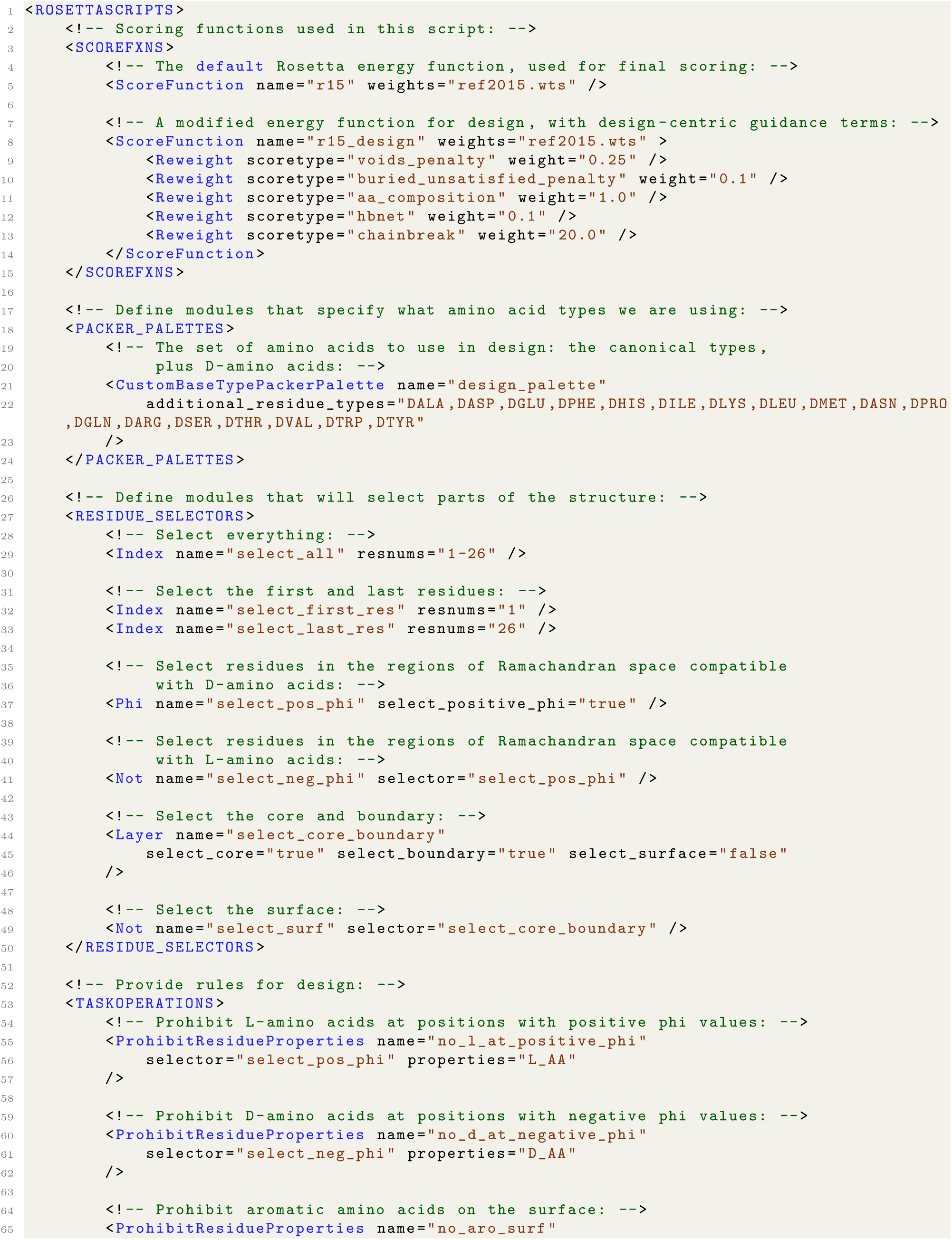
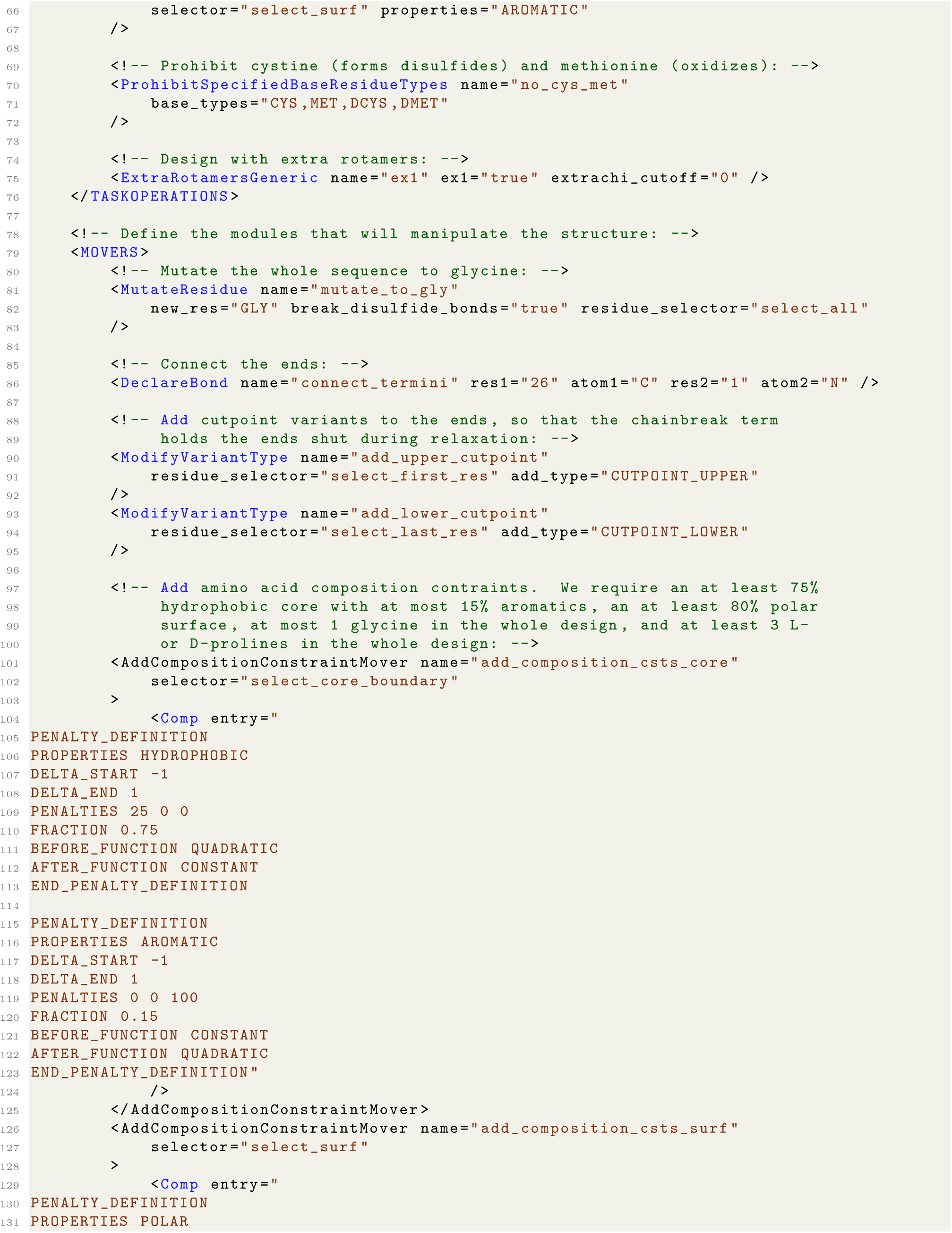
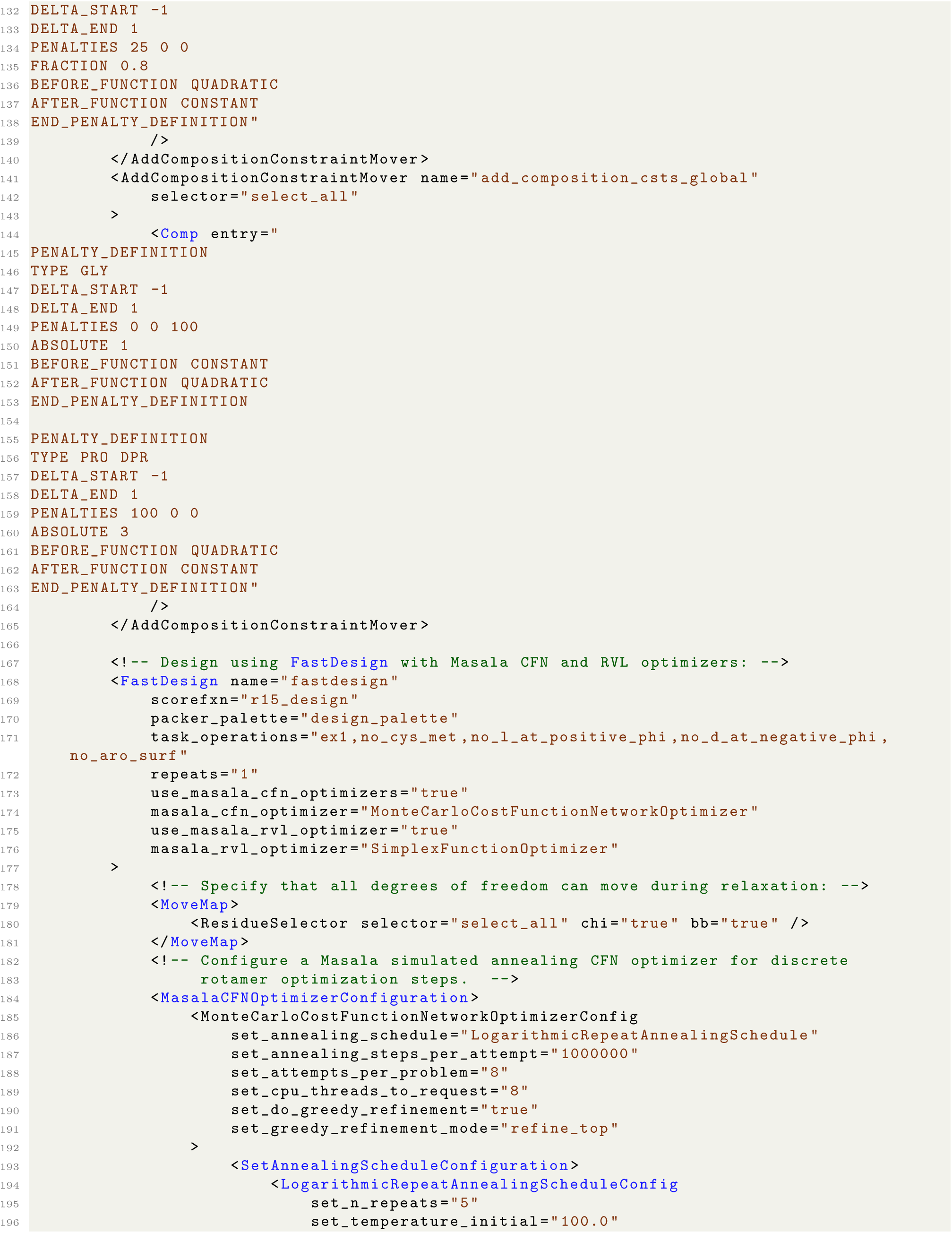
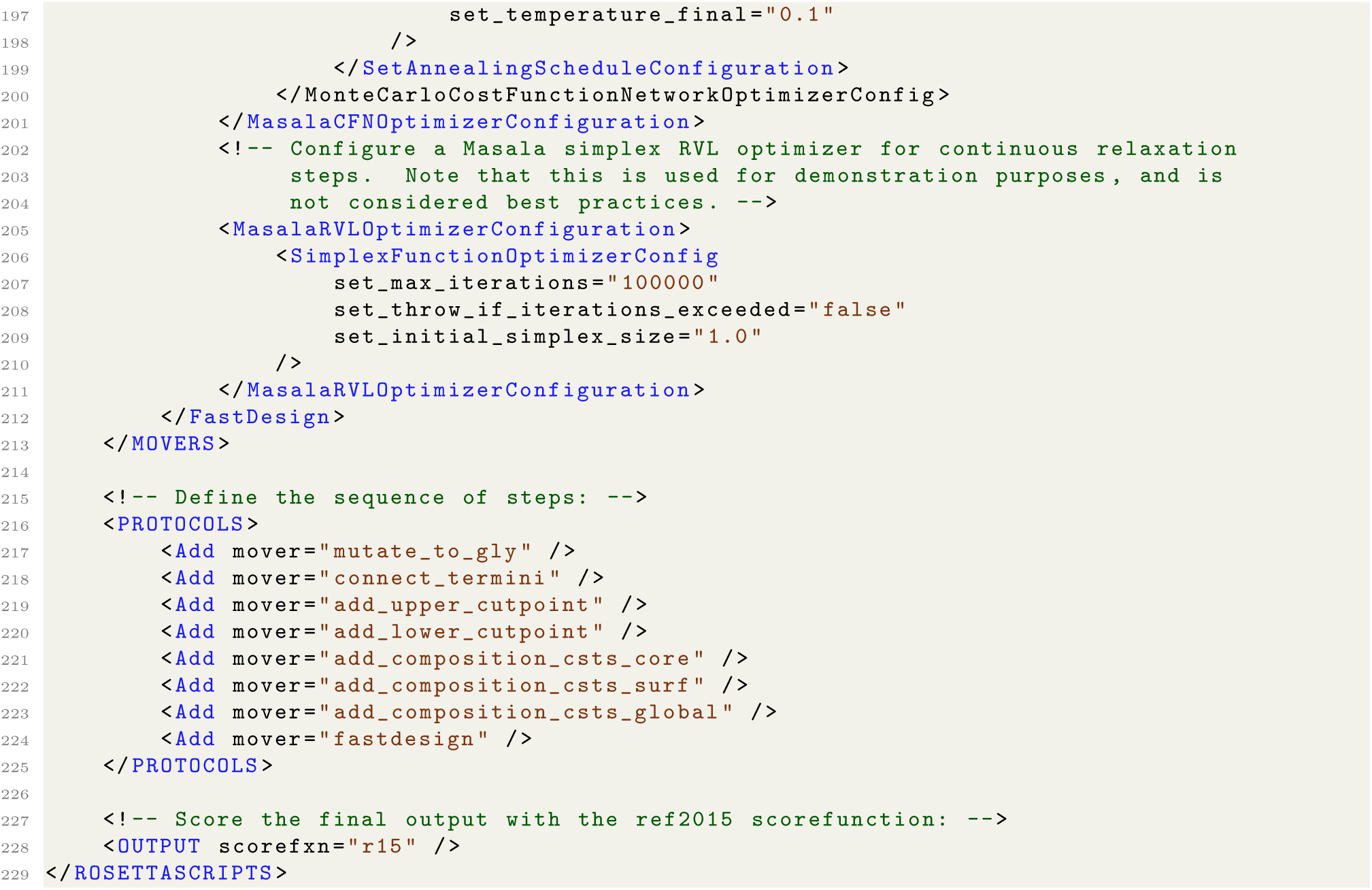

**Listing 2.** A RosettaScripts XML script that designs a 24-residue cyclic peptide, with Rosetta design-centric guidance terms `aa_composition`, `buried_unsatisfied_penalty`, `hbnet`, and `voids_penalty`, using a Masala CFN optimizer for discrete rotamer optimization and a Masala RVL optimizer for relaxation within the Rosetta `FastDesign` module. Note that this is not a “best practices” script: the use of the `SimplexFunctionOptimizer` is for illustrative purposes only. This optimizer, while useful for non-differentiable functions, is not likely to be as performant as the Rosetta gradient-descent `minimizer` or the Masala `BFGSFunctionOptimizer`.

Assuming that the input backbone file 5kx0.pdb is in an inputs/ subdirectory of the current working directory, that the design script shown in Listing 2 is `xml/design.xml`, and that the `$MASALA_STANDARD_- PLUGINS` environment variable has been set to point to the Standard Masala Plugins directory, then the design script can be executed with the command shown in Listing 3:

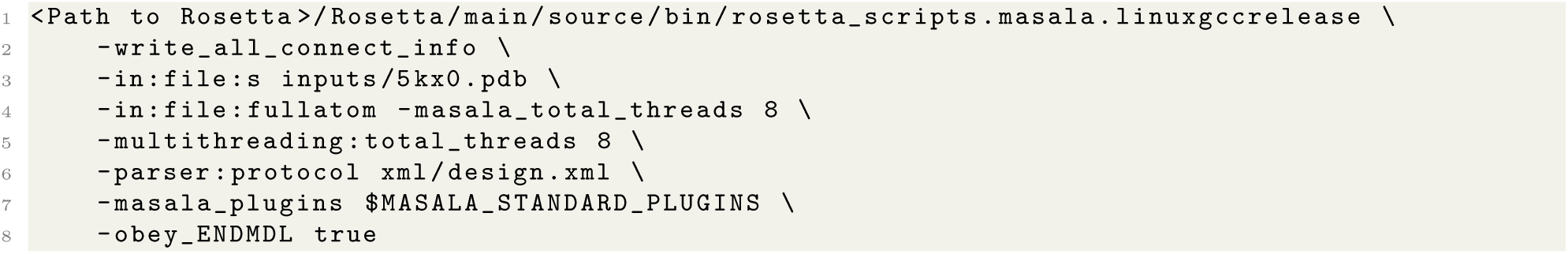

**Listing 3.** Commandline command to run the design script shown in Listing 2. For execution on other hardware or with versions of Rosetta compiled with other compilers, “linuxgccrelease” should be modified appropriately — for instance, “macosclangrelease” is typical on Macintosh systems.

Masala may also be used within a PyRosetta context by passing RosettaScripts tags to the `parse_my_- tag(·)` function of the `PackRotamersMover, MinMover, FastDesign, or FastRelax` Rosetta modules, with subtags configuring Masala plugin modules.

The existence of auto-generated syntax for configuring Masala plugin modules from RosettaScripts is of course only useful if it is discoverable for the user. Sections 3.4.3 and 3.4.4, below, describe how the user may access information about the available Masala plugin modules and the syntax for controlling these modules.

#### 3.4.3 Getting online help for Masala plugin modules

As described in section 2.1.1, the RosettaScripts application’s `-info <ROSETTA module name>` command-line flag permits users to access online help about particular Rosetta `Movers`, `Filters`, `SimpleMetrics`, or other Rosetta modules. Because the Masala API definitions permit automatic generation of human-readable descriptions of Masala plugin modules’ interfaces, when a Rosetta module that can use Masala plugin modules is interrogated with `-info <ROSETTA module name>` *and* the path to at least one Masala plugin library is passed with the `-masala_plugins <PLUGIN path>` flag, the information printed includes information about the RosettaScripts interface for the Masala plugin modules found in the provided Masala library or libraries that can be used by the Rosetta module. For example if the environment variable `$MASALASTAN-DARDPLUGINS` is set to the Standard Masala Plugins library’s path, then the command `./bin/rosetta-scripts.masala.linuxgccrelease -info FastDesign -masala_plugins$MASALA_STANDARD_PLUGINS` produces the output shown in Listing 4. Significantly, all of this user-facing documentation is produced at no additional cost to the developer, once they have written the `get_api_definition()` function for their Masala plugin module.

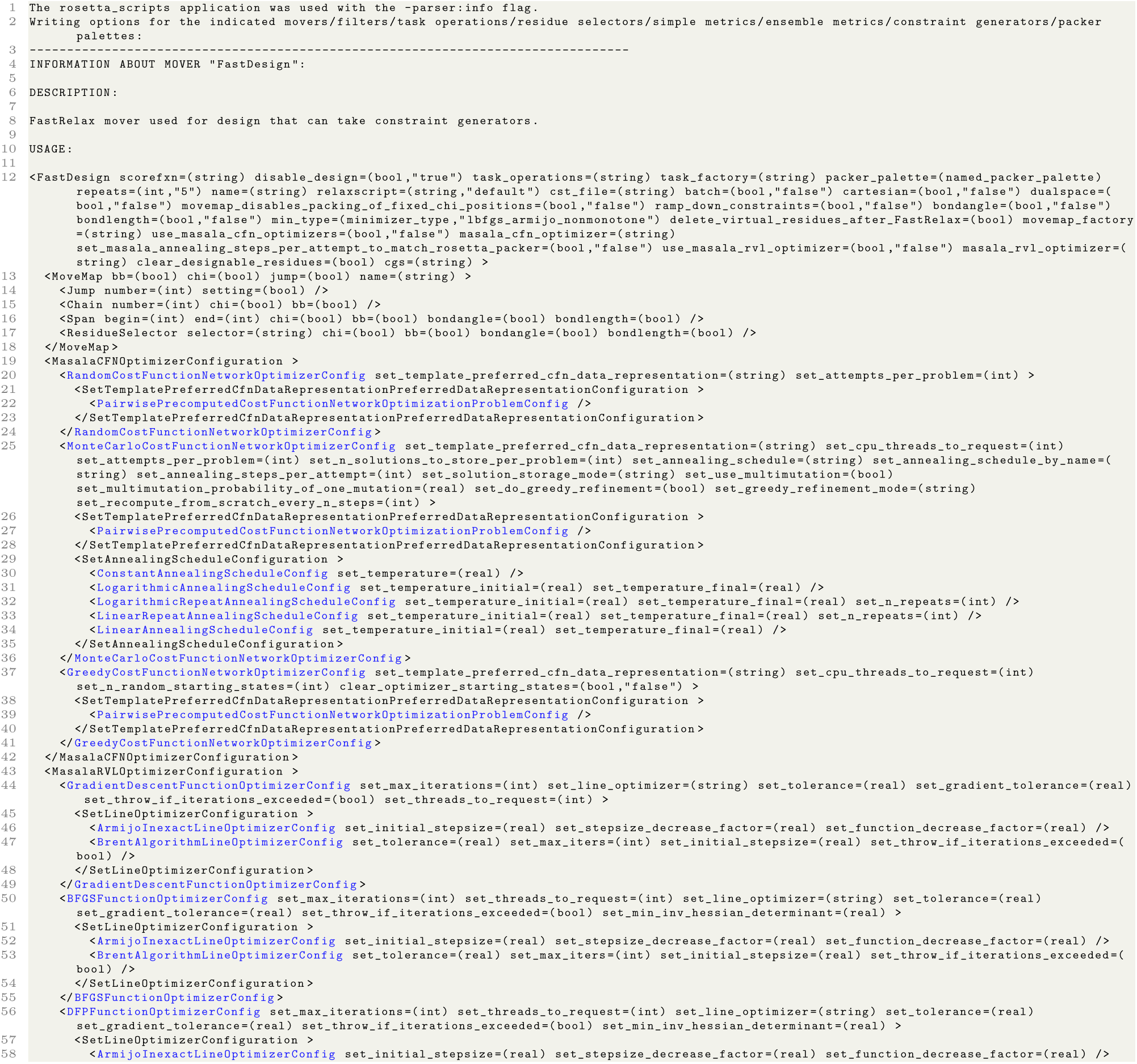
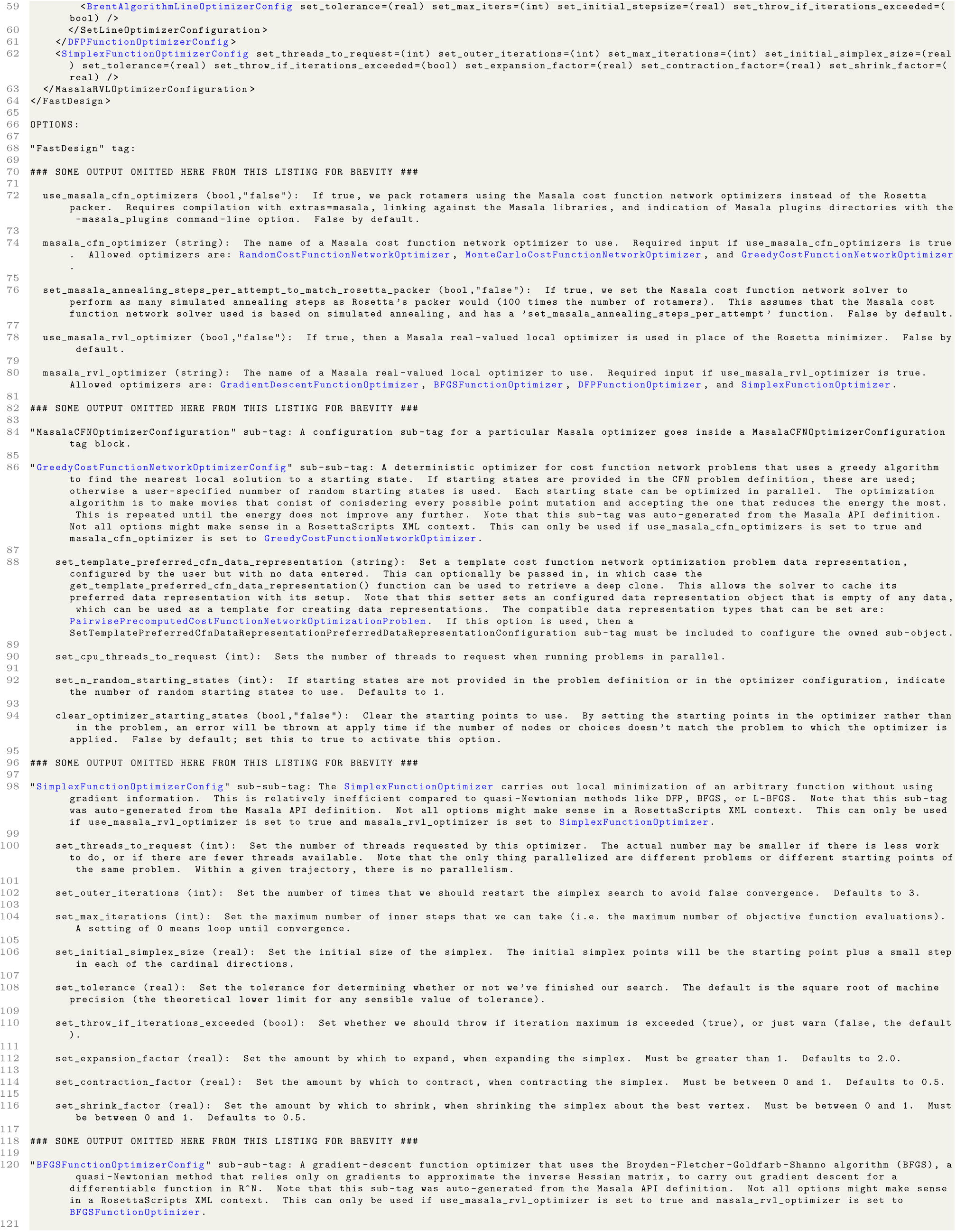
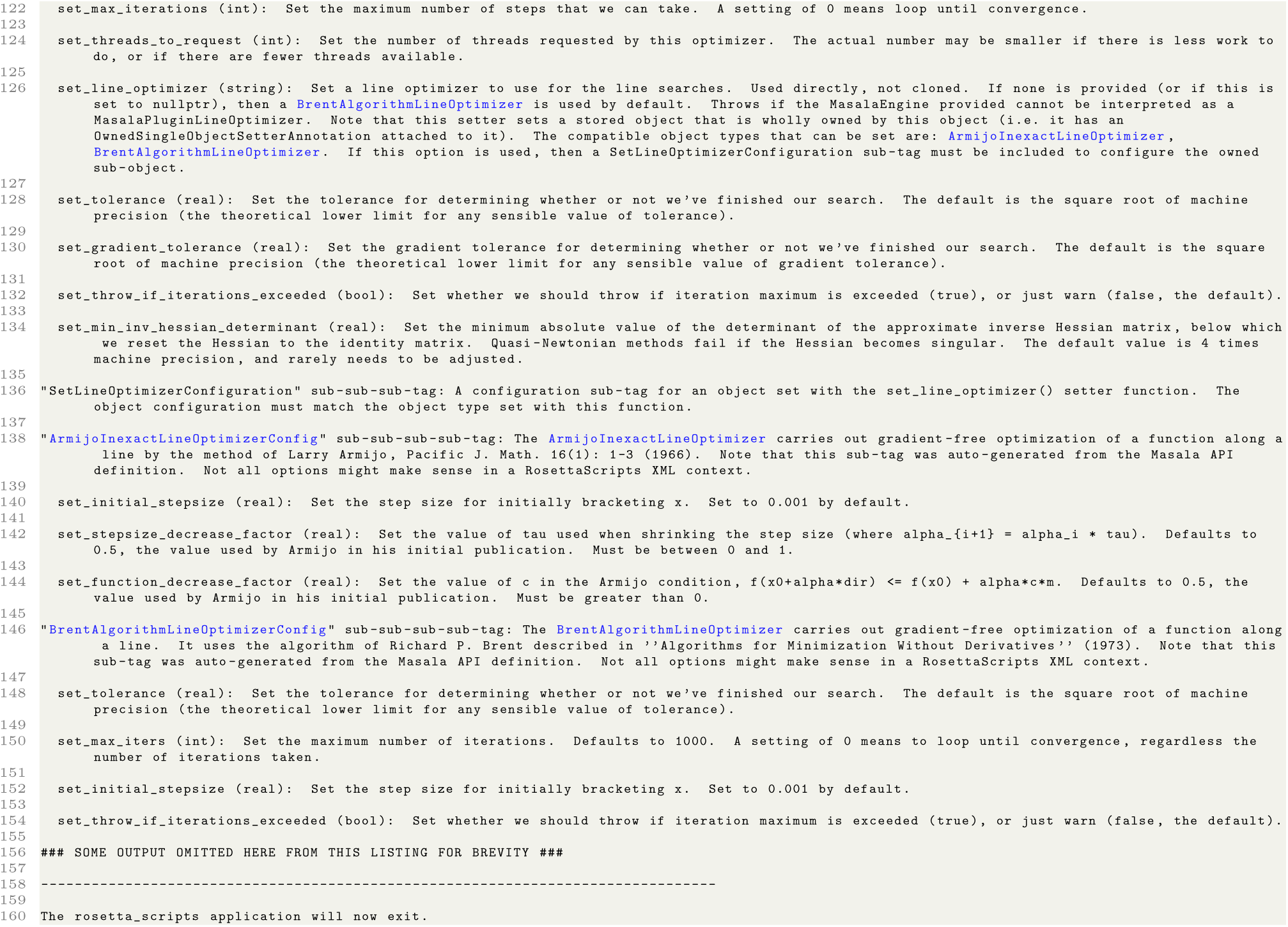

**Listing 4.** Output of running the command-line command `./bin/rosetta scripts.masala.linuxgccrelease -info FastDesign -masala plugins$MASALASTANDARDPLUGINS` to obtain information about the user interface for Rosetta’s `FastDesign Mover`. With the path to the Standard Masala Plugins, the output is augmented with information about the XML user interfaces for Masala CFN and RVL solver engines that can replace the Rosetta `packer` and `minimizer` calls. Tag names for Masala plugin modules unknown at compilation time, with user interfaces generated from API defintions at runtime, are highlighted in blue. Output is abridged; only a subset of the detailed descriptions of sub-tags are shown.

Note that in Listing 4, options that take as input a name of a Masala plugin module have descriptions that list the available Masala plugin modules that can be accepted. The `masala_cfn_optimizer` option in the `FastDesign` tag, for instance, has a description that includes, “The name of a Masala cost function network optimizer to use. Required input if use_masala_cfn optimizers is true. Allowed optimizers are: `RandomCostFunctionNetworkOptimizer`, `MonteCarloCostFunctionNetworkOptimizer`, and `GreedyCostFunctionNetworkOptimizer`.” This will update itself automatically as more CFN optimizers are added to the Standard Masala Plugins library, or if additional plugin libraries are loaded. This provides discoverability for users, pointing them to the modules that are available. Tags configuring each of these modules contain descriptions of what the modules are and what they do: for example, the `GreedyCostFunctionNetworkOptimizerConfig` sub-sub-tag has a detailed description that begins, “A deterministic optimizer for cost function network problems that uses a greedy algorithm to find the nearest local solution to a starting state.”

#### 3.4.4 Masala UI elements in integrated development environments

The RosettaScripts application’s `-output_schema <OUTPUT filename>` command can be used to write a machine-readable XML schema definition, or XSD, to an ASCII file. This is a complete description of all allowed commands and syntax for the entire RosettaScripts scripting language. When this command is used in conjunction with the `-masala plugins <PLUGIN path>` command, the XSD includes descriptions of the XML interface for any Masala plugin modules that can be configured in a RosettaScripts script. This is useful for users wishing to use integrated development environments (IDEs) like VSCode: these can generally read XSD files to enable handy features such as mouse-over help or tab completion when writing RosettaScripts XML. In order to enable this:

1. Generate an XSD file. For instance, if `$MASALA_STANDARD_PLUGINS` is an environment variable pointing to one’s Standard Masala Plugins directory, then running the command `<PATH to Rosetta source>/bin/rosetta scripts.masala.linuxgccrelease -output_schema rosettascripts.xsd masala_plugins$MASALA_STANDARD_PLUGINS` will generate a file, `rosettascripts.xsd`, containing a full description of the RosettaScripts language, including RosettaScripts commands for configuring the Masala plugin modules found in the Standard Masala Plugins.
2. Install XML support for your IDE. The `XML Complete` plugin for VSCode, for example, permits an XSD file to be read and used for tab completion and mouse-over help.
3. Tell your IDE where to find the XSD. When using the XML Complete plugin for VSCode, for example, this is accomplished by adding <root xmlns:xsi="http://www.w3.org/2001/XMLSchema-instance" xsi:noNamespaceSchemaLocation="file://<PATH to file>/rosettascripts.xsd" /> to the top of one’s RosettaScripts script. Note that this line must be commented out before *running* the script.

**Fig. 3B** shows the effect of mousing over a Masala element in an XML script when working in VSCode (top) and of pressing `ctrl + space` in the middle of a Masala element tag to bring up a list of allowed commands (bottom). This provides an easily discoverable means of working with new Masala modules, allowing users to grow familiar with their interfaces. As was true for the `-info <ROSETTA module>` flag (Section 3.4.3), the IDE support is provided with no additional effort for the developer, who need only define the `get_api_definition()` function for their Masala plugin module.

### 3.5 Summary: Masala as a support library for Rosetta and other software

This section has discussed both the developer and user experience when Masala is used as a support library for external software, using Rosetta as an example of existing software that can be extended with Masala plugin libraries. Masala’s modular architecture, API definition system, and plugin system are designed to maximize ease of use for the user and for those developing new modules of a given type (for instance, new CFN solver engines, or new RVL optimizer engines). For users, new functionality can be obtained simply by passing a path to a new plugin library at runtime, with user interfaces for new modules generated automatically and discoverable through online help or IDE interfaces, while for developers, new plugin modules can be implemented in a simple, self-contained fashion, with most of the effort of getting the new module to be usable as a plugin automated by the build system. A somewhat steeper learning curve must be navigated for those modifying existing software (such as Rosetta) to accept and use Masala plugin modules; however, because this need only be done once for all current and future species of a given genus of Masala plugin module, this cost is paid much less frequently.

The net effect is intended to be acceleration of development of scientific methods, permitting easy experimentation with new approaches, with most of a scientist’s time spent on scientific programming rather than infrastructural programming. We have already implemented support for using Masala CFN and RVL engines in Rosetta as drop-in replacements for the `packer` and `minimizer` respectively, in any Rosetta protocol. Since most Rosetta heteropolymer design and validation protocols spend the plurality, if not the majority, of their time on some combination of calls to the `packer` and to the `minimizer`, this ensures that the most computationally costly parts of design and validation protocols have now been opened up for easy experimentation, allowing development of new methods that could offer greater speed or generality.

Although the focus of this chapter has primarily been the Masala infrastructure and its potential for accelerating experimentation with new methods, our efforts have also yielded faster and more easily parallelized means of solving design problems. In the next section, we explore computational benchmarks of the single and multi-threaded performance of Masala CFN modules when used to solve real-world problems in a Rosetta context.

## 4 Computational benchmarks

In this section, we describe computational benchmarks performed on a set of realistic peptide design problems using the Masala `MonteCarloCostFunctionNetworkOptimizer`. These tests were carried out with and without the design-centric guidance functions described in Appendix D, sections 10.4 and 10.5. We make head-to-head comparisons of the `MonteCarloCostFunctionNetworkOptimizer`’s performance to that of the Rosetta `packer`. We constructed the benchmark set in the RosettaScripts scripting language [68], sampling a set of 135 conformations of a six-helix bundle and designing a 90-amino acid sequence. A RosettaScripts XML script implementing this benchmark is provided in section 4.1 (Listing 5), permitting its use to assess the performance of future Masala design tools.

We assess performance in two ways. First, we examine the *single-threaded* performance of the Masala `MonteCarloCostFunctionNetworkOptimizer`, comparing it to the performance of Rosetta’s `packer` (section 4.1). We also assess the *multi-threaded* performance of the `MonteCarloCostFunctionNetworkOptimizer`, to assess its ability to take efficient advantage of multi-core CPUs (section 4.2).

### 4.1 Single-threaded CPU performance

We carried out computational benchmarks to compare the single-threaded CPU performance of the Masala `MonteCarloCostFunctionNetworkOptimizer` with that of the Rosetta `packer`. Our benchmark was a design task: we set out to perform rotamer optimization-based sequence design on 135 conformations of a parametrically-generated 90-residue six-helix bundle, produced using Rosetta’s `BundleGridSampler` mover [19]. Our design protocol was scripted in RosettaScripts XML (Listing 5), and used the `PackRotamersMover` (which can either invoke the Rosetta `packer` or a Masala CFN optimizer, as described in section 10.3 of Appendix D) for the design step. We performed design with Rosetta’s `ref2015` energy function [20] as the objective function, and also with `ref2015` plus one of five design-centric guidance functions [42, 43]: `hbnet` (described in section 10.4 of Appendix D), which promotes hydrogen bond networks; `voids_penalty`, which discourages empty spaces in heteropolymer cores; `buried_unsatisfied_penalty`, which discourages buried hydrogen bond donors and acceptors that are not involved in hydrogen bonds; `aa_composition`, which promotes a desired amino acid composition by penalizing deviation from it; and `netcharge`, which encourages a desired net charge by penalizing deviation from it. In addition to `ref2015` plus each of these terms individually, we also benchmarked `ref2015` plus all of these terms, emulating a real-world heteropolymer design problem.

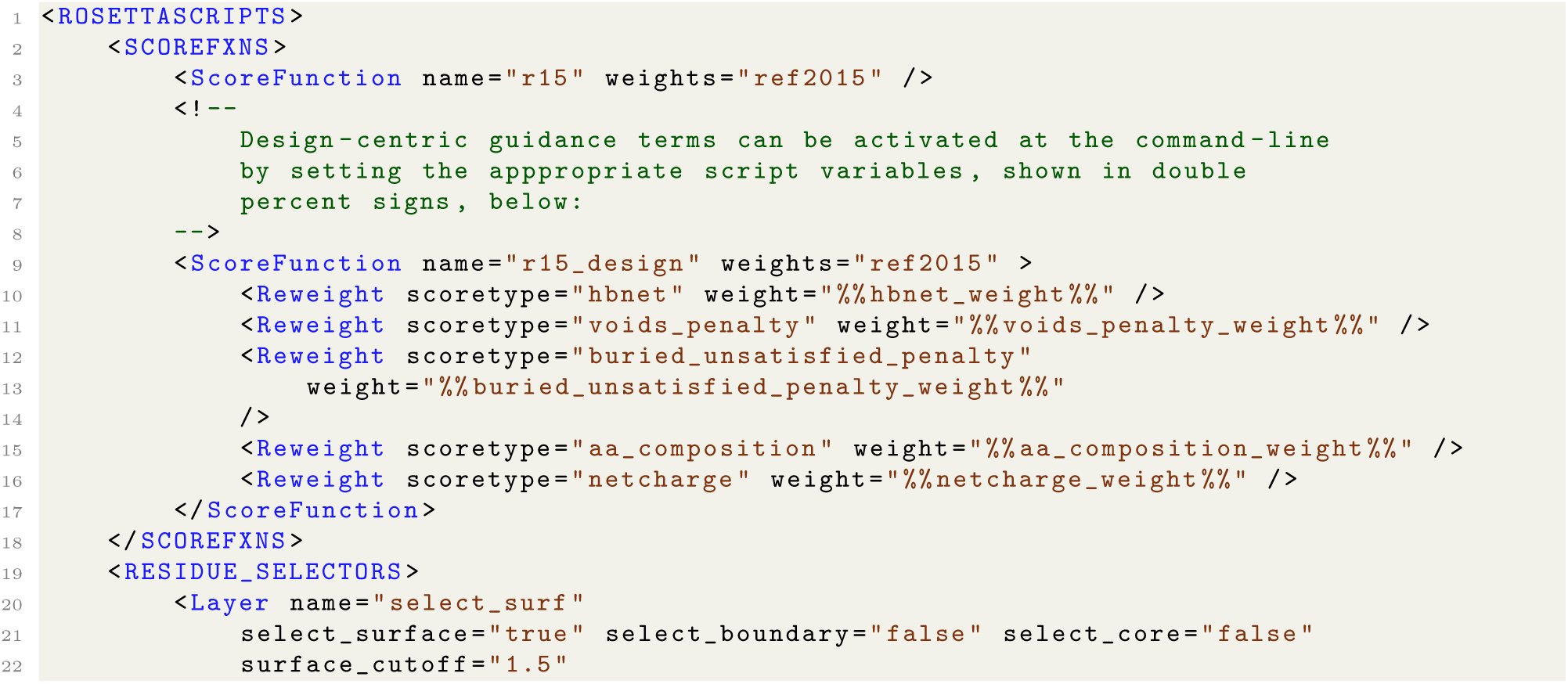
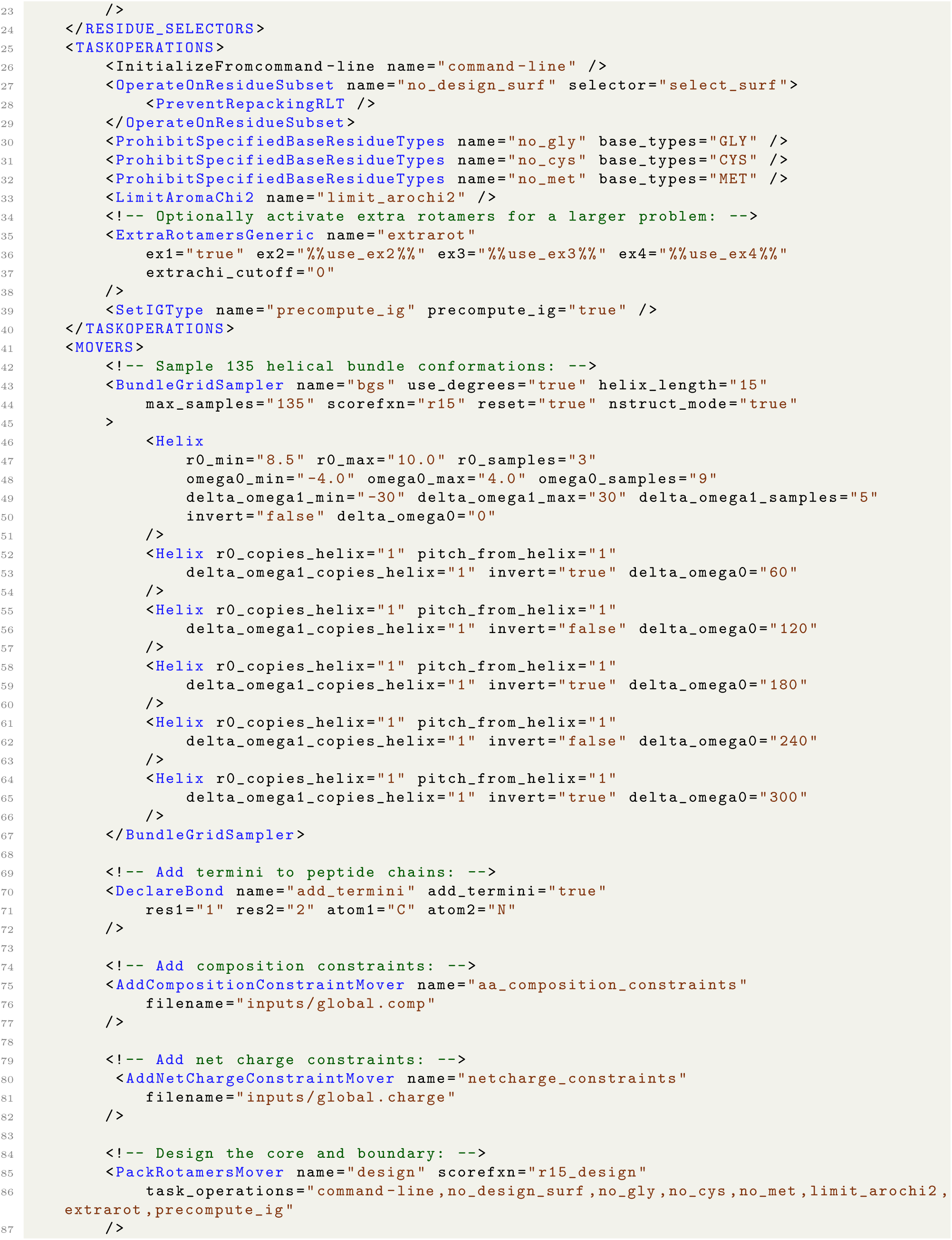
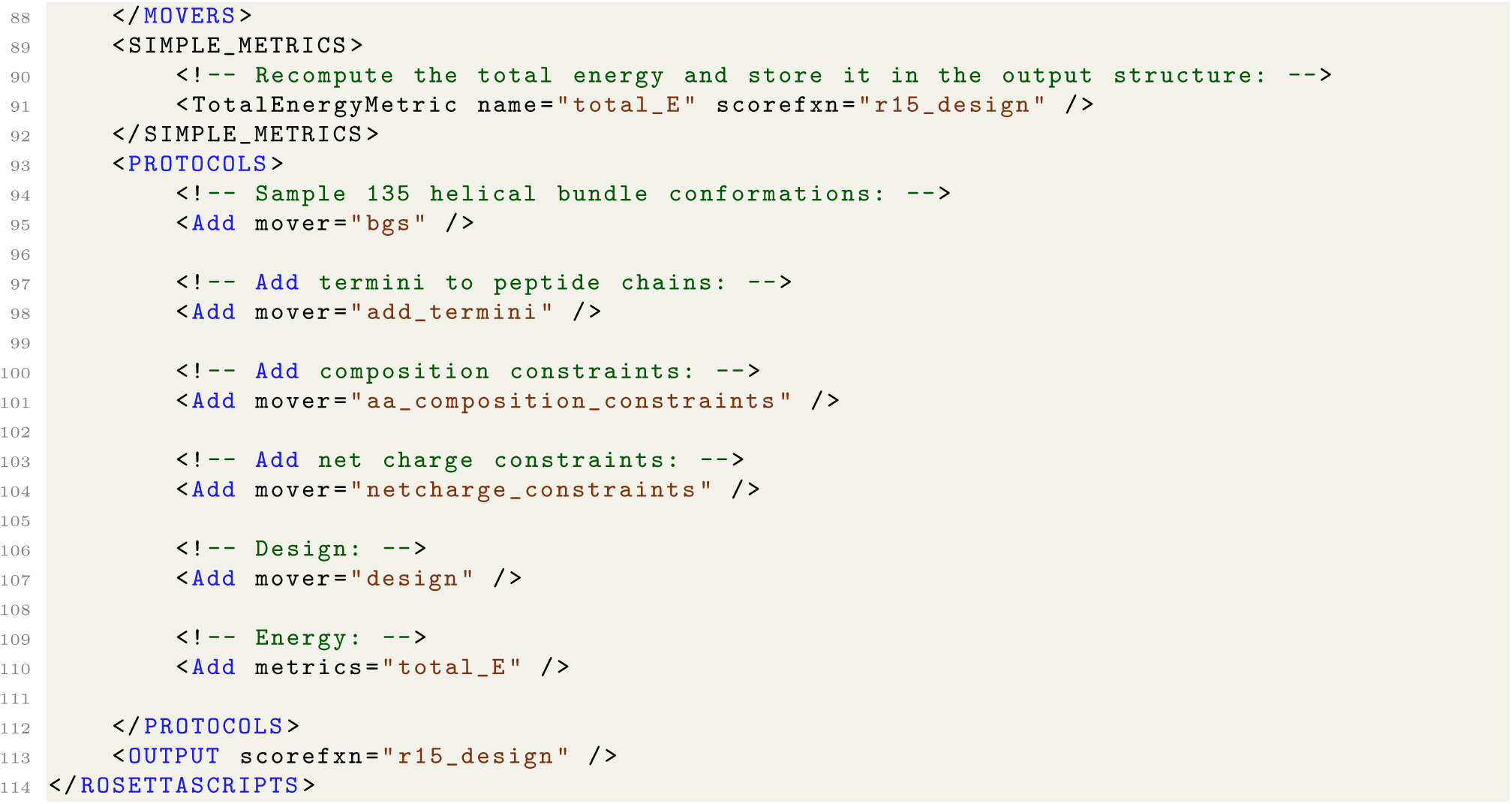

**Listing 5.** RosettaScripts XML defining the helical bundle design protocol used for benchmarking the Rosetta `packer` against Masala’s `MonteCarloCostFunctionNetworkOptimizer`. Script variables, enclosed in double percent signs, were set on the command-line to activate or deactivate particular design-centric guidance functions, or to enable extra rotamers to make problems larger (with the potential benefit of allowing better solutions to be found, given finer discretization of the rotamers).

The above script was dependent on the additional files `global.comp` and `global.charge`, defining global amino acid composition constraints and global charge constraints. These are shown in Listings 6 and 7.

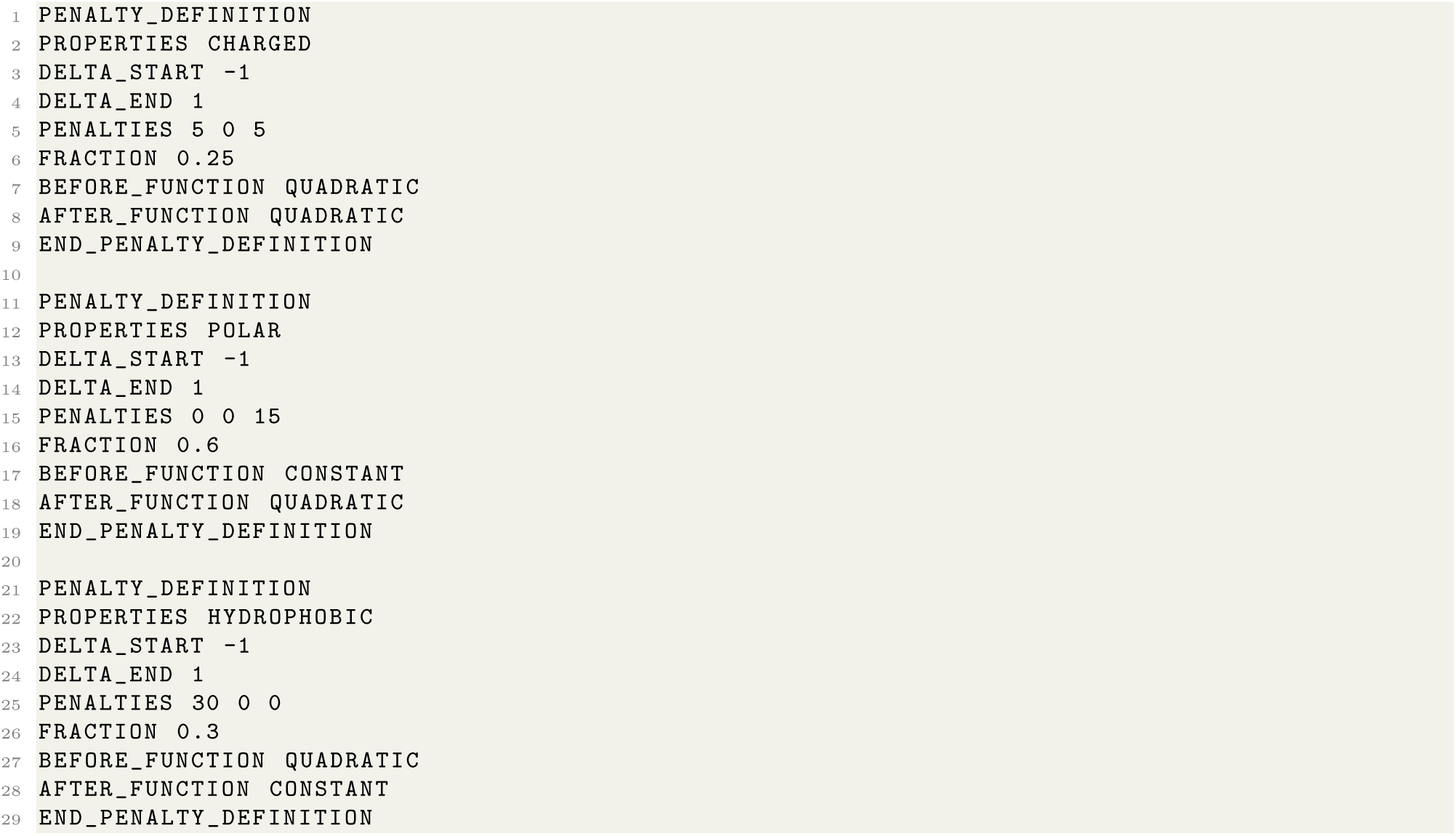
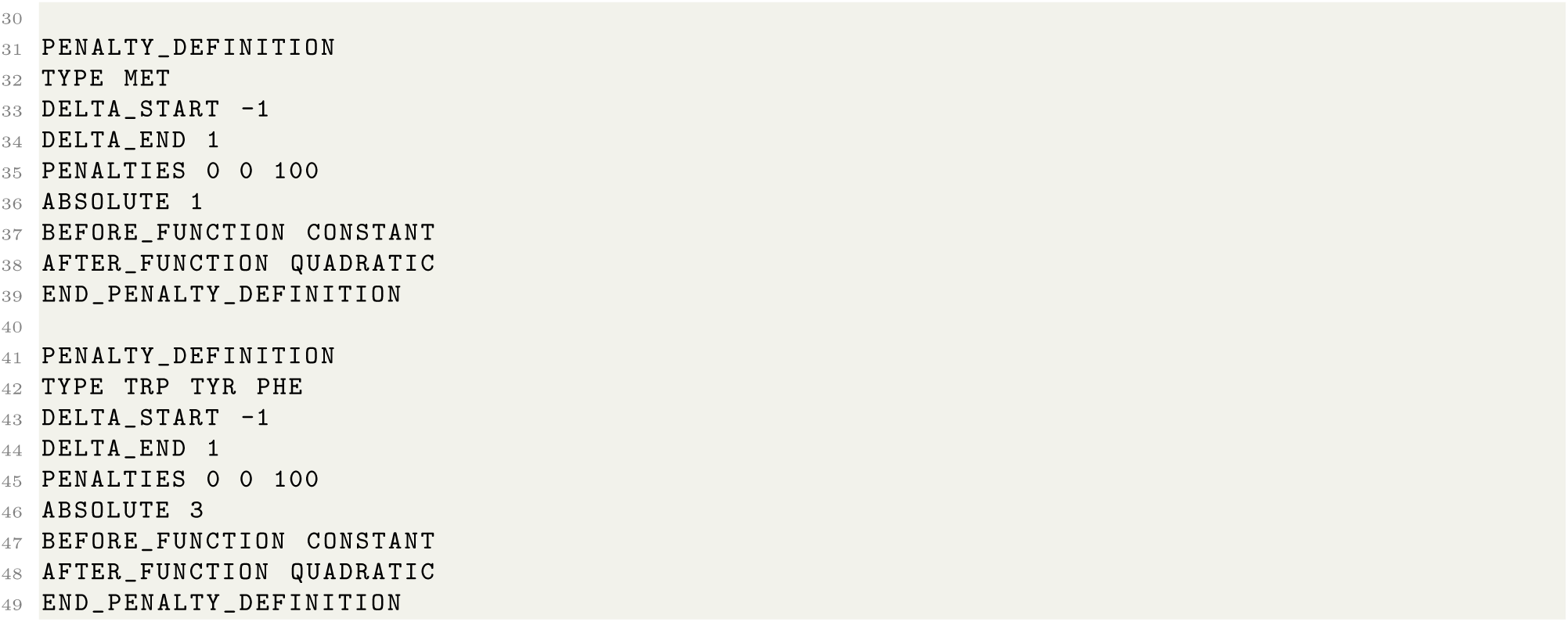

**Listing 6.** Contents of file `global.comp`, defining global amino acid composition constraints. This file specifies that the designs should be 20% charged, at most 60% polar, at least 30% hydrophobic, and should contain at most 1 methionine residue and 3 aromatic residues.

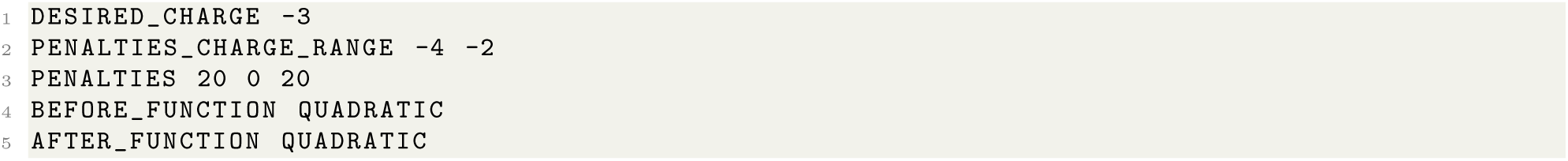

**Listing 7.** Contents of file `global.charge`, defining global net charge constraints. This file simply specfies that the net charge of the molecule should be -3.

The benchmark also requires a file masala cfn solver.xml (Listing 8) to configure the PackRotamersMover to use a Masala CFN solver engine in place of the Rosetta packer.

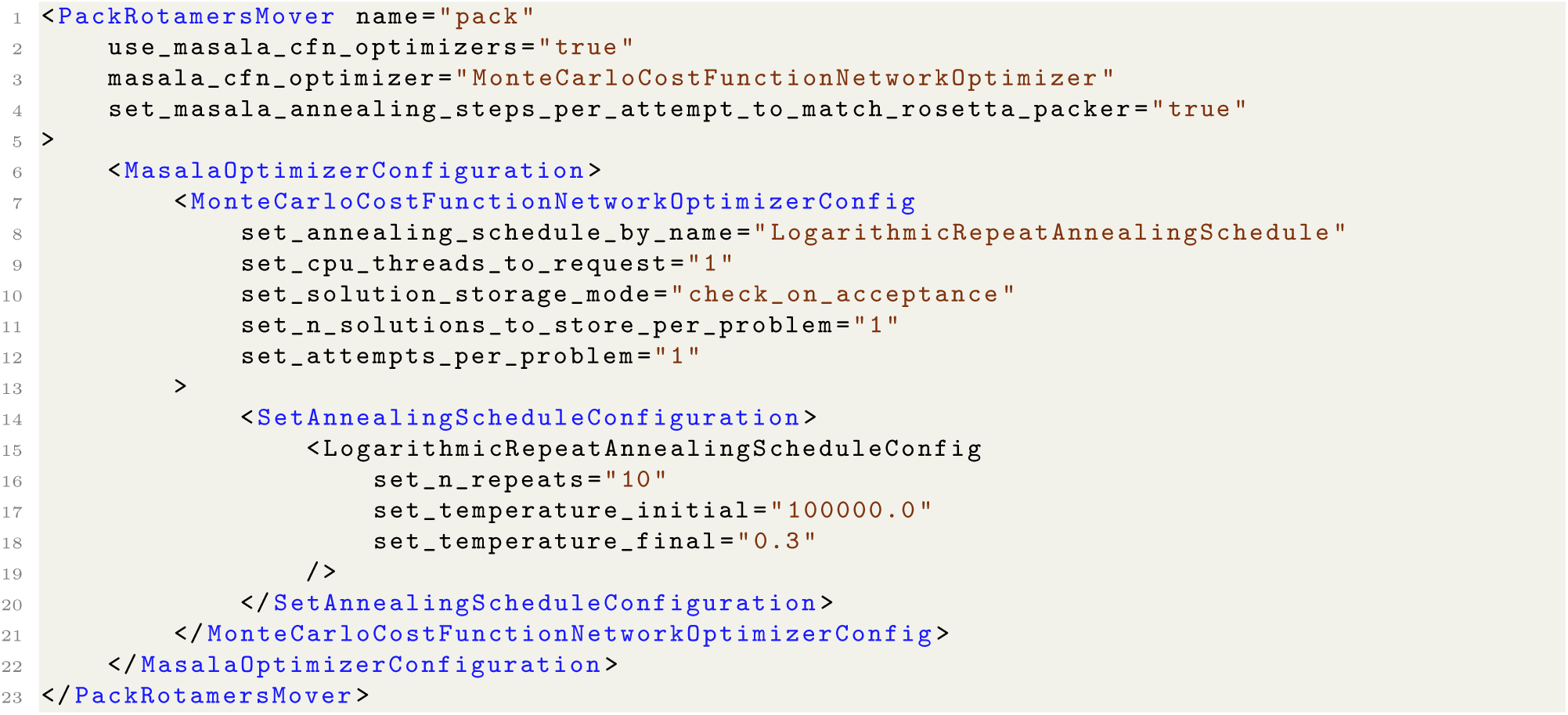

**Listing 8.** Contents of file `masala cfn solver.xml`, used to configure the `PackRotamersMover` to use the Masala `MonteCarloCostFunctionNetworkOptimizer` in place of the Rosetta `packer`.

Finally, RosettaScripts needs a dummy input sequence that will be discarded when the helical bundle is generated. This is provided as a sequence of single-letter amino acid codes (any sequence will do) in a `dummy.fasta` file. Assuming that the design script is in the `xml` sub-directory of the working directory, that the `dummy.fasta`, `global.comp`, `global.charge`, and `masala cfn solver.xml` files are in an `inputs` subdirectory, and that the `$MASALA_STANDARD_PLUGINS` environment variable points to the Standard Masala Plugins directory, the benchmark can be executed with the command shown in Listing 9. Note that the various script variables set which design-centric guidance terms are active, as well as optionally activating the “extra rotamers” options which provide finer discretization of conformational rotamers at the expense of a larger solution search space.

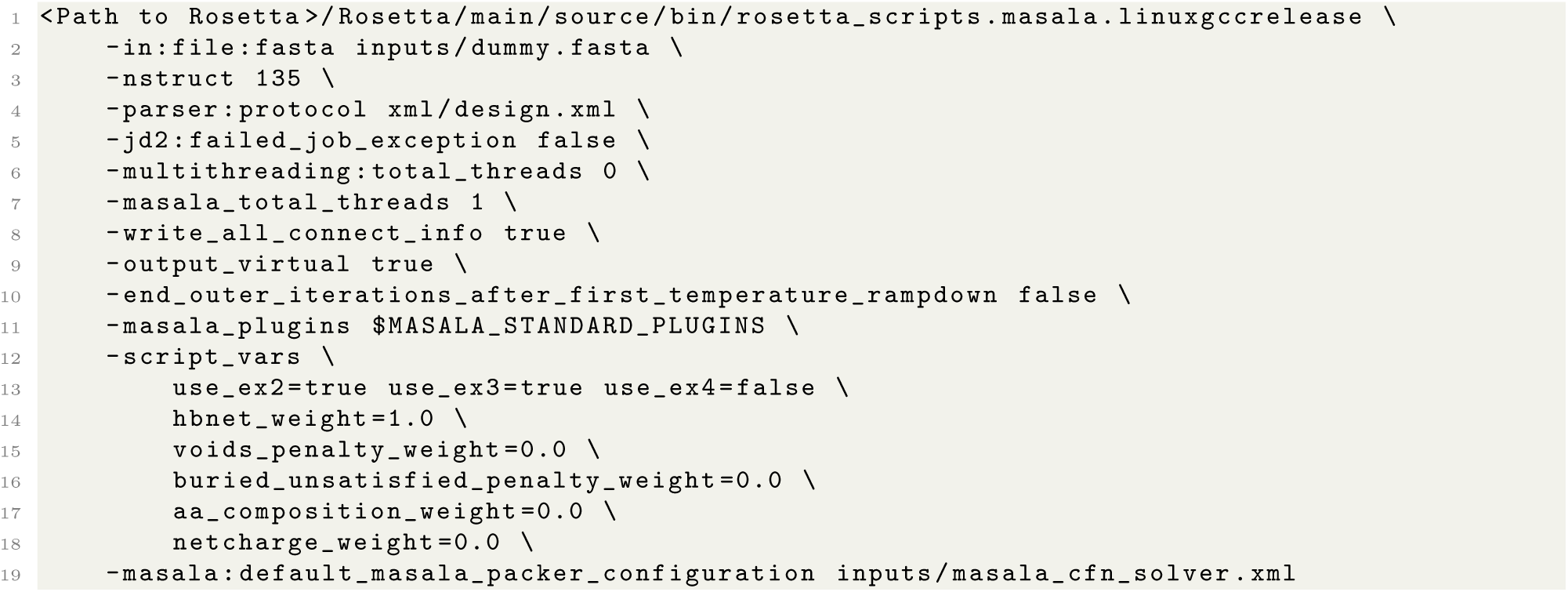

**Listing 9.** Command to execute the single-threaded Masala CFN performance benchmark. This particular example is configured to use the Masala `MonteCarloCostFunctionNetworkOptimizer` in place of the `packer`, to activate the `hbnet` design-centric guidance term, and to use finer discretization of rotamers at the first three sidechain dihedrals. The`-script_vars` option can be altered to change these settings. The `-masala:default_masala_packer_configuration`option can be omitted to pack with the Rosetta `packer` instead of the Masala CFN solver engine.

Since we were primarily interested in benchmarking speed and not accuracy, our measure of performance was the time taken to carry out a simulated annealing trajectory as long as that which the Rosetta `packer` (which ordinarily uses trajectory lengths of 100 times the number of rotamers in the problem) would perform. Results are shown in **Fig. 4**. Although the more general implementation for non-pairwise terms results in the wholly pairwise-decomposable `ref2015` energy function being very slightly slower to evaluate on a single CPU thread (**Fig. 4A**), this small performance decrease is less than twofold, and the ability to run efficiently in two or more threads (see section 4.2) means that this is gained back through parallelism. More importantly, any real-world design task involves using one or more design-centric guidance functions, and we found that our re-implementations accelerated CFN solving with non-pairwise guidance functions considerably in all cases except the `netcharge` term (which was neither faster nor slower with Masala). These speedups were considerable, generally being more than a factor of two (**Fig. 4B**-**F**). Notably, the slowest guidance function, the `buried_unsatisfied_penalty`, showed up to two orders of magnitude of speedup with Masala, as well as better performance scaling with problem size (**Fig. 4F**). The net effect of this is that, when problems are executed with all five design-centric guidance terms enabled (the most plausible real-world scenario), Masala provides a speedup of between half an order of magnitude and two orders of magnitude, with the biggest speedups applying to the largest problems (**Fig. 4G**) — *even* when executing on a single CPU core. Moreover, the data representations implemented here are not necessarily the most performant possible. One of Masala’s big advantages is the potential to implement more efficient data representations in the future, and to have these replace the ones currently used without having to refactor Rosetta code or recompile Rosetta.

**Fig 4.**
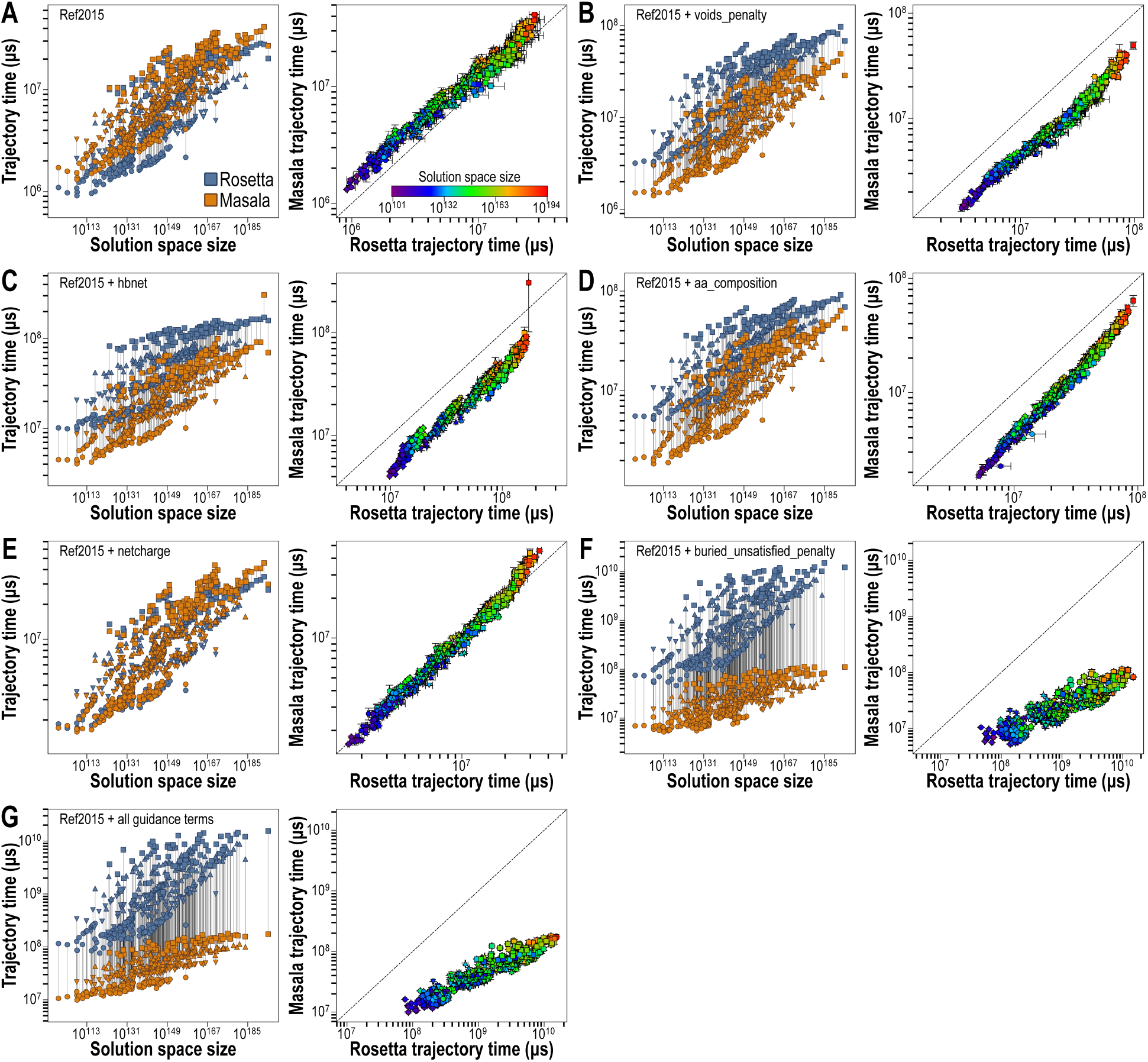
Single-threaded performance benchmark of the Masala `MonteCarloCostFunctionNetworkOptimizer` with the Rosetta `ref2015` scoring function and various combinations of design-centric guidance terms, across a range of problem sizes. Problems were carried out with finer discretization of rotamers at the first sidechain dihedral (circles), at the first and second (downward triangles), at the first through third (upward triangles), and at all sidechain dihedrals (squares). (Left part of each panel) Time to complete a trajectory with steps equal to 100 times the total number of rotamers, plotted as a function of problem size (number of possible solutions). Masala performance is shown in orange, and Rosetta performance in blue. Lines connect the same problem. (Right part of each panel) Masala trajectory time versus Rosetta trajectory time. Colours indicate the size of each problem. Dashed line is the line of equal performance. Error bars are the standard error of three replicates. (**A**) Single-threaded performance with `ref2015` alone. (**B** through **F**) Single-threaded performance of `ref2015` with `voids_penalty`, `hbnet`, `aa_composition`, `netcharge`, or `buried_unsatisfied_penality`, respectively. In all cases but `netcharge`, Masala shows a considerable performance advantage. Performance with `netcharge` is the same for Masala or Rosetta. (**G**) Single-threaded performance of `ref2015` plus `voids_penalty`, `hbnet`, `aa_composition`, `netcharge`, and `buried_unsatisfied_penalty`. The performance of Masala is orders of magnitude better than that of Rosetta, particularly at larger problem sizes. This best approximates a real-world design scenario.

### 4.2 Multi-threaded parallel CPU performance

Since November 2019, Rosetta has been able to parallelize the pre-computation of one- and two-body energies for rotamer optimization problems across CPU threads. Use of mutable data, use of shared pointers, and other architectural details in the Rosetta `packer` and related classes hindered efforts to parallelize the simulated annealing search of rotamer assignment space itself, however. In contrast, Masala was structured with this in mind, and the Standard Masala Plugin library’s `MonteCarloCostFunctionNetworkOptimizer` can carry out parallel simulated annealing trajectories in parallel threads. We carried out multi-threaded performance benchmarks of the `MonteCarloCostFunctionNetworkOptimizer` on three high-performance computing node architectures (a 64-core Intel Ice Lake node, a 96-core AMD Genoa node, and a 128-core AMD Rome node), as well as on a Macbook Pro laptop with a 16-core Apple M3 Max CPU. Benchmarking was performed with the Standard Masala Plugins library’s `benchmark_monte_carlo_cfn_optimizer` standalone application, which measures the time to carry out *N* simulated annealing trajectories of ten million steps in *N* threads for a test CFN problem, varying the thread count from 1 to the number of CPU cores available on the system and performing ten replicates with each thread count. All replicates of all thread counts were shuffled into random order to avoid artifacts that could be caused by a CPU altering its power usage when low thread counts are consistently being requested.

As shown in **Fig. 5**, the Standard Masala Plugins library’s `MonteCarloCostFunctionNetworkOptimizer` shows excellent performance scaling with thread count, particularly on high-performance compute nodes. On Intel Ice Lake and AMD Genoa nodes, better than 90% performance efficiency (defined as observed performance scaling divided by theoretically perfect performance scaling, based on linear extrapolation of performance scaling on a single core) was achieved to 90% saturation of a node. On the older AMD Rome architecture, multi-threaded performance was slightly worse, but better than 90% efficiency was observed to 55% saturation of the node, and better than 50% efficiency was maintained to saturation. (Note that multi-threaded performance depends both on writing software to take efficient advantage of multiple CPU cores, and on the CPU hardware itself.) Remarkably, on the AMD Genoa hardware, we were able to achieve peak performance of 746 *million* possible rotamer assignments considered per second on a single node. Our benchmark demonstrates that Masala’s architecture creates no barriers to the development of highly efficient multi-threaded algorithms.

**Fig 5.**
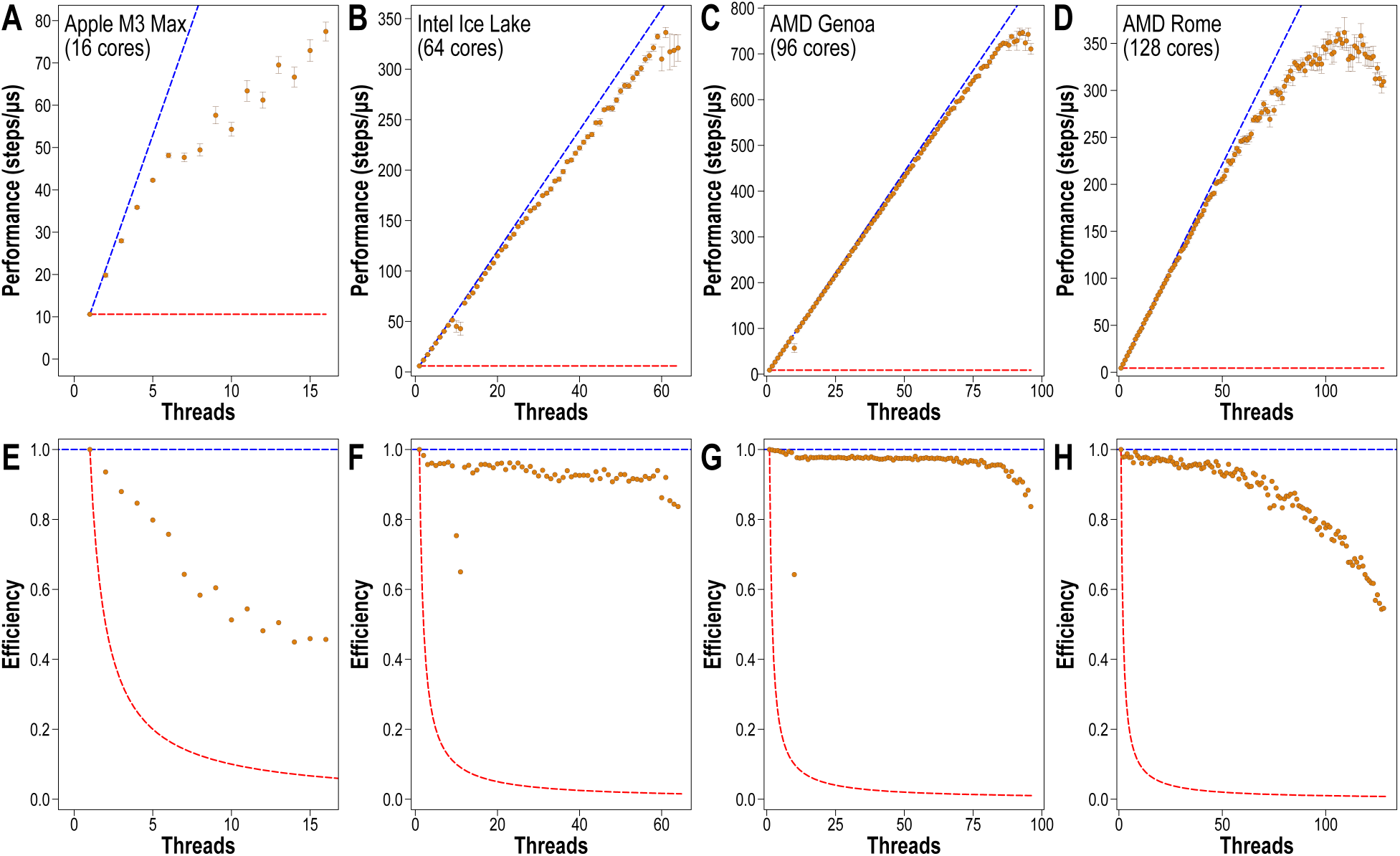
Multi-threaded performance benchmark for Masala’s `MonteCarloCostFunctionNetworkOptimizer`. (**A**-**D**) Absolute performance, measured as number of simulated annealing trajectory steps performed per second, as a function of software threads launched, on a Macbook Pro with a 16-core Apple M3 Max CPU (**A**), a 64-core Intel Ice Lake HPC node (**B**), a 96-core AMD Genoa HPC node (**C**), and a 128-core AMD Rome HPC node (**D**). Ten replicates of each measurement were performed, and all replicates were carried out in random order. Error bars show standard error of the mean across ten replicates. The dashed blue line shows theoretical best-case performance, based on linear extrapolation of single-threaded performance, and the dashed red line shows worst-case performance, with no improvements beyond the single-threaded performance. (**E**-**H**) Performance efficiency, defined as observed performance as a fraction of theoretical best-case performance, for the same hardware types shown in panels **A** through **D**. The blue dashed line shows the best-case scenario (no loss in efficiency at any thread count), and the red dashed line shows worst-case efficiency (no improvement in performance beyond the single-threaded case). In all cases, the observed performance was much closer to the best-case scenario than to the worst-case.

## 5 Future extensibility: The roadmap beyond Masala 1.0

Masala 1.0 is a functional platform for developing numerical methods for designing cyclic peptides and other folding heteropolymers, and for permitting these methods to be rapidly incorporated into production design pipelines and tested at scale against existing state-of-the-art methods. It also represents a step towards robust standalone software for implementing complete design and validation pipelines, and for executing these pipelines at scale on HPC clusters.

**Fig. 6A** shows Masala’s current functionality: other software, such as Rosetta, can use Masala’s optimizers, including its cost function network (CFN) and real-valued local (RVL) optimizers, as drop-in replacements for existing algorithms used for rotamer optimization/sequence design and for relaxation of structures, within existing design pipelines. As new Masala optimizers are added, these can be linked to Rosetta (or other software) at runtime, without making changes to Rosetta’s source code or recompiling Rosetta, permitting much faster rounds of development and testing. Even user interfaces for new Masala modules can be automatically generated, without developer effort, thanks to Masala’s API definition system. Most current effort has focused on numerical methods, however. As shown in **Fig. 6B**, for version 2.0 of Masala, we plan to fully develop the `MolecularSystem` class and the related data types that it contains. This will permit rapid prototyping of entire modelling manipulation or analysis operations (equivalent to Rosetta’s `Mover`, `Filter`, or `SimpleMetric` subclasses) as Masala plugin modules, again allowing runtime incorporation into Rosetta pipelines, with auto-generated user interfaces, without needing to alter Rosetta code or recompile Rosetta. For version 3.0 of Masala (**Fig. 6C**), we will implement a full Masala job distribution system, replacing Rosetta’s system of distributing one sample per job with a system of distributing one *batch* of samples per job. This will permit maximally efficient parallel execution on the CPU cores of a node, or on GPUs, by allowing many threads to carry out a single operation concurrently on many designs (the single instruction, multiple data, or SIMD, paradigm). The MPI job batch distribution will permit many design tasks to be distributed across many nodes.

**Fig 6.**
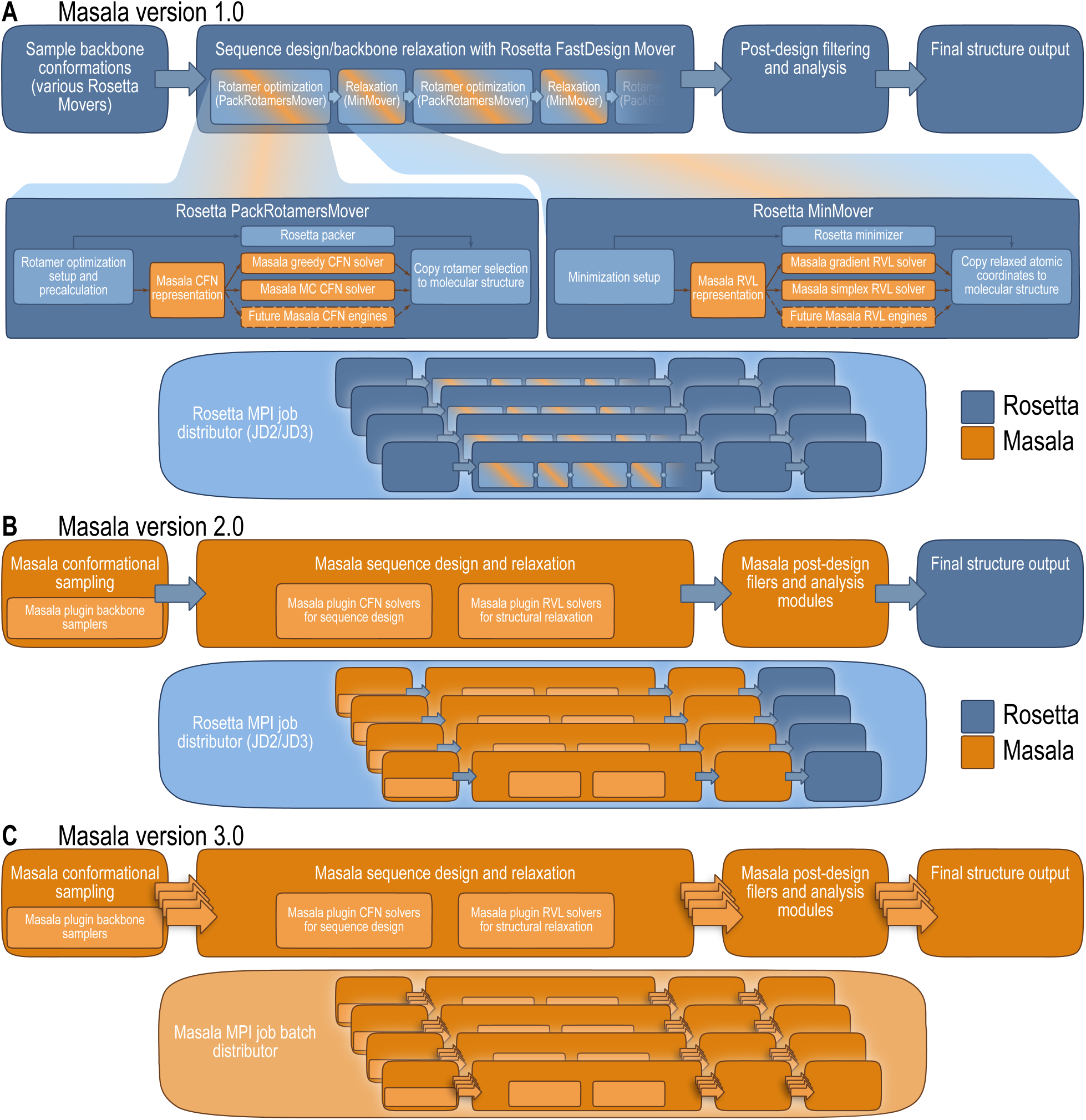
Roadmap for Masala development. Rosetta components are shown in blue, and Masala components, in orange. (**A**) Masala 1.0 (current release) replaces Rosetta optimizers (the `packer` and the `minimizer`), which collectively account for the majority of Rosetta’s runtime, with plugin Masala CFN and RVL optimizers (potentially able to harness finely-grained CPU or GPU thread parallelism) loaded at runtime. This allows rapid cycles of development to experiment with new means of designing and relaxing peptides and proteins. Samples (independent designs) are carried out in parallel by way of Rosetta’s MPI job distributor. (**B**) Masala 2.0 (planned) will flesh out the `MolecularSystem` class, and implement full molecular modelling and analysis protocols as plugin Masala modules while still relying on Rosetta’s MPI job distributor for coarsely-grained parallelism. (**C**) Masala 3.0 (planned) will permit full standalone execution, with coarsely-grained parallelism provided by an MPI job *batch* distributor to distribute work across HPC nodes, and finely-grained parallelism provided by synchronous execution of samples within a batch on a single node’s CPU or GPU by Masala plugin modules written for synchronous concurrency.

## 6 Conclusion

Unlike conventional software development, the creation of scientific software is intrinsically an ongoing process of experimentation. In this process, it must be possible to rapidly try new ideas in their full complexity, in the context of larger pipelines, and to accept or reject experimental ideas based on their merits, without being pigeon-holed into using one particular approach simply because it was what was settled on at one time. Much of the success of the Rosetta software suite, which for decades has been among the most popular macromolecular design, analysis, and prediction packages, has come from its versatile, modular nature, which enabled a good deal of experimentation. A decade and a half after the introduction of Rosetta 3, we have now learnt many lessons about what works and what does not in an experimental software package. This has enabled the creation of Masala, a next-generation macromolecular design and validation software toolkit intended to preserve Rosetta’s strengths while addressing its weaknesses.

In this work, we have introduced the features of the Masala software that make it uniquely well-suited for biologists and biochemists to develop new approaches for macromolecular modelling, without necessarily having extensive experience in software engineering. We have described Masala’s unique API definition system, which allows scientists to focus on scientific programming while automating most of the burdensome infrastructural and user interface programming that often plagued Rosetta development. And we have introduced Masala’s plugin infrastructure, which allows new ideas to be prototyped rapidly in self-contained libraries, with minimal amounts of hand-written code, and with automated generation of all the code needed for runtime linking. This permits new ideas to be tested immediately in existing macromolecular modelling pipelines.

Version 1.0 of Masala is built to be a set of extension libraries for existing software, such as Rosetta. This enables a distributed development model, with developers retaining command of their own modules, and permits libraries to be downloaded and used or discarded as users see fit, so that only the best ideas need to be perpetuated, while approaches that may not work can be pruned. This avoids the bloat that has plagued Rosetta and led it to swell to over three million lines of code. We have also described a roadmap to Masala becoming fully-featured, standalone software, offering a replacement for Rosetta.

## 7 Appendix A: Masala coding conventions

Broadly, Masala coding conventions have been chosen to be familiar to Rosetta developers. Exceptions are noted here.

### 7.1 Names, namespaces, and directory structure

Masala C++ code uses the suffices `.hh` for header files, `.cc` for implementations, and `.fwd.hh` for forward declarations. Template code is placed in files with suffix `.tmpl.hh`. Classes are given camel-case names (for example, `MonteCarloCostFunctionNetworkOptimizer`), and functions and variables are given lower-case names with box-car separations (for example, `run_mc_trajectory()`). Member variables of classes are given a trailing underscore (for example, `attempts_per_problem_` ). Longer, more descriptive names are favoured over shorter ones. All of these are conventions familiar to Rosetta developers.

Unlike Rosetta code, by convention, all Masala classes and functions are assigned a common root namespace for the Masala library. For example, the Core Masala library uses namespace `masala`, while the Standard Masala Plugins library uses namespace `standard_masala_plugins`. Within each library, code is located in sub-directories of a `src/` directory. Classes are defined in C++ files with names matching the class name. Sub-namespaces are assigned to match the directory structure: for example, the `MonteCarloCostFunctionNetworkOptimizer` class, defined in the Standard Masala Plugins library, is located in directory `src/optimizers/cost_function_network`, and in namespace `standard masala plugins::optimizers::cost_function_network`.

### 7.2 Doxygen documentation

Hand-written source code is commented with Doxygen tags, to permit automatic generation of source code documentation. Each function should at least be given a /// @brief description. Auto-generated API libraries (see section 2.2.3 and Appendix B, section 8.1.1) have auto-generated Doxygen tags produced from the API definitions.

We host Doxygen source code documentation for the Masala Core library online at https://users.flatironinstitute.org/~vmulligan/doxygen/masala_core_doxygen/index.html, and for the Standard Masala Plugins library online at https://users.flatironinstitute.org/~vmulligan/doxygen/masala_standard_plugins_doxygen/index.html.

### 7.3 AI-generated source code

For security, reliability, and intellectual property reasons, the use of large language models (LLMs) or other generative AI to write source code is strongly discouraged. If any AI-generated code is included in any Masala library, authors are encouraged to *very* carefully check it by hand, to cover it with hand-written unit tests (see section 7.11) and to enclose it in an `#ifndef PROHIBIT_AI_GENERATED_CODE … #endif` block. The latter will permit all AI-generated code to be disabled if necessary, or found easily should ever the need arise to check all AI-generated code. In version 1.0 of Masala and of the Standard Masala Library, there is no AI-generated source code.

### 7.4 Numerical types

Like Rosetta, Masala defines custom unsigned integer (`masala::base::Size`) and high-precision floating point (`masala::base::Real`) data types. By default, these are defined to be `std::size t` and `double`, respectively; however, by defining these and using them throughout the codebase, developers can rapidly benchmark the impact of changing numerical precision on complex protocols simply by altering the `typedef` statements in `src/base/types.hh` in the Masala Core library.

### 7.5 Raw pointers, smart pointers, and the new and delete keywords

As is true of Rosetta, Masala strives to avoid memory leaks by making heavy use of smart pointers (`std::shared_ptr`). However, some of the conventions are different. Where Rosetta functions frequently pass objects about using a `std::shared_ptr` to an object, Masala strives to have object lifetimes managed by a shared pointer at the root of a callstack, but to pass the object about using references (or, when it is possible that the shared pointer is `nullptr`, by raw pointer). This is for multi-threaded efficiency: when shared pointers’ reference counts are frequently incremented or decremented, threaded performance can suffer, since each increment or decrement requires transient locking of the shared pointer object.

An object managed by a smart pointer is expected to be allocated with `masala::make shared<T>(·)`, an alias for `std::make shared<T>(·)`. The use of the C++ keywords `new` and `delete` in Masala code is heavily discouraged in order to avoid dangling pointers, memory leaks, and double-free errors.

Finally, note that the Masala aliases `MASALA SHARED POINTER<T>`, `MASALA WEAK POINTER<T>`, and the allocation function `masala::make shared<T>(·)` should be used instead of directly using the Standard Template Library names `std::shared_ptr<T>`, `std::weak ptr<T>`, and `std::make shared<T>(·)`. This is described in greater detail in section 8.1.4 of Appendix B.

### 7.6 Zero-based indexing

Unlike Rosetta, which inherits one-based array and vector indexing from its FORTRAN days, Masala uses zero-based indexing to be more familiar to C++ and Python developers.

### 7.7 Naming functions

Each Masala class must implement public functions providing the class name (`std::string class_name() const override`) and namespace (`std::string class_namespace() const override`). These are overrides of pure virtual functions in the `MasalaObject` base class, and without these, the derived class remains pure virtual. It is recommended that each Masala class also implement static versions of these functions (named `class_name_static()` and `class_namespace_static()`) to be called by the non-static versions. These name and namespace functions are essential for the automatic API generation system (described in section 2.2.3 and Appendix B, section 8.1.1). They also facilitate output labelled with the module that produced the output.

### 7.8 The clone(), deep clone(), and make independent() class methods

Masala pure virtual base classes typically declare a pure virtual `clone()` function which must be implemented by derived classes. This function makes a copy of the original object and return a base class shared pointer to the copy. Note that this copy is *not* guaranteed to be independent: stored data could include pointers to other objects which remain shared.

The possibility of cloned objects remaining connected through shared data creates opportunities for difficult-to-diagnose bugs, particularly related to thread safety. This has proven to be a problem with Rosetta’s codebase. For this reason, in addition to the `clone()` function, Masala classes should implement a `make_independent()` function designed to deep-clone all of the private data stored within a class instance, and a `deep_clone()` function to deep-clone the class instance itself, returning a shared pointer to a wholly independent copy of a class instance. Masala classes with an API definition *must* implement `deep_clone()`, and the Masala build system will produce an error if they do not (since auto-generated API classes call this function when implementing their own `deep_clone()` functions). Outside of the API definition system, in contexts in which one is likely to have a pointer to a base class rather than to the actual derived class, it is recommended to call `clone()` and then call the new object’s `make_independent()` function rather than relying on `deep_clone()`, since `deep_clone()` is generally not implemented as a virtual function.

It is recommended that `make_independent()` and `deep_clone()` be implemented to call a common protected virtual member function called `protected_make_independent()`. Each class is responsible for overriding this function and implementing code to make all private member data independent by deep-cloning them, then calling the `protected_make_independent()` function of the parent class. The `protected make independent()` function implementations should deep-clone member data by the pattern described above, of calling `clone()` and then calling the copy’s `make_independent()` function.

### 7.9 Multi-threading and thread safety

The Rosetta software suite was built for single-threaded execution; much later, some limited support for multi-threading was added for tasks such as pre-computation of interaction graphs prior to rotamer optimization [2, 22]. One challenge, however, was Rosetta’s modular nature: if code executed in parallel threads were to call Rosetta modules that *also* sought to execute code in parallel threads, a potential thread explosion could occur. Given *N* levels of function calls, each triggering *T* parallel threads of execution, the system might try to execute work in *T ^N^* threads at once. In 2019, we therefore introduced a `RosettaThreadManager`, which maintains a thread pool of user-defined size to which work can be assigned. A similar paradigm has been followed in Masala: modules wishing to perform work in parallel threads must create a `MasalaThreadedWorkRequest`, populate this object with one or more independent, thread-safe tasks that can be performed sequentially or concurrently, and pass this object to the `MasalaThreadManager` for execution in up to as many threads as are requested. If fewer threads are available than are requested, the work executes in whatever threads are available, with a strong guarantee that at least the requesting thread will be assigned the work.

Masala modules are intended to be safe for threaded execution by default; however, writing thread-safe classes is challenging for novice developers. To facilitate this, automatic thread safety is provided by the Masala build system at the level of API classes, described in section 2.2.3. These provide automatically-generated mutex locks whenever a setter, getter, or work function is called. Advanced users wishing to write modules that internally parallelize their work are responsible for ensuring that internal data access is performed in a thread-safe manner, either by implementing appropriate mutexes of their own within their class, or by correctly ensuring that different threads are writing to or reading from different blocks of data.

To ensure thread safety, public and protected member data are prohibited: all member data of classes must be private, and accessed through appropriate getters and setters. Public functions should implement mutex locks (though API container versions of these will implement this automatically should a novice developer forget this). Protected functions often should omit mutex locks, but developer comments should indicate that the function is expected to be called from a mutex-locked context. This permits patterns in which a public function locks a mutex, then calls several different protected functions under the same mutex lock, or calls protected virtual function overrides which call the parent class version of the same function (as in the case of `protected_make_independent()`, described in section 7.8, above).

### 7.10 Global variables and global state

Related to thread safety are considerations of global data and global state. Rosetta protocols that altered global state have long been an obstacle to safe parallelization of Rosetta. Because any `Mover` or `Filter` could allocate and alter static non-constant variables, or even access non-constant global options and change these during execution, parallel threads of execution could be disrupted in unpredictable and difficult-to-reproduce ways. Even the loading of data intended to be global, such as the large rotamer libraries used for sidechain conformational searches (which one would not want to duplicate in each thread), led to data races when threads concurrently triggered lazy loading of these libraries. Much effort later went into refactoring code to implement data management schemes permitting thread-safe lazy loading.

Masala’s convention is to strictly prohibit global data outside of the `masala::base::managers` namespace. Within this namespace (discussed in greater detail in section 8.1 in Appendix B), static singletons are declared that manage needed global data and other elements of global state in a manner safe for parallelism.

### 7.11 Unit tests

Unit tests are implemented using the `Catch2` framework (https://github.com/catchorg/Catch2), and are located in a `tests` sub-directory adjacent to the `src` sub-directory that contains source code. Wherever possible, the directory structure in the unit tests parallels that in the `src` directory. Unit tests are compiled to self-contained executable files, symlinks to which are placed in `bin/tests` by the build system.

## 8 Appendix B: Organization of the Masala Core library, version 1.0

Masala’s Core library contains the central code in Masala, and is linked at compilation time by all external software projects that either link Masala plugin libraries at compilation time, or which are intended to make use of Masala plugin libraries linked at runtime. By design, the Core library avoids including any implementation of any particular engines, data representations, or other functional Masala components intended for molecular modelling: these are all intended to be implemented in plugin libraries, with only abstract base classes implemented in the Core library. What is present in the Core library is all of the infrastructure needed to define the plugin and API definition systems, as well as static singletons for managing hardware, disk access, memory, plugins, engines, data representations, output logging, and anything else sensitive to global state.

This section is intended for developers, and provides a detailed description of the Core library. The library is divided into three parts: the `base` sub-library (section 8.1) contains the lowest-level code, including the aforementioned static singletons. The `numeric` sub-library (section 8.2) contains abstract base classes for numeric methods relevant to molecular modelling. And the `core` sub-library (section 8.3), which is not to be confused with the Core library itself (which we will capitalize to distinguish it), defines abstract chemical concepts, as well as concrete container classes for chemistry-related data, such as the `MolecularSystem` class. Both the `numeric` sub-library and the `core` sub-library have companion auto-generated API sub-libraries (`numeric api` and `core api`, respectively) that serve as the public, stable interfaces.

### 8.1 The base sub-library

The `base` sub-library in Masala’s Core library contains the lowest-level code in Masala, defining much of the central infrastructure for the plugin system, for API generation, and for managing memory, threads, inputs, and outputs. Much of this is handled by static singletons called *managers*, each tasked with ensuring thread-safe handling of requests for shared or limited resources. Additionally, `src/base/types.hh` defines the numerical types commonly used in Masala (see section 7.4 in Appendix A).

All Masala classes derive from a common base class, `MasalaObject`, located in namespace `masala::base`. Derived from this in the same namespace is `MasalaObjectAPI`, a common base class for all API container objects.

Note that the Masala `base` sub-library is the only Masala library that violates Masala’s usual conventions regarding direct access by external code: because the `base` sub-library defines the infrastructure needed for the API definition system, it is necessary for external code to be able to access its classes and functions directly. There is no auto-generated `base-api` sub-library.

#### 8.1.1 Masala API definitions

Masala’s API definition system, described in section 2.2.3, is implemented in code located in namespace `masala::base::api`. Here, the `MasalaObjectAPIDefinition` class is defined, which serves as a container for the full description for the API for a Masala class. In namespaces `masala::base::api::constructor`, `masala::base::api::setter`, `masala::base::api::getter`, and `masala::base::api::work_function`, classes may be found for describing Masala class constructors, setters, getters, and work functions, respectively. Within each of these is a base class for a generic function description, and derived template classes for describing functions of each type with anywhere from zero to ten inputs. For instance, work function descriptions all derive from `MasalaObjectAPIWorkFunctionDefinition` (defined in namespace `masala::base::api::work_function`), and if one wished to include in the API definition for a class a work function that took a constant reference to a string and a boolean value as inputs, and returned a real number as output, one would use the derived template class `MasalaObjectAPIWorkFunctionDefinition_TwoInput< masala::base::Real, std::string const &, bool >`. Constructors for these template classes accept the function name, a human-readable description, names and descriptions of all input and output parameters, and a `std::function` object binding the function for a particular instance of the class. In addition to providing the means by which functions in plugin classes may be invoked by code not aware of the plugin classes at compilation time, including a `std::function` object in the constructor for the function definition also helps to catch developer error by producing a compilation error if the template class specialization does not match the function’s inputs and outputs.

A base class for function annotations is provided in namespace `masala::base::api::function_annotation`. Derived classes for function annotations for constructors (`MasalaConstructorAnnotation`), setters (`MasalaSetterFunctionAnnotation`), getters (`MasalaGetterFunctionAnnotation`), and work functions (`MasalaWorkFunctionAnnotation`) are located in sub-namespaces of the namespaces defining function descriptions, as are derived classes that implement particular function annotations. For instance, work function annotations are defined in namespace `masala::base::api::work_function::work_function annotation`. Function annotations may be attached to function descriptions to mark special features of functions. Of note are the deprecation annotations (`DeprecatedConstructorAnnotation`, `DeprecatedSetterAnnotation`, `DeprecatedGetterAnnotation`, and `DeprecatedWorkFunctionAnnotation`) used to indicate that a function will be deprecated in a certain version of the code, and to cause the build system to automatically remove its equivalent from the API classes once that version is reached (see section 2.2.4). Masala’s deprecation annotations are easily added with a single line of code in the `get api defintion()` function of a class. An example is found in the `MolecularSystem` class (namespace `masala::core::molecular_system`), as shown in Listing 10.

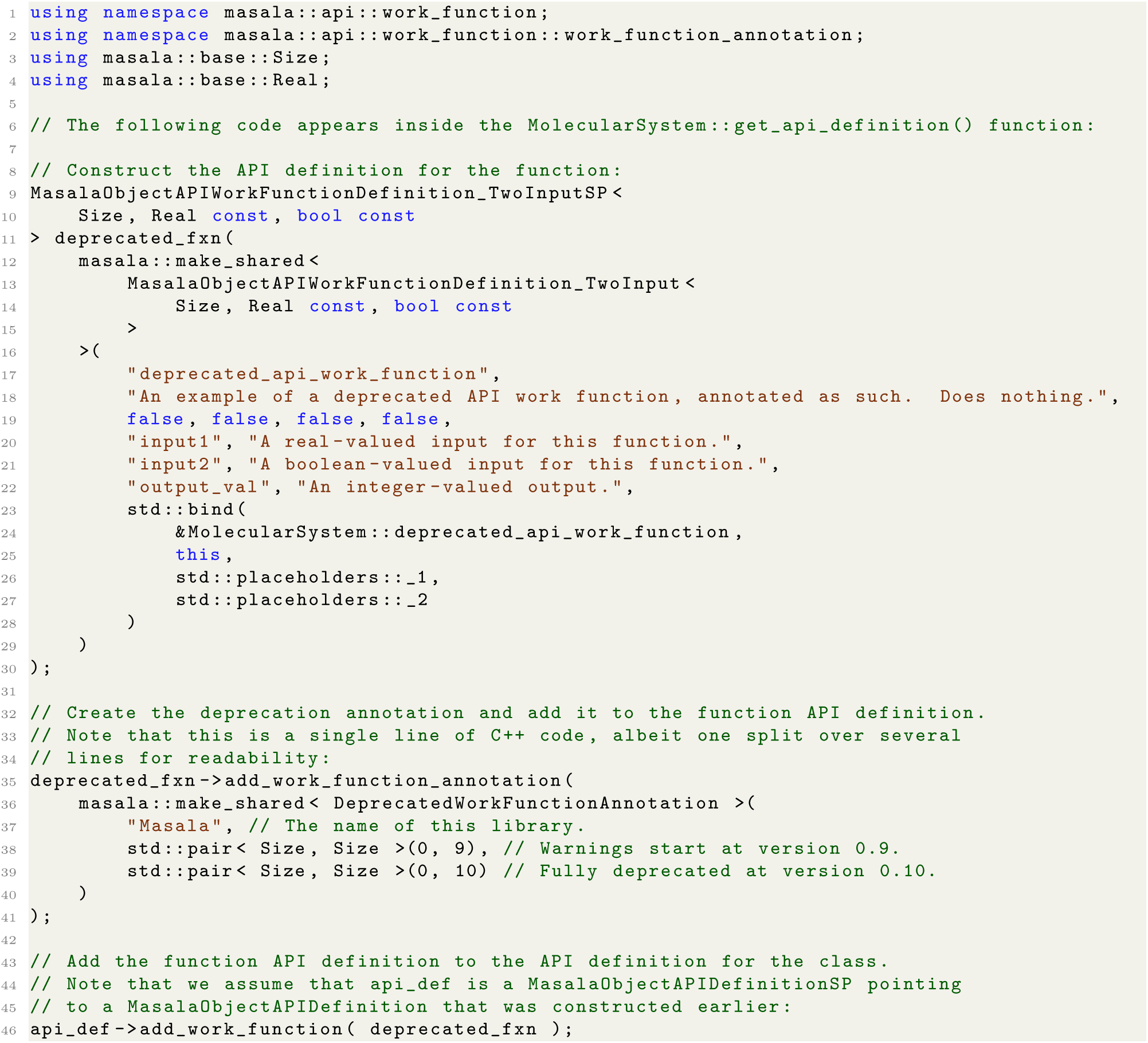

**Listing 10.** An example of the use of Masala function deprecation annotations to mark the `deprecated api work function()` function for deprecation. Note that this code appears within the `get_api_definition()` function of the `MolecularSystem` class.

Also of note are the no-UI annotations (NoUIConstructorAnnotation, NoUISetterAnnotation, NoUIGetterAnnotation, and NoUIWorkFunctionAnnotation), which provide hints to external code suggesting that a function so annotated ought not to be included in user-facing interfaces.

#### 8.1.2 Disk, environment, configuration, and database management

One of the lessons learned from Rosetta is that many issues that hinder performant production runs, particularly when one scales up to running code on large supercomputers, come from poor management of disk input and output (I/O). Even a single, single-threaded process that writes to disk too frequently or in the midst of an intensive calculation can experience significant slowdowns; when hundreds of thousands of parallel processes, each with multiple threads, are all trying to write to disk, jobs can slow to a standstill. In Rosetta, any class or function could write to or read from disk at any time, and much tedious work has gone into tracking down instances of stray disk reads or writes, and implementing better caching of data and more centralized reading and writing.

To avoid this in Masala, Masala modules are not permitted to read from or write to disk directly. All disk access in Masala is required to go through the `MasalaDiskManager`, which ensures thread-safe file reads and writes, and provides a single central location where disk access can be refactored in the future should the need arise (for instance, to add better buffering if many processes are all writing to disk). The `MasalaDiskManager` is located in namespace `masala::base::managers::disk`.

A specialized manager, the `MasalaEnvironmentManager`, has also been implemented to handle reading and caching environment variables lazily, on first demand, to avoid repeated system calls. This is located in namespace `masala::base::managers::environment`. Similarly, a specialized `MasalaConfigurationManager` (namespace `masala::base::managers::configuration`) exists to permit configuration files for particular Masala modules to be lazily read once and cached (permitting, for instance, custom configurations on different hardware without users having to remember to change inputs). Finally, a manager for lazily loading large blocks of potentially protocol-specific data, the `MasalaDatabaseManager`, has been implemented in `masala::base::managers::database`.

#### 8.1.3 Tracer management

The `masala::base::managers::tracer` namespace contains the `MasalaTracerManager`. This is intended to be the central manager of Masala output logs, written either to the screen or to disk. By requiring all Masala modules to go through the tracer manager, central control of verbosity is possible. Additionally, the `MasalaTracerManager` ensures thread-safe output (with originating threads, and, in MPI contexts, MPI ranks labelled in each line of output), preventing broken output from multiple threads writing to the output log simultaneously.

#### 8.1.4 Memory management

Although namespace `masala::base::managers::memory` has been created in case a dedicated memory manager proves necessary in future releases, version 1.0 of Masala does not have one. At present, this namespace contains only utility code that defines the `MASALA_SHARED_POINTER<TYPENAME T>` and `MASALA_WEAK_POINTER<TYPENAME T>` template types (currently defined to be `std::shared_ptr` and `std::weak ptr`, respectively, though having this in one location permits it to be changed in the future should the need arise). Additionally, `masala::make_shared<TYPENAME T>(·)` is defined to be `std::make_shared<TYPENAME T>(·)`, though this too can be redefined should the need arise. Masala convention is to use `MASALA_SHARED_POINTER`, `MASALA WEAK POINTER`, and `masala::make_shared<TYPENAME T>(·)` throughout the Masala codebase, and to avoid naming the STL equivalents directly.

#### 8.1.5 Plugin management

Namespace `masala::base::managers::plugin_module` defines two static singleton classes, the `MasalaPluginLibraryManager` and the `MasalaPluginModuleManager`. The `MasalaPluginLibraryManager` manages the loading and unloading of plugin libraries, keeping track of the names of loaded plugin libraries and of the library handles. Libraries are loaded by calling `MasalaPluginLibraryManager::load_and_register_plugin_library(·)` and passing it a path and filename, or by calling `MasalaPluginLibraryManager::load_and_register_plugin_libraries_in_subdirectories(·)` and passing it a directory path. The latter will search for `registration api.dll` (Windows), `registration api.so` (Linux), or `registration api.dylib` (MacOS) files and attempt to load these. Libraries that lack an `extern "C" void register_library()` function will result in an error being thrown. The `MasalaPluginLibraryManager::reset()` function will unload all loaded plugin libraries, purging the classes registered with other managers.

The `MasalaPluginModuleManager` manages individual plugin modules defined in plugin libraries loaded by the `MasalaPluginLibraryManager`. In addition to these two static singletons, the `MasalaPlugin` class, a base class for all plugin modules, is also defined in the `masala::base::managers::plugin_module` namespace.

This class (which itself inherits from `MasalaObject`) has pure virtual functions `MasalaPlugin::get_categories()` and `MasalaPlugin::get_keywords()`. The first must be implemented by derived classes to return a vector of vectors of strings, with each inner vector representing a possible hierarchical classification of the plugin module (for instance, [ “Food”, “Fruit”, “Citrus”, “Tangerine” ] may be the hierarchy for a tangerine, if we were classifying food rather than plugins). A given plugin may be in multiple hierarchical categories, and the categories allow external code to search for particular types of plugins (for instance, solvers for particular types of problems). The second function, `MasalaPlugin::get keywords()`, must be implemented by derived classes to return a vector of keyword strings. These keyword strings are intended for user interfaces, to permit users to search for particular plugin modules to use for particular tasks. For instance, one may search for “gpu” if one wanted to use a GPU compute node to solve a particular problem and needed a plugin module compatible with GPU computing. A plugin library’s `extern "C" void register_library()` function registers each plugin in the library with the `MasalaPluginModuleManager`. The `MasalaPluginModuleManager` holds a map of categories to sets of `MasalaPluginCreatorCSP` smart pointers. Each of the `MasalaPluginCreator` objects to which these pointers point is a dedicated class that instantiates a particular type of plugin object. Creators are auto-generated by the build system (see section 2.2.3), and all derive from `MasalaPluginCreator`, a common base class in the same namespace. External code requesting a particular plugin module or type of plugin module receives a vector of `MasalaPluginCreator` objects, each of which can be used to create instances of the particular object type by calling `MasalaPluginCreator::create_plugin_object()`. This function returns a suitable subclass of a `MasalaPluginAPI` container object by `MasalaPluginAPISP` shared pointer. The `MasalaPluginAPI` base class (which inherits from `MasalaObjectAPI`) is also defined in the `masala::base::managers::plugin_module` namespace, and the container classes for derived `MasalaPlugin` objects are auto-generated by the build system, and follow the same hierarchy as the classes that they contain. All `MasalaObjectAPI` auto-generated derived classes implement `get_api_definition_for_inner_class()`, permitting the API definition for the contained class to be retrieved.

As described in sections 8.1.6 and 8.1.7, certain classes of plugins (*file interpreters*, *engines*, and *data representations*) are given privileged status, and have their own managers to permit efficient retrieval of appropriate modules, though all of these can also be retrieved through the general MasalaPluginManager.

#### 8.1.6 File interpreter management

*File interpreters* are a privileged class of Masala plugin modules. Each is responsible for reading a particular file format and converting the data contained to a suitable Masala data representation in memory, and/or for writing particular a file format based on data in memory. The `masala::base::managers::file_inter-preters` namespace defines all classes related to file interpreters. These include `MasalaFileInterpreter`, a base class for all file interpreters that itself inherits from `MasalaPlugin` (see section 8.1.5), `MasalaFileInterpreterAPI`, a base class for all API container classes that contain file interpreters that inherits from `MasalaPluginAPI`, and `MasalaFileInterpreterCreator`, a base class for all file interpreter creator objects that inherits from `MasalaPluginCreator`.

The `masala::base::managers::file interpreters` namespace also contains the `MasalaFileInterpreterManager`. This is a static singleton in which all file interpreters are registered. Whenever a set of plugin modules is registered with the `MasalaPluginModuleManager` (see section 8.1.5), the `MasalaPluginModuleManager` checks each to determine whether it is a file interpreter. Modules identified as file interpreters are automatically registered with the `MasalaFileInterpreterManager`. External code linking the Masala Core library may interrogate either the `MasalaPluginModuleManager` or the `MasalaFileInterpreterManager` to get file interpreters for particular file formats. However, using the latter is slightly advantageous, first because one is then guaranteed to retrieve *only* file interpreters (whereas the generic plugins that one retrieves from the `MasalaPluginModuleManager` would have to be dynamically-cast to determine whether they were file interpreters), and second, because the `MasalaFileInterpreterManager` offers convenience functions for searching for file interpreters that read or write particular file types. Specifically, the function `MasalaFileInterpreterManager::get_file_interpreters_by_file_type_descriptor(·)` returns a vector of file interpreter creators producing those file interpreters that read or write files with a particular descriptor (where the descriptor may be a string like “protein data bank”). Additinally, the function `MasalaFileInterpreterManager::get_file_interpreters_by_file_type_extension(·)` returns a vector of file interpreter creators producing those file interpreters that read or write files with a certain extension (where the extension may be a lowercase string like “pdb”).

#### 8.1.7 Engine and data representation management

Like file interpreters, *engines* and *data representations* are privileged classes of Masala plugin modules. As discussed in Section 2.2.2, engines are highly optimized modules that solve a particular hard problem on a particular type of hardware as efficiently as possible. To maximize the efficiency, they accept one or more compatible types of data representations, which are containers for the information that defines the problem, and which put the information into a form that maximizes the efficiency with which the problem can be solved. The namespace `masala::base::managers::engine` defines both the `MasalaEngine` and `MasalaDataRepresentation` base classes, both of which derive from `MasalaPlugin` (see section 8.1.5). It also defines the associated `MasalaEngineAPI` and `MasalaDataRepresentationAPI` base classes for API containers for engines and data representations respectively, both of which derive from `MasalaPluginAPI`, as well as `MasalaEngineCreator` and `MasalaDataRepresentationCreator` base classes for creators, both of which inherit from `MasalaPluginCreator`.

**Fig. 7** shows the class inheritence hierarchy for `MasalaEngine`, including a number of numerical solvers implemented in the Standard Masala Plugins library that are discussed in Appendix C (section 9). Each `MasalaEngine` defines a list of engine categories and engine keywords by providing implementions of the pure virtual base class methods `MasalaEngine::get engine categories()` and `MasalaEngine::get engine keywords()`; these may or may not return exactly the same categories and keywords as the similar functions defined in the `MasalaPlugin` base class. In addition, each engine must provide an implementation of the pure virtual base class method `MasalaEngine::data_representation_is_incompatible_with_engine(·)`, which takes a constant reference to a `MasalaDataRepresentationCreator` and returns a Boolean. This allows engines in plugin libraries to determine whether a particular data representation encodes a problem of a type that the engine can solve, either by dynamic casting it to engine subclasses defined in the plugin library, or by interrogating its categories, keywords, or other properties.

**Fig 7.**
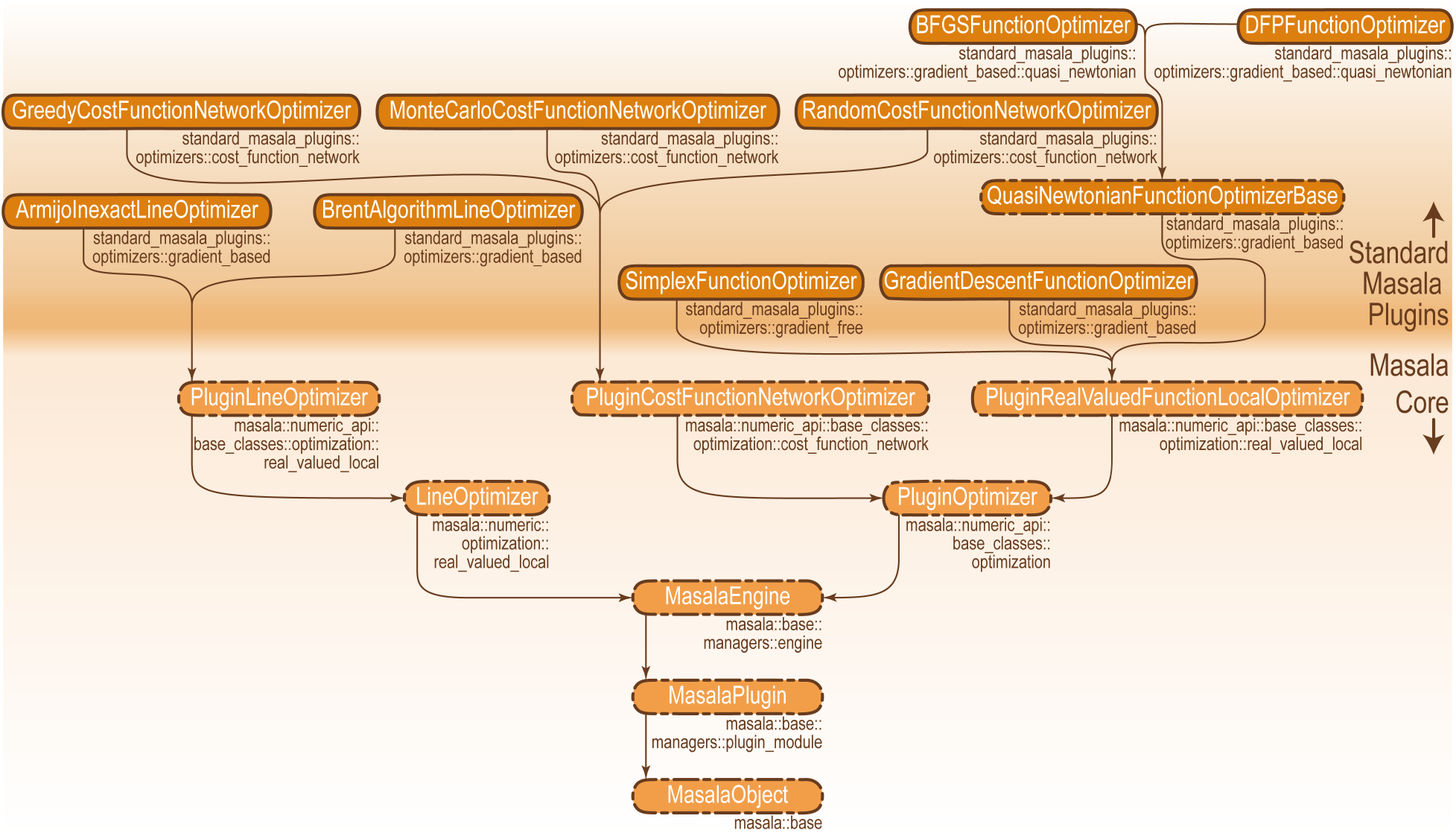
Class inheritance hierarchy for the `MasalaEngine` class. Arrows flow from child to parent classes. Namespaces are shown in smaller print below each box containing a class name. Dashed outlines indicate classes that cannot be instantiated by the Masala plugin system (base classes). Not shown are derived classes in other plugin libraries.

Engines are also managed by a static singleton, the `MasalaEngineManager`, defined in the same namespace, which maintains a list of engine creators by hierarchical engine category. The `MasalaEngineManager` fields requests from external code for sets of engine creators matching particular criteria. Criteria are provided to the `MasalaEngineManager` using `MasalaEngineRequest` objects (another class defined in the same namespace, derived from `MasalaObject`). These contain one or more `MasalaEngineRequestCriterion` objects, defined in namespace `masala::base::managers::engine:engine_request`. A `MasalaEngineRequestCriterion` may require that an engine fall into a particular engine category, have a particular name, have a particular keyword, or, through Boolean logic, require that certain keywords be absent. `MasalaEngineRequestCriterion` subclasses for AND, OR, and NOT logic allow relatively complex rules to be constructed.

Data representations are similary managed by a `MasalaDataRepresentationManager` static singleton, defined in namespace `masala::base::managers::engine`, which maintains a list of data reperesentation creators by hierarchical data representation category. In addition to categories and keywords, data representations can return lists of properties that they have (*e.g.* “linearly scaling”, or “for sparse matrices”), and may optionally return nonexhaustive lists of properties that they lack. They also may optionally return nonexhaustive lists of known engines with which they are compatible, or known engines with which they are incompatible. All of these, and the `MasalaEngine::data_representation_is_incompatible_with_engine(·)` functions implemented by engines, are used by the `MasalaDataRepresentationManager` when fielding requests for data representations through `MasalaDataRepresentationRequest` objects passed by external code to the manager. Like engine requests, data representation requests contain one or more `MasalaDataRepresentationRequestCriterion` objects, defined in namespace `masala::base::managers::engine::data_representation_request`. In addition to requests based on category, name, keyword, or Boolean combinations of any of the above, requests can also be based on compatibility with a particular engine.

#### 8.1.8 Thread, GPU, and MPI management

The `MasalaThreadManager`, definded in namespace `masala::base::managers::threads`, is a static singleton that maintains a CPU thread pool of user-defined size, and fields requests for work to be done in threads. Although this is inspired by the `RosettaThreadManager` that we implemented in Rosetta in 2019, it is a complete rewrite with some efficiency enhancements. By Masala convention, Masala classes or external code should not launch or manage threads on their own. Instead, the workflow is for Masala modules or external code wishing to perform work in parallel to construct an instance of a `MasalaThreadedWorkRequest` (a class defined in the same namespace), load it with a set of independent tasks that can be performed in any order, set a number of threads to request, and pass the `MasalaThreadedWorkRequest` to the `MasalaThreadManager`. The `MasalaThreadManager` will perform the work in at most the number of threads requested, though if fewer threads are available, the work will be carried out in fewer. There is a strong guarantee that at least the thread making the request will be used to do the work. This system permits nested calls for work to be multi-threaded: that is, it is possible for module A to request *N* tasks to be done in *N* threads each of which involves invoking module B, which will also request *N* tasks to be done in *N* threads. Were modules independently launching threads, this could result in a thread explosion, with *N ^r^* threads being launched given *r* levels of recursion. But becuase the thread manager maintains a set thread pool of given size, if the thread pool is saturated, work is performed serially. Unlike the Rosetta thread manager, the `MasalaThreadManager` releases threads back to the thread pool as soon as their work finishes and no new jobs exist for them to do, even if other threads are still executing work from the same `MasalaThreadedWorkRequest`. This permits these threads to be assigned new work from a new `MasalaThreadedWorkRequest`, if one is available, permitting better load balancing.

An example of multi-theaded work is shown in Listing 11. In this example, taken from the `MonteCarloCostFunctionNetworkOptimizer` defined in the Standard Masala Plugins library (see Appendix C, section 9.1.1), we are submitting a request for a user-defined number of Monte Carlo trajectories to be carried out in parallel, to produce a number of solutions to a number of cost function network optimization problems, in a user-defined number of threads. Results are stored in the `solutions by problem` vector, which is passed to the function that is executed in threads as an input parameter. Note that the `MasalaThreadManager::do work in threads(·)` function produces only a summary of the work done as output. Each task in the threaded work request must include storage for its own output. Also note that if the number of tasks is less than the number of threads requested, the `MasalaThreadManager` assigns only the number of threads needed, not the number requested.

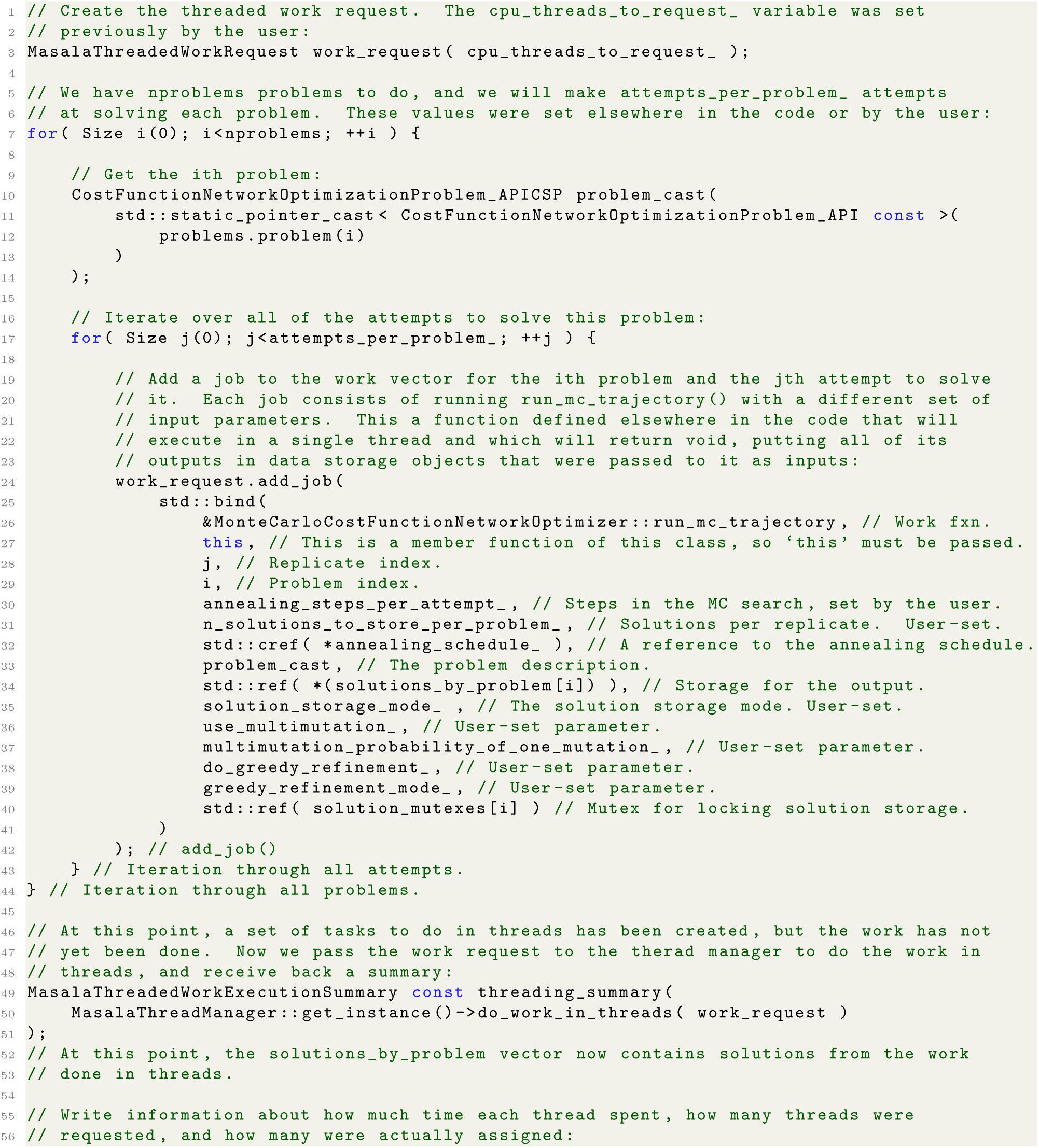
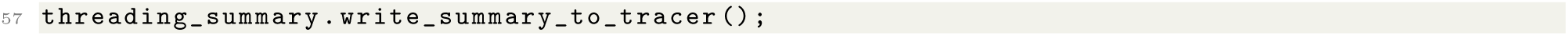

**Listing 11.** An example of work carried out in threads using the `MasalaThreadManager` and a `MasalaThreadedWorkRequest`, taken from the `MonteCarloCostFunctionNetworkOptimizer` in the Standard Masala Plugins library.

The namespace `masala::base::managers::gpu` has been prepared for a similar GPU manager to manage work done on one or more GPUs. Implementation details will be discussed in a future publication focusing on Masala plugin optimizers for GPU.

The namespace `masala::base::managers::mpi` defines the `MasalaMPIManager`. While this is ultimately intended to manage Message Passing Interface (MPI) processes and ensure safe communication during parallel work without deadlock, its full functionality will be described in a future publication once internal MPI job distribution is implemented. Version 1.0 of Masala is only MPI-aware insofar as the `MasalaMPIManager` keeps track of the MPI rank and total number of ranks if external code that links Masala launches MPI processes.

#### 8.1.9 Random number generation

Random numbers are central to many stochastic optimization algorithms used for heteropolymer structure prediction and design. An important lesson from Rosetta was that, for scientific reproducibility, the central management of random number generators and random seeds is essential. Additionally, scientific rigour requires the use of quasi-random number generators producing sufficiently stochastic distributions of generated numbers, to avoid biasing results. In Masala, all random number generation is handled by the `MasalaRandomNumberGenerator`, located in namespace `masala::base::managers::random`. This currently uses the Mersenne Twister algorithm [77] implemented in the C++ Standard Template Library’s `std::mt19937 64` random engine; however, by providing a central location for random number generating code, it is easy to swap this out for more robust or faster random number engines as they are developed. The `MasalaRandomNumberGenerator` assigns a unique generator with a unique random seed to each Masala thread and MPI process. By Masala convention, Masala modules should not make their own calls to other (potentially less scientifically rigorous) random number generators.

#### 8.1.10 Version management

Masala’s Core library and its plugin libraries are separately versioned. Library versions take the form of `<MAJOR version>.<MINOR version>`, where the major and minor versions are nonnegative integers. (Note that version 3.10 follows version 3.9 by this scheme.) The version is set in the `PROJECT` line near the top of the `CMakeLists.txt` file in the `cmake/` subdirectory of each library.

Versioning permits version dependencies. When plugin modules register themselves with the `MasalaPluginManager` (see section 8.1.5), they must provide their own version information, and can optionally provide version requirements for other plugin libraries. Once all plugins are loaded, the `MasalaVersionManager`, located in namespace `masala::base::managers::version` iterates through all version requirements and checks that all are satisfied, throwing an informative error message if they are not. Plugin libraries can specify that other plugin libraries are required or optional, and can specify required version ranges for both required and optional libraries. Listing 12 shows the `register_library()` function (see Section 2.2.1) for the Standard Masala Plugins library (described in Appendix C, section 9). In addition to iterating through all sub-libraries and calling auto-generated code to register their plugin modules with the `MasalaPluginModuleManager`, this function provides information to the `MasalaVersionManager` about the Standard Masala Plugins library’s version, and specifies a minimum version for the Masala Core library.

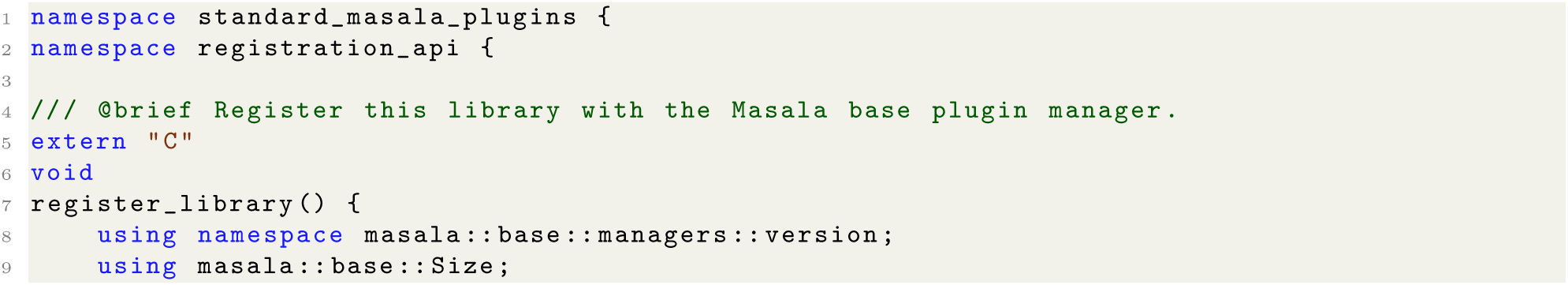
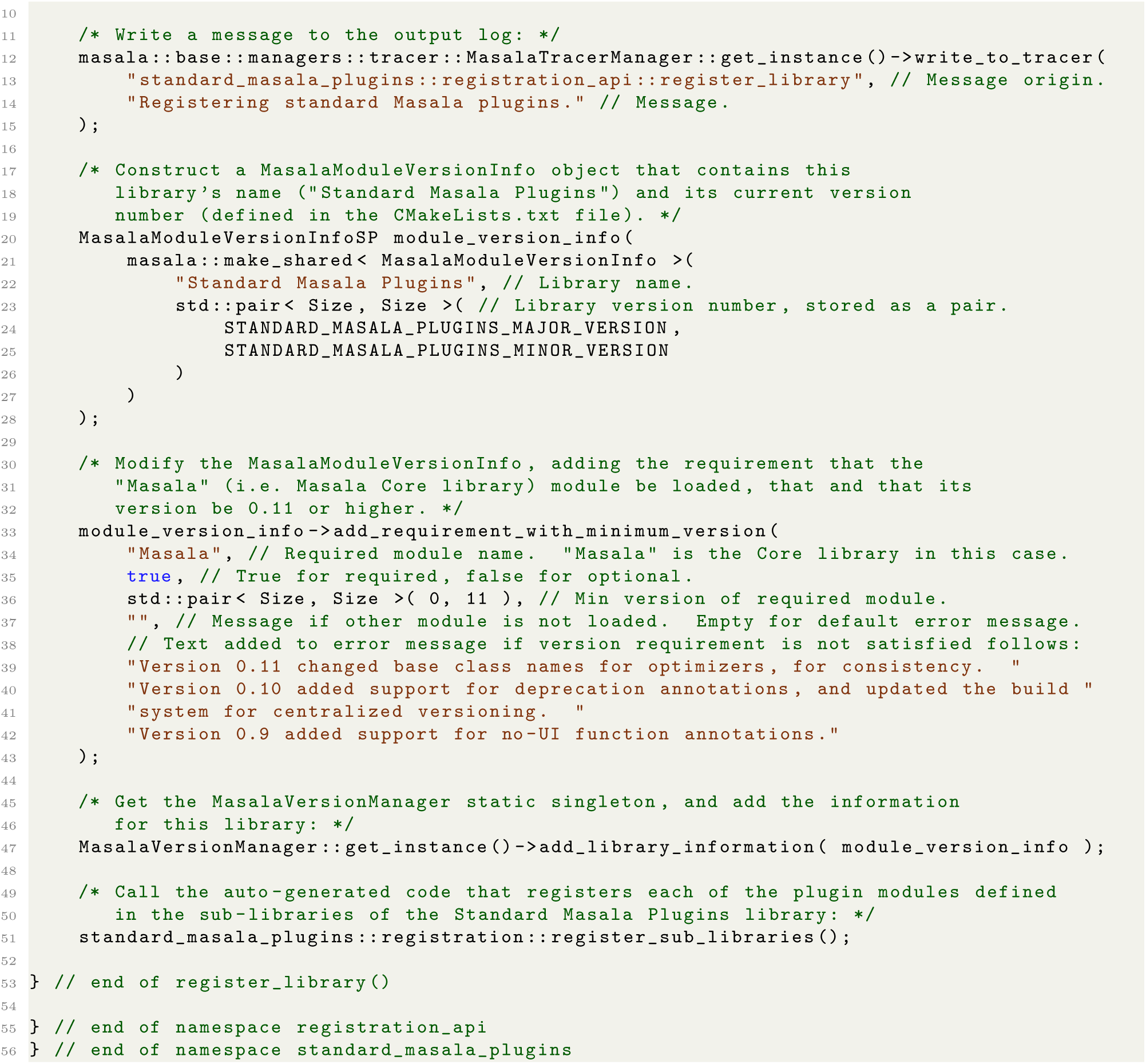

**Listing 12.** The `register_library()` function for the Standard Masala Plugins library, showing specification of version requirements. This code is located in file `register_library.cc`, in subdirectory `src/registration api/` in the Standard Masala Plugins library repository.

#### 8.1.11 Other base sub-library namespaces

The `base` sub-library contains a handful of other namespaces. These include `masala::base::error`, `masala::base::enums`, and `masala::base::utility`.

In the `masala::base::error` namespace, error handling classes such as the `MasalaException` are defined, as well as the useful macros `MASALA_THROW` (which throws a Masala exception), `CHECK_OR_THROW` (which checks that an assertion is true and throws a Masala exception if it is not), and `CHECK_OR_THROW_FOR_CLASS` (which does the same as `CHECK_OR_THROW`, but automatically includes information in the error message about the class and function in which the error was produced). Note that `CHECK_OR_THROW_FOR_CLASS` must not be used from class constructors, since it makes use of the `class_namespace()` and `class_name()` overrides, and virtual function calls cannot be made from a constructor.

Enum classes are short lists of elements known at compilation time, which can be converted to integers by the compiler. They provide a convenient means by which a developer may refer to a descriptive name while the machine code that gets produced by the compiler refers only to integers (which are far more efficient to compare or interpret than strings). Although the general rule in the Masala Core library is for no chemical or biochemical concepts to exist below the level of the `core` sub-library, the `masala::base::enums` namespace is an exception, containing enum classes describing particular broadly-useful chemical concepts. These include the `ChemicalBondType` enum (which enumerates chemical bond types, such as `SINGLE_BOND`, `DOUBLE_BOND`, `TRIPLE_BOND`, `HYDROGEN_BOND`, *etc.*) and the `AtomHybridizationState` enum (which enumerates hybridization states of atoms, such as *s*, *sp*, *sp2*, *etc.*).

The `masala::base::utility` namespace contains useful utility functions for manipulating containers (such as `std::vector` or `std::list`), for editing `std::string` objects, and for measuring time (particularly in benchmarking contexts). It also contains Masala macros for handling execution policies, a C++ language feature that changed between C++17 and C++20.

### 8.2 The numeric and numeric api sub-libraries

Although Masala’s plugin and engine systems are versatile enough to permit plugin libraries to offer numerical engines intended to solve problems of types completely unknown to the Core library, with interfaces accessed entirely through the API definition objects, it is useful for particular privileged problem types to be defined in the Core library, permitting implementation of abstract base classes for their data representations and for their solver engines, with certain interface elements known to calling code at compilation time. This helps to ensure that plugin implementations of the data representations and solver engines for these problem types have common external interfaces, though they may also have additional specialized configuration functions unique to particular derived classes that are accessible only through API definitions.

All optimization problems in Masala derive from a common `OptimizationProblem` (singular) base class, in namespace `masala::numeric::optimization`; this class in turn derives from `MasalaDataRepresentation` (see Section 2.2.2). A container class for collections of optimization problems, `OptimizationProblems` (plural), is defined in the same namespace. Similarly, this namespace defines the `OptimizationSolution` (singular) class, intended to store a single solution to an optimization problem, and the `OptimizationSolutions` (plural) class, which serves as a container for a set of `OptimizationSolution` objects representing multiple solutions to the same problem. Masala optimization engines operate on vectors of optimization problems and yield vectors of solutions in accordance with the SIMD paradigm, though of course it is always possible to provide a vector containing only one problem instance.

It is often the case with optimization problems that, once all data defining the problem have been loaded into the `OptimizationProblem` data representation, there may exist sensible manipulations that can be performed in order to facilitate solving the problem more efficiently. The `OptimizationProblem` class therefore provides a function `finalize()` which calls a virtual `protected_finalize()` function that may be overridden by derived classes. This function provides the opportunity to perform any data manipulation or reorganization (sorting, pre-computation of values, *etc.*) needed for efficiency. (Note that the `finalize()` function handles mutex locking for thread safety. The `protected_finalize()` function is expected to be called from a mutex-locked context, and need not implement any thread safety mechanisms of its own. This is a common pattern followed throughout the Masala codebase.) Once `finalize()` has been called, the problem description is expected to be locked, and further calls to setter functions to input additional data will throw exceptions with error messages.

Auto-generated API classes `OptimizationProblem_API, OptimizationProblems_API, OptimizationSolution_API`, and `OptimizationSolutions_API` are produced by the build system at compilation time, and reside in namespace `masala::numeric_api::auto_generated_api::optimization`. These serve as parent classes for the API containers for particular types of optimization problems, possibly implemented in plugin libraries: as discussed in section 2.2.3, Masala’s build system ensures that auto-generated API container classes have class inheritance patterns matching the classes that they contain. All code in the `masala::numeric_api::auto_generated_api` namespace is generated automatically from API definitions; however, the adjacent `masala::numeric_api::base_classes` namespace contains hand-written code defining additional base classes that are dependent on the problem and solution API classes. It is here, in `masala::numeric api::base_classes::optimization`, that the PluginOptimizer class, which derives from MasalaEngine, is defined. This serves as a common base class for all optimizers defined in plugin libraries.

Version 1.0 of Masala’s `core` library defines base classes, which themselves inherit from `OptimizationProblem`, for two broad classes of optimization problems: discrete cost-function network optimization (CFN) problems, and continuous real-valued local optimization (RVL) problems. These are described further in sections 8.2.1 and 8.2.2, below. Derived classes implementing particular strategies for solving these problems are not implemented in the Core Masala library, but in plugin libraries.

#### 8.2.1 Discrete cost-function network optimizer base classes

Discrete cost function network optimization (CFN) problems have been at the heart of protein and non-canonical heteropolymer design for decades [55, 56, 59]. A cost function network is a set of *N nodes* (though some authors use the term *variables*). The *i^th^* node has a set of *D_i_* discrete *choices* associated with it. We express a candidate selection of one choice per node as a vector 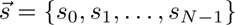, where *s_i_* is an integer in the interval [0*, D_i_ −* 1] representing the index of the choice selected at the *i^th^* node. An optimal selection 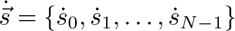 of one choice per node is sought that minimizes an objective function 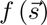. Note: in this chapter, we will use an arrow (*e.g.* 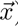) to indicate a vector, a dot (*e.g. ẋ* ) to represent a scalar value that is a global optimum for a one-dimensional function, and a dot and an arrow (*e.g.* 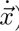) to represent a vector that globally optimizes a multidimensional objective function. In section 8.2.2, we will also introduce *ẍ* and 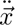 to indicate scalar and vector quantities representing local minima for one- or *N*-dimensional objective functions.

A concrete example of the above is the fixed-backbone rotamer optimization problem in heteropolymer structure prediction and design. In this case, one has *N* positions in a protein or other heteropolymer, with *D_i_* possibilities at the *i^th^* position for the identity and conformation, collectively called the *rotamer*, of the sidechain. The challenge is to produce an optimal selection 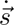 of one rotamer per position, so that an objective function (often based on the physical energy of a heteropolymer given the sidechain identities and conformations) is minimized. When the set of rotamers at each position consists of many different conformations of the sidechain of a single type of chemical building-block, this is a *packing* problem, a sub-problem solved during structure prediction. When the rotamers vary in both sidechain conformation and chemical building-block identity, then the selection of one rotamer per position fixes the identity along with the sidechain conformation, producing the sequence as output, making this a *design* problem. Mixed problems are also frequently encountered, for example in one-sided interface design, in which one may wish to keep the sequence of a target fixed (but permit its sidechain conformations to change) while simultaneously designing the sequence of a binder.

CFN problems like the rotamer optimization problem are NP-hard, growing intractable as either *N* grows large (*e.g.* if one wished to design a very long peptide or other heteropolymer), or as *D* grows large (*e.g.* if one wished to design a peptide or heteropolymer with many possibilities for the building-block identity at each position). Non-canonical peptide design replaces the 20 canonical amino acids with a much-expanded palette of hundreds or thousands of synthetic building-blocks, vastly increasing the difficulty of the optimization problem solved during design. For this reason, fast, clever algorithms are critical, as is having easy means of experimenting with new methods.

Rosetta’s `packer` module, discussed in section 2.1.1, uses a simulated annealing-based search to find the optimal solution 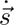, or a near-optimal solution. Unfortunately, the `packer` is hard-coded to operate on shared pointers to Rosetta `Residue` objects, used to represent rotamers, making it difficult to apply to more abstract CFN problems representing other, related types of problems. We have shown previously that the multi-body docking problem can also be mapped to a CFN problem representation [73]. So too can other problems of biological relevance. Additionally, the `packer` was developed with the assumption that the objective function is *pairwise decomposable* at the level of rotamers: that is, 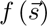 is given by Eq. (4), and is divisible into terms each depending on the choice at a single node (the rotamer at a single position in a peptide or other heteropolymer) and the pair of choices at two nodes (the pair of rotamers at two interacting positions in a peptide or other heteropolymer). While this assumption can be used to accelerate iterative searches for a solution by permitting pre-computation of one- and two-body energies (the various *α_i_* (*s_i_*) and *β_j,k_* (*s_j_, s_k_*) values in Eq. (4)), many design problems are enhanced by augmenting the objective function with non-pairwise *design-centric guidance terms* intended to penalize undesirable features or promote desired features. These, discussed further in sections 10.4 and 10.5 of Appendix D, and reviewed in [42] and [43], were a late addition in Rosetta, and required significant modifications to the `packer`. Unfortunately, while the `packer` offers some ability to sub-class and vary the annealer, it is difficult to experiment with alternative approaches – particularly since most of Rosetta is built atop the `core` libraries in which the `packer` is defined, requiring re-compilation of much of the source code and re-linking of nearly everything whenever a change is made.

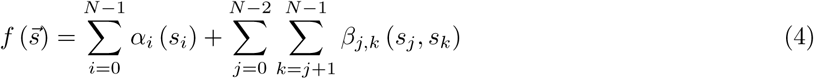

Masala in contrast provides an abstract base class for CFN solvers, the `PluginCostFunctionNetworkOptimizer` class defined in namespace `masala::numeric_api::base_classes::optimization::cost_function_network`. This permits derived classes, using diverse algorithms or even diverse hardware (CPU, GPU, FPGAs, *etc.*), to be written in plugin libraries that can be swapped in and out at runtime. The particular CFN solvers available in the Standard Masala Plugins library are discussed in section 9.1.1 in Appendix C. Some CFN solvers may make use of simulated annealing or other related methods. For convenience, a base class for annealing schedules, `AnnealingScheduleBase`, and a derived class for plugin annealing schedules, `PluginAnnealingSchedule`, are provided in namespaces `masala::numeric::optimization::annealing` and `masala::numeric_api::base_classes::optimization::annealing`, respectively. These define the non-constant pure virtual work function `Real temperature()`, which returns a real temperature value and increments an internal tracker of the point in the annealing trajectory, as well as the constant pure virtual work function `Real temperature( Size time_index )`, which takes a trajectory step as input and produces a temperature as output. These must be implemented by derived classes implementing particular annealing schedules, found in plugin libraries.

Namespace masala::numeric::optimization::cost function network defines a base class for a single CFN problem (CostFunctionNetworkOptimizationProblem, singular), as well as for a collection of problems (CostFunctionNetworkOptimizationProblems, plural). These derive from OptimizationProblem and OptimizationProblems, respectively. This namespace also defines a base class for a single CFN solution (CostFunctionNetworkOptimizationSolution, singular) and one for a collection of solutions to one problem (CostFunctionNetworkOptimizationSolutions, plural), which derive from OptimizationSolution and OptimizationSolutions, respectively. The PluginCostFunctionNetworkOptimizer base class includes a pure virtual work function `run_cost_function_ network_optimizer(·)` that must be implemented by derived classes. This function accepts a `CostFunctionNetworkOptimizationProblems_API` object (a collection of CFN problems, encapsulated in an API container), and produces a vector of `CostFunctionNetworkOptimizationSolutions_APICSP` (a vector of collections of CFN problem solutions, with each collection encapsulated in an API container and handled by constant shared pointer). One CFN solution collection is returned for each problem.

While the `CostFunctionNetworkOptimizationProblem` and `PluginCostFunctionNetworkOptimizer` base classes make no assumptions about the nature of the objective function, for efficiency it can be useful to handle the special case of objective functions that are pairwise-decomposable (*i.e.* which satisfy Eq. (4), above). For this reason, a special class for CFN problems that are pairwise-decomposable, `PluginPair-wisePrecomputedCostFunctionNetworkOptimizationProblem`, is defined in namespace `masala::numeric -api::base_classes::optimization::cost_function_network`. This derives from the `CostFunctionNet-workOptimizationProblem` class, and provides pure virtual functions, `set_onebody_penalty(·)` and `set - twobody penalty(·)`, which must be defined by derived classes, to input the *α_i_*(*s_i_*) and *β_j,k_*(*s_j_, s_k_*) values in Eq. (4). Derived classes may optionally implement a `protected_finalize()` function to reorganize input data into a more efficient form for aiding a particular solver when the `finalize()` function is called.

The `masala::numeric::optimization::cost_function_network` namespace also defines a special class, `CFNProblemScratchSpace`, intended to accelerate repeated calculations by permitting any data needed for fast updates to be cached and rapidly retrieved. This base class defines no setters, getters, or work functions, and is annotated to be a no-API class (i.e. one for which no API container gets auto-generated at compilation time). Derived classes may implement their own interfaces, usable only by the particular solvers or data representations that they are intended to accelerate. One derived `PluginPairwisePrecomputedCFNProblemScratchSpace` class is also provided in `masala::numeric_api::base_classes::optimization::cost_function_network`, to serve as a base class for scratch spaces for pairwise-decomposible CFN problem data representations.

Masala provides support for non-pairwise objective functions in two ways: if the objective function bears no resemblance to Eq. (4), it is always possible to implement a new `CostFunctionNetworkOptimizationProblem` derived class, though it is the developer’s responsibility to provide means of making this efficient to solve. Alternatively, a non-pairwise function 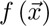 may resemble Eq. (5): it may be a pairwise function augmented with *M* non-pairwise terms 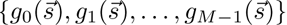. To handle this case, a `PluginCostFunctionNet-workOptimizer` may have additional non-pairwise *cost functions* appended to it. A base class for cost functions, `CostFunction`, is defined in `masala::numeric::optimization::cost_function-network::cost function`, as is a no-API scratch space class, `CostFunctionScratchSpace`. The classes `PluginCostFunction` and `PluginCostFunctionScratchSpace`, which derive from `CostFunction` and `CostFunctionScratchSpace`, respectively, are located in `masala::numeric_api::base_classes::optimization::cost_function_network::cost_function`. These serve as base classes for cost functions (and their scratch space classes) implemented in plugin libraries.

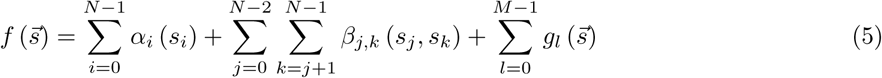

#### 8.2.2 Continuous real-valued local optimizer base classes

Continuous real-valued local optimization (RVL) solvers find a local minimum 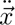 of an arbitrary continuous function 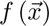 that maps ℛ*^N^ →* ℛ. (Here, we use a double dot to indicate a scalar or vector quantity that locally minimizes an objective function.) RVL solvers may or may not make use of the gradient 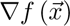 in order to do so; if they do, then the function 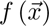 must be at least *C*^1^ continuous, and may need to be *C*^2^ continuous if the solver approximates the inverse of the Hessian matrix from multiple gradient evaluations, as quasi-Newtonian solvers do. Note that RVL problems are the class of problem solved by Rosetta’s `minimizer`, which uses quasi-Newtonian gradient descent methods to find local minima in the conformational energy landscape when relaxing macromolecular structures [1].

In Masala, the base class for the RVL problem data representation is class `RealValuedFunctionLocalOptimizationProblem`, in namespace `masala::numeric::optimization::real_valued_local`, and this derives from the `OptimizationProblem` class in the parent namespace. Common API elements for RVL problem objects include the setters `set_objective_function(·)`, which accepts and stores a `std::function` object that takes as input an *N*-dimensional `Eigen::Vector` and returns a `masala::base::Real` value, `set_objective_function_gradient(·)`, which accepts and stores a `std::function` object that takes as input an *N*-vector coordinate and the address of another *N*-vector, and which populates the second *N*-vector with the gradient while returning the function value at the point, and `add _starting_point(·)`, which accepts and stores an *N*-vector to use as the starting point for iterative searches for a local minimum. A container class `RealValuedFunctionLocalOptimizationProblems` (plural), derived from the `OptimizationProblems` (plural) base class, is also provided.

RVL solvers implemented in plugin libraries derive from the common base class `PluginRealValued-FunctionLocalOptimizer`, in namespace `masala::numeric_api::base classes::optimization::real_- valued_local`. The PluginRealValuedFunctionLocalOptimizer class implements a pure virtual work function `run_real_valued_local optimizer(·)`, which must be implemented by derived classes. This function takes as input a RealValuedFunctionLocalOptimizationProblems API object (an API container for a collection of RVL problems) and produces as output a vector of `RealValuedFunctionLocalOptimization-Solutions_APICSP` objects (a vector of constant shared pointers to API containers for collections of RVL solutions, with one solution collection for each problem).

One special base class is also provided: the `PluginLineOptimizer` class. This is found in namespace `masala::numeric_api::base classes::optimization::real_valued_local`. Line optimizers solve the local optimization problem in one dimension, searching along a vector in *N*-dimensional space for a nearby local minimum. Since many optimization algorithms involve solving line optimization problems as sub-problems, and since there are many algorithms for solving line optimization problems, it makes sense to have a dedicated set of `PluginLineOptimizer` modules that can be mixed-and-matched with larger *N*-dimensional RVL solvers. The `PluginLineOptimizer` class derives from `LineOptimizer`, located in namespace `masala::numeric::optimization::real_valued_local`; this in turn derives from `MasalaEngine`. Note that this means that the `PluginLineOptimzier` class is *not* derived from `PluginOptimizer`. This is by design, since it allows line optimizers to be lightweight classes with unique interfaces that can be invoked with less packaging of problems in problem containers. Each `PluginLineOptimizer` must implement a `run_line_optimizer(·)` work function, which takes as input an objective function 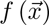 (in the form of a `std::function` object that accepts an *N*-vector 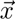 and returns a real number), a starting point 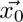 in the search space, the value of 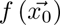, the gradient 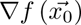, an *N*-vector for a search direction 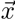, a non-constant reference to a vector 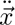 that will be updated by the function to lie in a local minimum 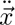 along the search direction 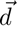, and a non-constant reference to a `masala::base::Real` value in which the value of 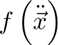 will be stored.

Particular real-valued local optimizers implemented in the Standard Masala Plugins library are discussed in section 9.1.2 in Appendix C.

### 8.3 The core and core api sub-libraries

The `core` sub-library (not to be confused with the Core library, of which the sub-library is a part) contains some of the lowest-level code that defines concepts related to chemistry or biochemistry. It links the `base`, `numeric`, and `numeric_api` sub-libraries, permitting it to directly invoke the classes defined there, though only the `base` and `numeric_api` libraries are guaranteed to have a stable interface. External code may access classes and functions in `core_api` (which is guaranteed to present a stable interface) but not in `core` (which is mutable hand-written code). Namespace `core_api::auto_generated_api` contains automatically generated API classes produced from the API definitions in `core` at compilation time, and the adjacent namespace `core_api::base classes` contains hand-written abstract base classes, which are expected to remain stable from version to version, from which derived classess implemented in plugin libraries may inherit.

#### 8.3.1 Atoms, bonds, molecular geometry, and kinematics

Masala’s `core` sub-library defines many concepts essential for modelling atoms and molecules, and provides convenient classes for representing these concepts. These are intended to provide a stable representation of the components of a molecule as one moves from module to module in a protocol; they are not necessarily built for efficient operations. The expected workflow is that protocols will request appropriate `MasalaEngine` instances for performing particular manipulations of molecular representations, a given `MasalaEngine` will provide a suitable `MasalaDataRepresentation` for the operation, the protocol will copy a subset of the data from the `core` molecular representation into the specialized `MasalaDataRepresentation`, the `MasalaEngine` will operate on the `MasalaDataRepresentation` and produce a result, and data from the result will be copied back to the `core` molecular representation.

##### Atoms and atom iterators

Atoms are the fundamental building blocks for all of chemistry. Namespace `masala::core::chemistry::atoms` defines the `AtomInstance` class, which is used to store all information about a particular atom in a molecule *except* for its spatial coordinates. The information stored includes such things as the formal charge, partial charge, element number, van der Waals radius, and hybridization state (an important concept missing in Rosetta, provided by the `AtomHybridizationState` enum class defined in namespace `masala::base::enums`, as described in section 8.1.11), as well as a constant shared pointer to a record of the element’s full properties. A shared pointer to an `AtomData` object is also provided in this class to allow additional arbitrary data to be attached to an atom. `AtomData` is an abstract base class defined in namespace `masala::core::chemistry::atoms::data`, which derives from `MasalaPlugin` (see section 8.1.5). The `PDBAtomData` class, defined in the same namespace, derives from `AtomData`, and provides storage for information stored from a Protein Data Bank file’s `ATOM` or `HETATM`, such as the atom index in the molecule at read-in, or the PDB atom name. The pure virtual class `PluginAtomData` also derives from `AtomData`, and is defined in namespace `masala::core_api::base classes::chemistry::atoms::data` to serve as a base class for atomic data containers defined in plugin libraries.

The `masala::core::chemistry::atoms` namespace also defines a lightweight iterator class for atoms, the `AtomInstanceConstIterator`. In Rosetta, developer errors frequently arose in molecular container classes when components, like atoms or residues, were primarily accessed by an arbitrarily-assigned integer index, since insertions or deletions would propagate to alter all downstream indices. The `MolecularSystem` class (see section 8.3.2) therefore deliberately omits any means of addressing atoms by an integer atomic index. To avoid such developer errors, atom iterators, which retain their values on insertion or deletion, are the means by which atoms may be accessed.

##### Chemical bonds

While molecules are in reality held together by quantum-mechanical effects arising from the electron clouds surrounding the nuclei, chemical bonds are a convenient fiction commonly used in chemistry [78]. The `ChemicalBondInstance` class, defined in namespace `masala::core::chemistry::bonds`, stores information about the bond type between two `AtomInstance` objects. Bonds are annotated as being of one of several types enumerated in the `ChemicalBondType` enum class, defined in namespace `masala::base::enums` (see section 8.1.11).

##### Molecular geometry

Atoms and chemical bonds are collected in a container called a `MolecularGeometry` object, defined in namespace `masala::core::chemistry`. This is roughly equivalent to Rosetta’s `Conformation` class [1], but with one major difference: where Rosetta’s `Conformation` class is a container for `Residue` objects, representing individual amino acid residues, each of which in turn contains records of individual atoms, the `MolecularGeometry` object is a container for atoms and bonds, as well as atom annotations. Groups of atoms may be annotated as residues, but Masala avoids the indexing headaches that come from ordered arrays of `Residue` objects, in which indices change as residues are inserted and deleted, as well as the problems that arise from the necessity of arbitrarily choosing cutpoints between residues, by not having residues as part of the fundamental organizational hierarchy of molecules. As discussed before, the `MolecularGeometry` object provides no accessors for atoms that are based on integer atomic indices, since indexing can change as sequences change; instead, that `AtomInstanceConstIterator` is the handle by which atoms can be accessed.

Since individual atom records do not store atomic coordinates, the `MolecularGeometry` object itself stores these. This is for efficiency: although the `MolecularGeometry` object is not intended to be a universal representation of molecular data permitting efficient computation in all circumstances (*i.e.* it is expected that internal data will be converted to other data representations before being operated on by high-efficiency engines), the `MolecularGeometry` representation can at least be *fairly* efficient in *many* circumstances by representing atomic coordinates in matrices, on which linear algebra operations (heavily optimized for vectorized execution on modern CPUs or possibly on GPUs) may be performed. To this end, an abstract `AtomCoordinateRepresentation` (deriving from `MasalaDataRepresentation`) and a concrete derived `EigenLinalgCartesianAtom-CoordinateRepresentation` are defined in namespace `masala::core::chemistry::atoms::coordinates`. The latter uses the Eigen linear algebra library [79] to store and permit manipulation of matrices of atom coordinates. The `masala::core_api::base classes::chemistry::atoms::coordinates` namespace also defines a pure virtual `PluginAtomCoordinateRepresentation` class, derived from `AtomCoordinateRepresentation`, to serve as a base class for atom coordinate representations defined in plugin libraries.

The `MolecularGeometry` object stores a master atomic coordinate data representation (currently defaulting to the `EigenLinalgCartesianAtomCoordinateRepresentation`). It may optionally store other representations of atomic coordinates, permitting caching of data for sequential `MasalaEngines`, though ultimately all coordinates will be converted back to the primary representation to carry data forward to later steps of a protocol which may expect data in completely different representations. This also was inspired by lessons learned from Rosetta: in Rosetta, dual Cartesian (*x*, *y*, and *z*) and internal-coordinate (bond length to parent atom, bond angle to grandparent, torsion angle to great-grandparent) representations are used within the `Conformation` object to represent atomic coordinates. Rosetta maintains both in all protocols, regardless which is more efficient for any given step, possibly resulting in some wasted clock cycles for unneeded updates; moreover, Rosetta has no support for alternative representations. In Masala, we wanted to permit both greater efficiency (allowing one representation at a time to be used by any given engine) and greater versatility (allowing new types of representations to be defined by plugin modules).

It is expected that the MolecularGeometry class will expand in future versions of Masala as Masala’s direct structure manipulation abilities are built out further.

##### Atom chain kinematics

As discussed in section 2.1.1, one of Rosetta’s strengths is its kinematic model, based around internal degrees of freedom like bond lengths, bond angles, and torsion angles, rather than on the Cartesian coordinates of atoms. This permits conformational sampling and gradient-descent minimization in torsional space, allowing larger moves in a lower-dimensional space. While powerful, this model is not universally applicable. In some cases, inverse kinematic models borrowed from the robotics field have been useful for closing loops or sampling macrocycle conformations, and these underlie Rosetta’s `GeneralizedKIC` module. Parametric methods, in which conformations are described by generating equations with parameters that may be continuously varied to alter conformation, have also proved powerful [19], but these were a late addition to Rosetta. Since these do not directly work with the kinematic model, propagation of derivatives has not been supported, preventing gradient-descent minimization in parameter space. In addition to lacking the versatility to support alternative kinematic models, Rosetta’s kinematic system is also hard-coded to propagate conformational effects and derivatives using the CPU. Until recently, it did this with recursive code which had the unfortunate effect of limiting the depth of an atom tree to the recursion limit of a user’s system. This was refactored in 2020 to replace the updates with a non-recursive algorithm, albeit one that still unfortunately takes advantage of modern parallel CPU architecture poorly. There is still no support for atom coordinate updates on GPU in Rosetta.

Masala’s Core library implements no kinematic model of its own, but provides base classes for kinematic data representations to be implemented in plugin libraries. The `KinematicDataRepresentationBase` class, defined in namespace `masala::core::molecular system::kinematics`, derives from `MasalaDataRepresentation` (see Section 2.2.2). The build system auto-generates API class `KinematicDataRepresentationbase-api` in `masala::core_api::auto_generated_api::molecular system::kinematics`, from which all API containers for plugin kinematic data representations automatically derive. Plugin kinematic data representations themselves may inherit from `PluginKinematicDataRepresentation`, in namespace `masala::core_api::base - classes::molecular system::kinematics` (which inherits from `KinematicDataRepresentationBase`).

#### 8.3.2 Molecular systems

The `MolecularSystem` class, defined in namespace `masala::core::molecular_system`, is intended to be Masala’s general purpose representation of a molecule or collection of molecules, plus the environment around them. This makes it analogous to Rosetta’s `Pose` class [1], albeit with one important difference: where a valiant effort was made to ensure that `Pose` could be both versatile and performant (in many different contexts), the `MolecularSystem` class is intended primarily for versatility. The `MolecularSystem` class internally stores a `MolecularGeometry` object, and will likely expand to include additional data as needed. Code that demands performance is expected to convert data from a `MolecularSystem` into an appropriate `MasalaDataRepresentation` and to pass it to an appropriate `MasalaEngine`, then copy the result back into the `MolecularSystem`.

#### 8.3.3 Basic Protein Data Bank (PDB) file format support

Although Masala’s system of file interpreters (see section 8.1.6) is ultimately intended to provide almost all functionality for reading and writing particular file formats, we found that it was useful for the Core Masala library to offer basic support for reading Protein Data Bank (PDB) files, to allow testing of the `MolecularSystem` class and related classes without relying on any plugin library. The `BasicPDBReader` class, defined in namespace `masala::core::io`, permits read-in of atomic coordinates and atom types from an ASCII PDB file. It offers no support for advanced PDB functionality, such as `CONECT` records, `REMARK` lines, crystallographic information like `CRYST1` records, *etc.*, and will ignore any lines that it cannot parse. At present, only `ATOM` and `HETATM` lines are understood.

It should be stressed that the `BasicPDBReader` is intended for unit testing only. It should not be relied upon for production protocols. Full support for PDB, mmCIF, MOL2, and other common molecular formats is planned for file interpreters found in future releases of the Standard Masala Plugins library.

#### 8.3.4 Selections and selectors

A selection is any collection of “things” chosen from a larger set, and is a concept that arises many times in molecular modelling when configuring protocols: one may wish to select atoms, or bonds, or residues, or chains, for various purposes. Masala defines a `Selection` base class, derived from `MasalaPlugin` (see section 8.1.5), in namespace `masala::core::selection`. The derived class `AtomSelection` (namespace namespace `masala::core::selection::atom_selectiom`) is a concrete selection of atoms in a `MolecularSystem`.

In addition to having selections, it is necessary to have a means of generating them. The `ResidueSelector` class and its derived classes were a late addition to Rosetta [2], and prior to their introduction, the problem of permitting the user to select parts of a molecule when configuring some `Mover` that would operate on a subset of a structure was solved inconsistently, either by requiring users to generate tedious lists of residue indices or atom names as inputs to Rosetta protocols, or using rules-based selection modules intended for very different purposes. Learning from this, Masala defines a `Selector` base class, derived from `MasalaPlugin`, in namespace `masala::core_api::base classes::selectors`. A `Selector` operates on a `MolecularSystem`, applying some rule to produce a `Selection`. Derived from this are selector classes for selecting particular things, such as `AtomSelector` (in namespace `masala::core_api::base classes::selectors::atom selectors`), which produces an `AtomSelection`. Particular selectors with their own rules are defined in plugin libraries. See, for example, the `ElementTypeAtomSelector` described in section 9.3 in Appendix C.

#### 8.3.5 Scoring

Many molecular modelling protocols involve performing some sort of analysis on a structure or group of structures and producing a score. Commonly, this score represents a physical energy, such as an estimate of the free energy change on folding, an estimated binding free energy change, or a simpler quantity like the enthalpy of a salt bridge, hydrogen bond, or *π*-*π* stacking interaction. All of this is collectively grouped under the concept of *scoring*.

The Core Masala library defines the `ScoringTerm` base class for all modules that do some sort of analysis producing a score. This is found in namespace `masala::core::scoring`. While scoring terms are primarily intended to operate on a `MolecularSystem`, they are intended to be general. For this reason, generic base classes `ScoringTermAdditionalInput` and `ScoringTermAdditionalOutput` are provided in the same namespace, allowing scoring terms to take into acount additional information provided in additional inputs, or to produce outputs beyond a simple real value. Because inputs to scoring terms are expected to be constant, this namespace also defines a `ScoringTermCache` base class, permitting information from one round of scoring to be cached to facilitate or accelerate subsequent rounds of scoring.

Because the masala::core namespace lies outside of the Masala Core library’s API, the derived classes PluginScoringTerm, PluginScoringTermAdditionalInput, PluginScoringTermAdditionalOutput, and PluginScoringTermCache are found in the masala::core api::base classes::scoring namespace. The PluginScoringTerm class defines a pure virtual constant work function score derived(·) with the function signature shown in Listing 13. This must be defined by plugin scoring terms. Called by the non-virtual function score(·), which handles all mutex locking and thread safety, implementations of this function should assume that they are being called from a mutex-locked context, and do not need to implement any thread safety features themselves. This function accepts a constant vector of MolecularSystem objects (handled by constant shared pointer to API containers), plus pointers (which may be nullptr if no input is provided) to vectors of additional inputs, scoring term caches, or additional outputs. By accepting vectors of work rather than individual molecular structures to operate on one by one, the potential for highly efficient vectorized operations by the implementations of these classes is unlocked. Note that all scoring terms derived from these classes are expected to be implemented in plugin libraries, not in the Core Masala library.

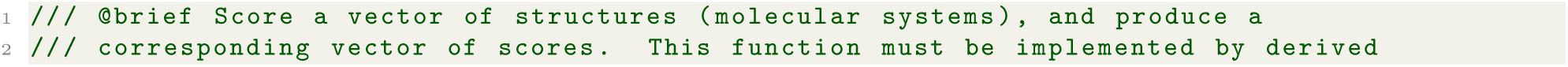
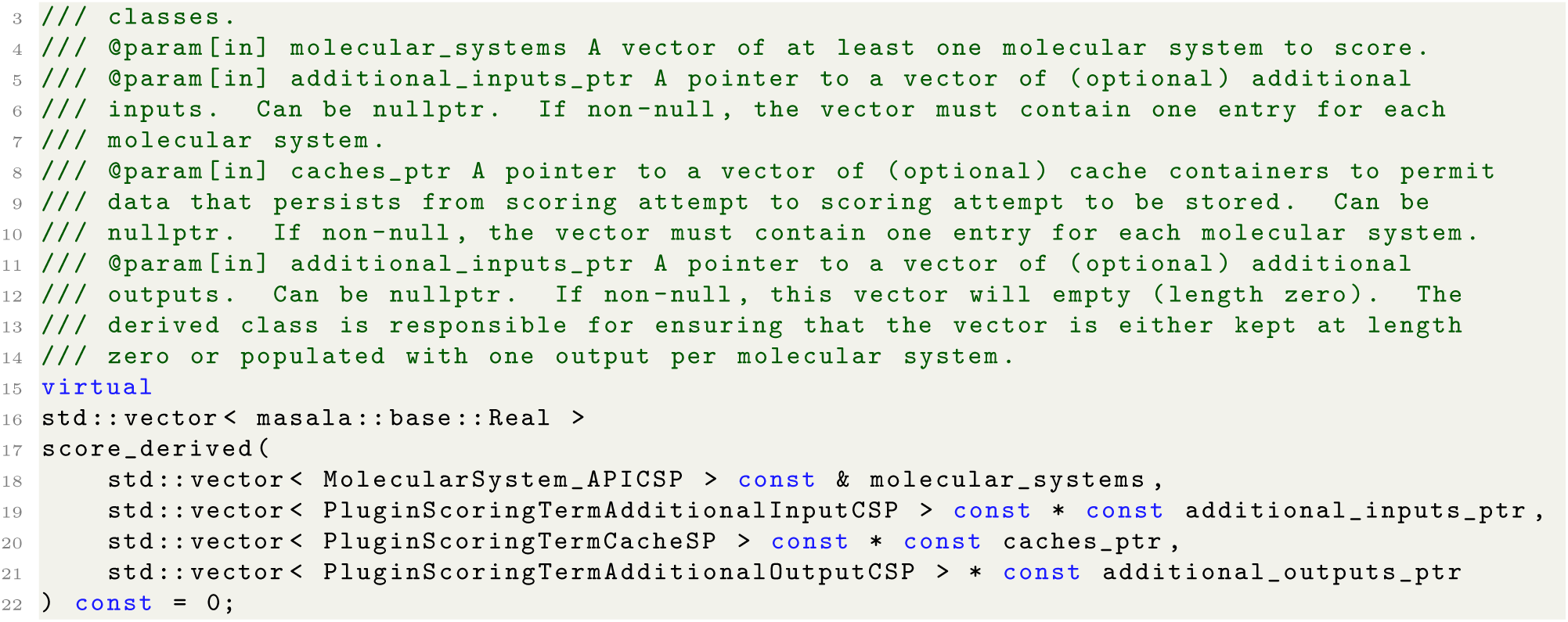

**Listing 13.** The function signature for the `PluginScoringTerm::score derived(·)` pure virtual constant work function, which must be defined by derived plugin scoring terms.

Scoring terms could conceivably depend on fewer or more inputs: in Rosetta, for example, there exist one-body scoring terms (dependent on a single amino acid residue), two-body scoring terms (dependent on pairs of interacting residues), and whole-body energies (dependent on an entire Rosetta `Pose` or molecular structure). Masala currently defines a pure virtual class that derives from `PluginScoringTerm`, called `PluginWholeMolecularSystemScoringTerm`, intended to serve as a base class for scoring terms that are dependent on an entire `MolecularSystem` object. Future extension may add additional types of scoring terms.

### 8.4 Summary: the Core Masala library

The overall goal of the Core Masala library has not been to provide a comprehensive set of tools for macromolecular modelling, but to provide infrastructure for permitting those tools to be implemented in plugin libraries and to be made available to external code that links the Core library. The `base` library defines the most fundamental base classes, the API and plugin system, and the managers of resources, plugin modules, engines, data representations, file interpreters, versions, I/O, threads, and so forth. The `numeric` library has abstract base classes for types of optimizers and data representations defined in other libraries, and the `core` library has classes defining broad chemical concepts (atoms, bonds, molecular geometry, molecular systems), as well as abstract base classes for modules that might operate on molecular systems (such as the scoring base classes). External code should only interface directly with the `base`, `numeric_api`, and `core_api` sub-libraries, since these provide stable APIs.

The framework described here provides a means of developing methods that are both versatile and efficient, while ensuring that most of the effort of structuring the software infrastructure is automated, so that scientists can focus on developing scientific methods implemented in derived classes in plugin libraries. While the Core library may expand in the future, it is anticipated that the framework implemented here should change minimally from version to version, with most development occurring in plugin libraries. The next section describes the Standard Masala Plugins library, version 1.0, in which Masala’s current functionality (which, at the time of this writing, is being used to design peptides, proteins, and peptoids) is implemented.

## 9 Appendix C Organization of the Standard Masala Plugins library, version 1.0

As discussed in Appendix B (section 8), Masala’s Core library is intended to define software infrastructure and base classes, but *not* to provide particular implementations of engines or data representations. The

Standard Masala Plugins library is intended to be a Masala plugin library featuring a general, broadly-useful toolkit of optimization engines, the associated data representations, and other modules. As the “standard” part of the name suggests, it is anticipated that this will be a plugin library that most users will choose to load most of the time. Modules in this library are optimized for execution on multi-core CPUs. GPU versions of these modules are in development, and will be released as a separate plugin library.

Version 1.0 of the Standard Masala Plugins library has five manually-written sub-libraries, with five corresponding auto-generated API sub-libraries. The `optimizers` and `optimizers api` sub-libraries, described in section 9.1, provide the most extensive functionality, featuring high-performance optimization engines for solving cost function network optimization (CFN) problems and real-valued local optimization (RVL) problems. The `file_interpreters` and `file_interpreters api` sub-libraries, described in section 9.2, currently contain plugin modules for reading data formats associated with these optimization problems, though in the future we anticipate expanding this with support for common molecular file formats. The `selectors` and `selectors api` sub-libraries, and the `scoring` and `scoring api` sub-libraries, described in sections 9.3 and 9.4, respectively, define modules for selecting atoms and for computing scores, respectively. And the `registration` and `registration api` sub-libraries, described in section 9.5, contain all code needed to register the Standard Masala Plugins plugin library with the `MasalaPluginLibraryManager`, and ensure that creators for all plugin modules end up registered with the `MasalaPluginModuleManager`, the `MasalaFileInterpreterManager`, the `MasalaEngineManager`, and/or the `MasalaDataRepresentationManager`, as appropriate.

### 9.1 The optimizers and optimizers_api sub-libraries

The `optimizers` sub-library (namespace `standard masala plugins::optimizers` provides hand-written implementations of particular solver engines for optimization problems, and the `optimizers api` sub-library (namespace `standard masala plugins::optimizers api`) contains the auto-generated API containers for these engines. In version 1.0 of the Standard Masala Plugins library, engines for solving CFN optimization problems and RVL optimization problems are provided, and are described in sections 9.1.1 and 9.1.2, below.

#### 9.1.1 Discrete cost function network optimizers

Version 1.0 of the Standard Masala Plugins library implements three types of solvers for cost function network (CFN) problems in the namespace `standard_masala_plugins::optimizers::cost_function_network`, along with an associated data representation. The adjacent namespace `standard masala plugins::optimizers::annealing` also implements five types of annealing schedule for use with those CFN optimizers that perform temperature ramps as part of an iterative search.

##### Data representations for CPU-based CFN solvers

The optimizers sub-library (namespace `standard_masala_plugins::optimizers::cost_function_network`) defines a `PairwisePrecomputedCostFunctionNetworkOptimizationProblem` data representation, which derives from PluginPairwisePrecomputed-CostFunctionNetworkOptimizationProblem in the Core Masala library. This is a general-purpose data representation for pairwise-decomposable CFN problems in which all one- and two-body penalties are pre-computed, with support for augmentation with non-pairwise cost functions (Eq. (5)). Although the CFN solver engines implemented in the Standard Masala Plugins library can operate on other data representations, this data representation is the preferred representation for these solvers, and is likely to provide optimal performance.

The `standard_masala plugins::optimizers::cost_function_network::cost_function` namespace defines a number of non-pairwise cost functions that can be appended to a pairwise-decomposable CFN problem. These derive from `PluginCostFunction` in the Core Masala library. Among other possible applications, these are useful for design-centric guidance terms. While we will not exhaustively explore these here, these include the `GraphBasedCostFunction` and its related classes for efficient computation of the Rosetta `hbnet` design-centric guidance term, discussed further in section 10.4 of Appendix D, as well as other cost functions for other design-centric terms discussed in section 10.5 of the same appendix.

##### Solving CFN problems by greedy descent to find local minima

The first CFN solver implemented in namespace `standard_masala_plugins::optimizers::cost function network` is the `GreedyCostFunctionNetworkOptimizer`. Starting from some staring point 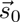, this checks every possible single-node choice substitution for its effect on the value of 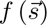. The optimizer then commits the substitution that results in the biggest decrease, and repeats the process over and over until no move can be made that further decreases the objective function. Note that the solution 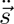 that this provides represents a local minimum in the choice assignment landscape, not necessarily the global minimum.

Obviously, the choice of the starting point 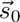 greatly influences the success of this method. The `GreedyCostFunctionNetworkOptimizer` respects candidate starting points provided in the `CostFunction-NetworkOptimizationProblem` instances that it receives as input; alternatively, it can start from a random starting point, though this is not expected to produce a solution close to the global optimum for any but the smallest problems. Generally, greedy descent is a useful refinement technique for improving solutions found by other methods, but does poorly as a global search strategy on its own.

##### Exploring CFN solutions by simulated annealing to seek the global minimum

The second CFN solver implemented in namespace `standard_masala_plugins::optimizers::cost_function network` is the `MonteCarloCostFunctionNetworkOptimizer`. This most closely emulates the behaviour of Rosetta’s `packer`, carrying out simulated annealing-based searches of node choice selections *s⃗* to find those minimizing *f* (*s⃗*). The `MonteCarloCostFunctionNetworkOptimizer` differs from the Rostta `packer` in six key ways, however. First, by operating on abstract CFN problems consisting of nodes and choices, rather than concrete rotamer optimization problems, it can be applied to problems beyond sidechain conformational prediction or sequence design. Second, where the `packer` makes moves consisting of picking a single position at random and assigning a single new, randomly-selected rotamer at random (*i.e.* making a random choice substitution at a random node), the moves that the `MonteCarloCostFunctionNetworkOptimizer` makes by default involve choosing *n* nodes, where *n* is sampled from a Poisson distribution, and substituting random choices at each of those *n* nodes. The Poisson distribution is biased most heavily towards making a single node substitution at a time, but the possibility of making more means that it is easier to escape from local minima in the choice assignment landscape, particularly when the escape may require concerted substitutions at two or three positions. Third, where the `packer` uses a hard-coded stair-stepped annealing schedule that approximates a logarithmic decay, the `MonteCarloCostFunctionNetworkOptimizer` accepts an abstract `PluginAnnealingSchedule` object that defines a custom annealing schedule. Fourth, the `MonteCarloCostFunctionNetworkOptimizer` uses the Metropolis criterion to accept moves (Eq. (1)) rather than the modified Metropolis criterion (Eq. (2)) used by Rosetta’s `packer`. This avoids assumptions about the significance of the value of zero for `f` (*s⃗*), making the `MonteCarloCostFunctionNetworkOptimizer` more general. Fifth, where the `packer` is a strictly single-threaded algorithm, the `MonteCarloCostFunctionNetworkOptimizer` is efficiently multi-threaded (see performance benchmarks in section 4.2), permitting it to perform a separate simulated annealing trajectory in each available thread to allow more regions of choice assignment space to be searched simultaneously, or to solve many different problems at once. And sixth, where the Rosetta `packer` returns only one solution (the lowest-scoring solution encountered during the search), the `MonteCarloCostFunctionNetworkOptimizer` returns a user-defined number of solutions, choosing the lowest-scoring *n* solutions from all simulated annealing trajectories. This can be a more efficient means of generating diversity.

Solutions produced by the `MonteCarloCostFunctionNetworkOptimizer` may be further refined by the `GreedyCostFunctionNetworkOptimizer`. Since both classes are defined in the same sub-library, they are aware of one another, and the former may directly invoke the latter without going through the Masala engine manager. The `MonteCarloCostFunctionNetworkOptimizer` API functions include setters that configure greedy descent-based refinement, with options for performing greedy descent on all solutions found or on the top solutions found. Optionally, the sub-optimal solutions (prior to greedy descent) may also be included in the set of solutions returned.

##### Finding random CFN problem solutions as a control

The third CFN solver implemented in namespace `standard_masala_plugins::optimizers::cost_function_network` is the `RandomCostFunctionNetworkOptimizer`. This optimizer produces one or more completely random solutions to a CFN problem. This is not intended for production runs; rather, it is a useful negative control when benchmarking a new CFN optimization strategy, since, at a minimum, a “good” CFN-solving algorithm should find low-scoring solutions faster than random sampling.

##### Annealing schedules

Five annealing schedules, for use with the `MonteCarloCostFunctionNetworkOptimizer` or any other modules that perform simulated annealing searches, are implemented in the `optimizers` sub-library, in namespace `standard masala plugins::optimizers::annealing`. All of these derive from `PluginAnnealingSchedule`. Since these are Masala plugin modules, these are usable outside of the Standard Masala Plugins library.

The first of these is the `ConstantAnnealingSchedule`. As the name implies, this returns a constant temperature, meaning that it results in a constant-temperature Monte Carlo search. This can be useful for physics-based simulations, in which the goal is to produce a thermodynamic distribution of states at a particular temperature. It is generally not recommended for optimization purposes, however, since a fixed-temperature search cannot alternate between periods of “exploration” (moving up out of local minima to find new parts of the landscape to explore) and “exploitation” (drilling down into local minima), and will find high-quality solutions more slowly.

The second and third annealing schedules are the `LinearAnnealingSchedule` and `LinearRepeatAnnealingSchedule`. The former provides a linear ramp-down from a user-set starting temperature to a user-set ending temperature. The latter carries out a user-defined number of iterations of this, with sharp temperature jumps from the ending temperature to the starting temperature from iteration to iteration, resulting in a saw-toothed temperature graph over time.

The fourth and fifth annealing schedules are the `LogarithmicAnnealingSchedule` and the `LogarithmicRepeatAnnealingSchedule`, which behave like the `LinearAnnealingSchedule` and `LinearRepeatAnnealingSchedule`, respectively, but with the temperature changing linearly in logarithmic space. The `LogarithmicRepeatAnnealingSchedule` provides the behaviour closest to the Rosetta `packer`’s stair-stepped annealing schedule.

Additional annealing schedules may be implemented independently in other plugin libraries and used seamlessly with the MonteCarloCostFunctionNetworkOptimizer, allowing easy experimentation with new approaches.

#### 9.1.2 Real-valued local optimizers

As discussed in section 8.2.2 of Appendix B, real-valued local optimization (RVL) solvers are Masala optimization engines which, given a starting point 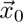 and a continuous real-valued function 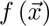 that maps ℛ*^N^ →* ℛ, seek a nearby local optimum 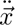 that locally minimizes 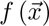. RVL solvers come in two flavours: gradient-based and gradient-free. All RVL solvers derive from `masala::numeric_api::base - classes::optimization::real valued local::PluginRealValuedFunctionLocalOptimizer`.

##### Gradient-based RVL optimizers

Gradient-based RVL optimizers, defined in namespace `standard - masala plugins::optimizers::gradient based`, use the gradient 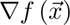 in order to march downhill from 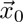 to a local minimum 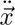. These assume that the function is at least *C*^1^ continuous, and possibly (if an optimizer approximates the Hessian based on successive gradient evaluations) *C*^2^ continuous. Version 1.0 of the Standard Masala Plugins library implements the `GradientDescentFunctionOptimizer`, the `DFPFunctionOptimizer`, and the `BFGSFunctionOptimizer`. The `GradientDescentFunctionOptimizer` performs simple gradient descent by computing the gradient, then carrying out a line search along the gradient direction. On line search convergence, gradient evaluation is repeated until a local minimum is reached.

The `DFPFunctionOptimizer` and the `BFGSFunctionOptimizer`, defined in namespace `standard_masala_- plugins::optimizers::gradient based::quasi_newtonian`, are both quasi-Newtonian RVL optimizers deriving from the common non-instantiable base class `QuasiNewtonianFunctionOptimizerBase` defined in namespace `standard_masala_plugins::optimizers::gradient based`. Both approximate the inverse Hessian matrix from successive gradient evaluations, permitting faster convergence than the `GradientDescentFunctionOptimizer` for many functions. The `DFPFunctionOptimizer` uses the Davidon-Fletcher-Powell approximation scheme [80, 81] shown in Eq. (6):

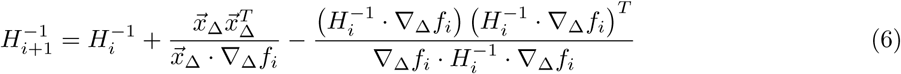

In the above, *H^−^*^1^ is the approximation of the inverse Hessian matrix at the *i*^th^ gradient evaluation step. The vector 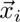 is the coordinate at gradient evaluation step *i*, 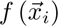 is the function being minimized evaluated at this coordinate, 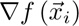 is the function gradient evaluated at this coordinate, 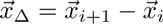 is the difference vector between the *i* + 1^st^ and *i*^th^ coordinate, and 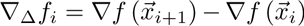 is the difference between successive gradients. The `BFGSFunctionOptimizer` uses the Broyden-Fletcher-Goldfarb-Shanno variant [81, 82] of this update rule shown in Eq. (7)

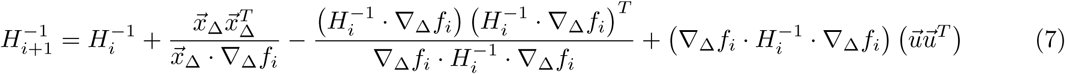

In Eq. (7), *u⃗* is given by Eq. (8).

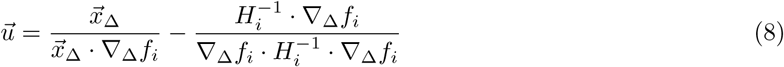

Note that Rosetta’s `minimizer` by default uses the low-memory variant of the BFGS algorithm (L-BFGS) [83], though it also implements standard BFGS (incorrectly named “dfpmin”). There is no implementation of the DFP algorithm in Rosetta.

Since line searches are a typical sub-problem solved repeatedly during a gradient-based search for a local minimum, the Standard Masala Plugins library includes various sub-classes of the `masala::numeric - api::base classes::optimization::real_valued_local::PluginLineOptimizer` class. Line optimizers are similar to RVL solvers, but operate on a function *g* (*y*) that maps ℛ → ℛ. They start from a starting value *y*_0_ and find a new value *y^′^ < y*_0_. While some line optimizers, such as the `BrentAlgorithmLineOptimizer` implementing Brent’s method for line searches [81,84], ensure that *y^′^* is *ÿ*, a local minimum along the line, this is not *necessarily* the case for all line optimizers. The `ArmijoInexactLineOptimizer`, for instance, uses the Armijo-Goldstein convergence criterion [81, 85] that prioritizes speed over accuracy, only guaranteeing some minimum amount of decrease in the objective function; often this is the most efficient approach given iterated line searches when performing multidimensional gradient descent. Line optimizers are implemented alongside gradient-based RVL solvers in namespace `standard masala plugins::optimizers::gradient based`.

##### Gradient-free RVL optimizers

In some cases, it may not be straightforward to compute the gradient 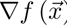, or the gradient may not be continuous or smooth. It is therefore convenient to have RVL solvers that are based solely on 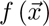, and which only assume continuity of 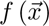 itself and not of its derivatives. Namespace `standard masala plugins::optimizers::gradient free` implements such RVL solvers, such as the `SimplexFunctionOptimizer`, which uses the Nelder-Mead simplex method [81, 86] to search for a local minimum 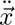 from a starting point 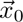. The greater versatility of these gradient-free methods often comes at the expense of efficiency: gradient information can accelerate the search for a local minimum.

### 9.2 The file_interpreters and file_interpreters_api sub-libraries

Implementations of particular file interpreters are found in the `file_interpreters` sub-library, in namespace `standard_masala_plugins::file_interpreters`. Auto-generated API containers for these file interpreters are found in the `file interpreters api` sub-library, in namespace `standard_masala_plugins::file interpreters api`. Version 1.0 of the Standard Masala Plugins library provides file interpreters for two file formats: the ASCII and binary cost function network optimization problem formats that are written by Rosetta’s `InteractionGraphSummaryMetric` module, and read by Rosetta’s `ExternalPackerResultLoader`. While Rosetta can be linked directly against Masala to test Masala optimizers in Rosetta protocols, it can also be useful to store problem descriptions on disk in a format that both Rosetta and Masala can read and write, providing more versatility in benchmarking and experimentation. The `ASCIICostFunction-NetworkProblemRosettaFileInterpreter` reads and writes Rosetta-compatible ASCII representations of CFN problems, and the `BinaryCostFunctionNetworkProblemRosettaFileInterpreter` reads and writes the binary versions. Both are derived from `MasalaFileInterpreter`, defined in the Core Masala library (see section 8.1.6 in Appendix B), and both are implemented in namespace `standard_masala_plugins::file - interpreters::cost_function network`.

### 9.3 The selectors and selectors api sub-libraries

The `standard_masala_plugins::selectors` namespace defines plugin modules derived from subclasses of `Selector` (see section 8.3.4 in Appendix B). A simple example is the `ElementTypeAtomSelector`, a class derived from `AtomSelector` that is defined in namespace `standard_masala_plugins::selectors::atom - selectors`. This selector accepts a `MolecularSystem` and selects all atoms of a given element type specified by the user.

The `standard_masala_plugins::selectors` namespace is expected to grow in future releases of the Stanard Masala Plugins library as Masala gains more plugin modules that directly operate on molecular representations rather than abstract numerical data.

### 9.4 The scoring and scoring api sub-libraries

The `scoring` sub-library is intended to implement scoring terms and associated helper classes. The namespace `standard_masala_plugins::scoring::scoring_terms` contains scoring terms, with the sub-namespace `standard_masala_plugins::scoring::scoring_terms::molecular system` containing terms whose evaluation is dependent on a whole molecular system. These are derived from `PluginWholeMolecularSystem-ScoringTerm` in the Core Masala library (see Appendix B, section 8.3.5). Here, the `ConstantScoringTerm` serves as the simplest possible example of a scoring term. This scoring term simply returns a constant value, set with the API setter function `set_constant_value(·)`.

Future versions of the Standard Masala Plugins library will significantly expand the scoring terms implemented. However, the present infrastructure provides easy means of adding new terms to an energy function, like the Rosetta energy function.

### 9.5 The registration and registration api sub-libraries

All Masala plugin libraries must implement `registration` and `registration_api` sub-libraries. These are needed for interaction with the `MasalaPluginLibraryManager` and registration with the `MasalaPluginModuleManager` (see section 8.1.5 in Appendix B). The `registration` sub-library contains only one important hand-written component: the `register_sub_libraries.cc` file, implementing utility functions `register - sub libraries()` and `unregister_sub_libraries()`. These call auto-generated functions in each of the other sub-libraries, as shown in Listing 14.

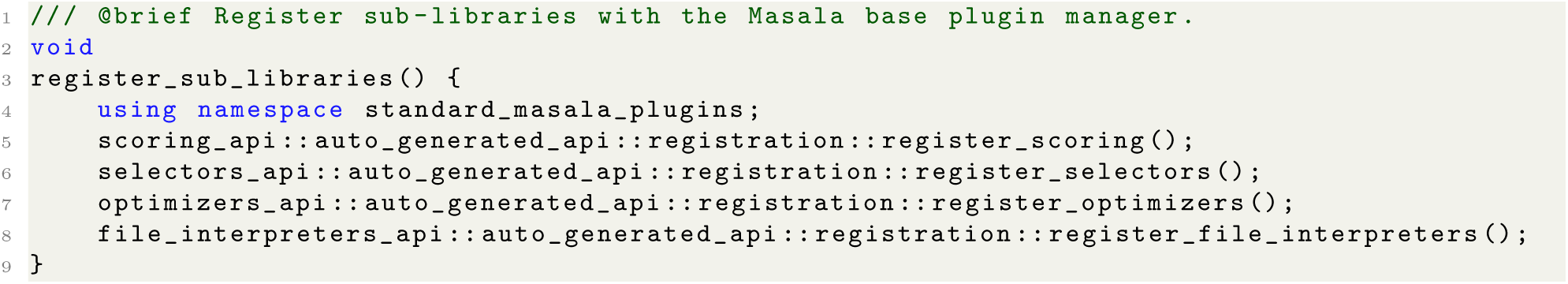

**Listing 14.** The Standard Masala Plugins library’s manually-written `register sub libraries()` utility function, which calls auto-generated code in each of the other sub-libraries to register plugin modules defined in those sub-libraries with the `MasalaPluginModuleManager`. If an additional sub-library were added to the Standard Masala Plugins library, a call to its auto-generated registration function would need to be added here by hand.

The purpose of these function calls is to permit plugin modules in each of these other sub-libraries to be registered with the `MasalaPluginModuleManager`. The auto-generated functions that are called pass to the `MasalaPluginModuleManager` a vector containing one creator for each plugin module in the sub-library. For example, the `register_optimizers()` function looks something like what is shown in Listing 15:

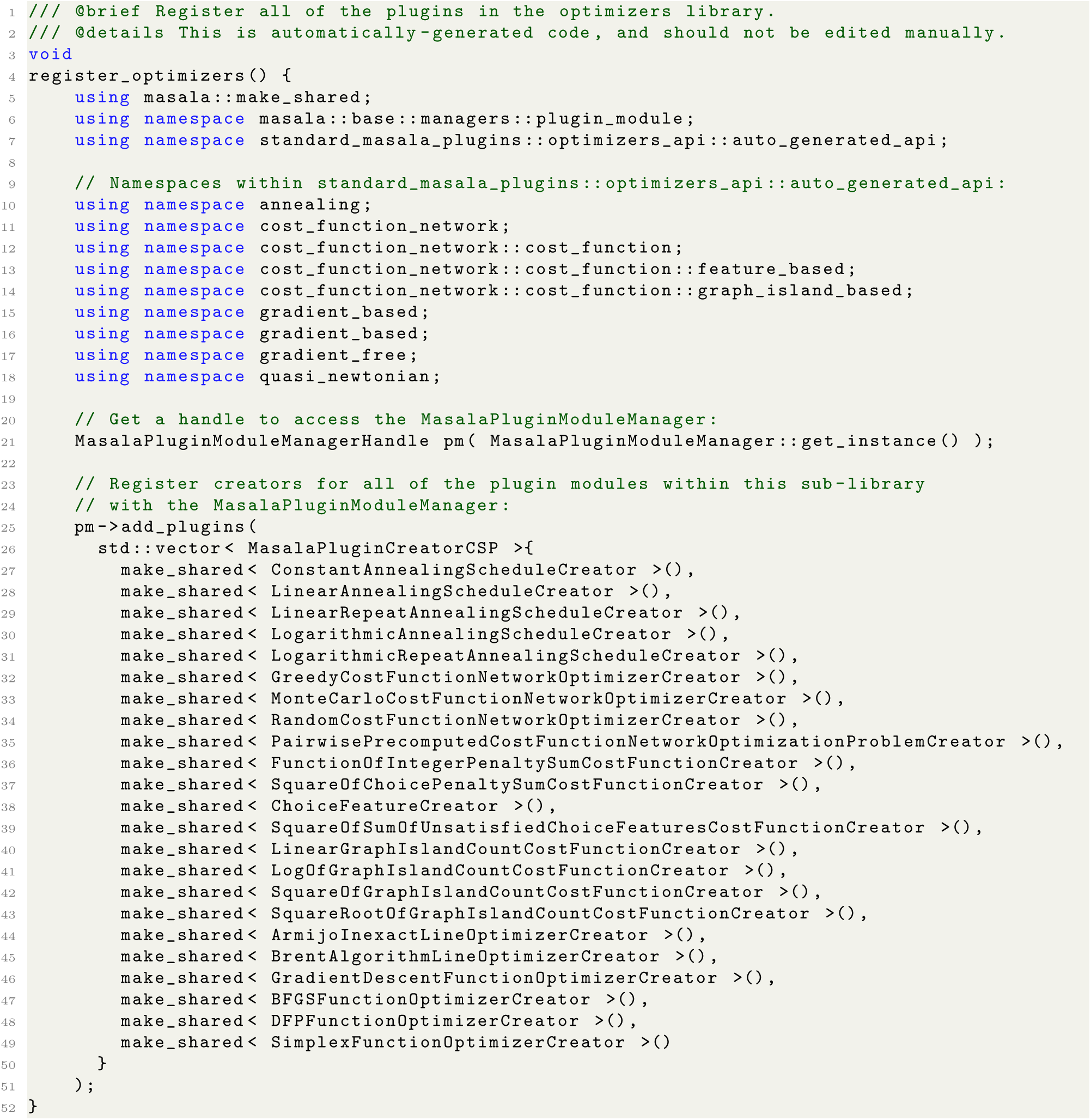

**Listing 15.** The Standard Masala Plugins library’s automatically-generated `register_optimizers()` function, which constructs a vector of creator objects, one for each plugin class in the `optimizers` sub-library, and passes this to the `MasalaPluginModuleManager`. This list is automatically updated by the Masala build system as developers add plugin modules.

The `registration_api` sub-library, unlike most API sub-libraries, contains no automatically-generated code. The only contents are the manually-written `register_library.cc` and `register_library.hh` files, containing the `extern "C" void register_library()` function shown previously in Listing 12 (see Appendix B, section 8.1.10). In addition to setting version requirements for the Standard Masala Plugins library, this function calls `standard_masala -plugins::registration::register sub libraries()` (Listing 14).

### 9.6 Summary: the Standard Masala Plugins library

The primary functionality implemented in version 1.0 of the Standard Masala Plugins library is found in the `optimizers` sub-library (section 9.1), which provides high-efficiency CPU-based optimizers for discrete cost function network optimization (CFN) problems and for continuous real-valued local optimization (RVL) problems. These correspond to the problem types solved by the Rosetta `packer` and `minimizer`, respectively. The integration with Rosetta that allows Masala’s optimizers to be used in place of Rosetta’s optimizers is described in Section 3, and in Appendix D (section 10), below. The Standard Masala Plugins library serves as an accumulator of high-performance optimization methods, permitting, thanks to the versatile plugin infrastructure, experimentation with different approaches within a common context.

The `file_interpreters`, `scoring`, and `selectors` sub-libraries all implement some basic functionality at present, but are intended to be extended further in the future. As described in section 5, Masala’s future roadmap focuses on developing greater ability to manipulate molecular structures from within Masala code, and in Masala 2.0 and 3.0, development efforts will press forward toward Masala as standalone software. By focusing on Masala’s optimizers for version 1.0, we have first ensured that it can serve as a useful extension library for existing software.

## 10 Appendix D Using Masala modules from external code

This appendix describes modifications made to Rosetta to allow Masala modules loaded at runtime to be configured by the user through Rosetta’s user interfaces and used in place of Rosetta’s internal modules. In section 10.1, we describe modifications to Rosetta’s initialization subroutines to permit Masala plugin libraries to be loaded. Section 10.2 describes Rosetta utility functions that modify the RosettaScripts [68] user interface at runtime, adding user-facing syntax that permits configuration of Masala modules loaded at runtime through a RosettaScripts XML script. Section 10.3 explains how we modified the Rosetta `PackRotamersMover` [1], which ordinarily invokes the Rosetta `packer` to perform rotamer optimization and sequence design, to allow it to substitute Masala CFN optimizers loaded at runtime. This section also describes how these changes propagate to modules that use the `PackRotamersMover`, such as `FastRelax` [87, 88], `FastDesign` [18], and Rosetta applications configured at the command-line. Sections 10.4 and 10.5 describe modifications to design-centric guidance functions to permit them to be used with Masala CFN solvers (with greater efficiency than they could be used with the Rosetta `packer`) and to request appropriate Masala data representations for a given solver. In section 10.6, we describe template preferred data representations, which allow users to specify which of several data representations compatible with a given solver should be used. Section 10.7 briefly covers modifications made to the `MinMover` [1], which normally calls the Rosetta `minimizer`, to allow it to call Masala RVL optimizers loaded at runtime (analogous to the `PackRotamersMover` modifications described in section 10.3). These too propagate to `FastRelax`, `FastDesign`, and other larger protocols and applications that use the `minimizer`. Finally, section 10.8 summarizes how new Masala plugin engines and data representations can be added in the future and used in Rosetta without any further modifications to the Rosetta source code.

### 10.1 Modifications to initialization subroutines of external code (Rosetta)

In order for external software to load Masala plugins, it is typically necessary to modify the subroutines that are invoked on initialization, querying user inputs for paths to Masala plugins and calling the `MasalaPluginLibraryManager` to load these libraries and register the Masala plugin modules that they contain. The Masala patch for Rosetta adds the function shown in Listing 16, which is called by Rosetta’s `core::init()` function, which in turn is called on initialization. Similar code could be added to any other software to load Masala plugins from user-provided plugin paths.

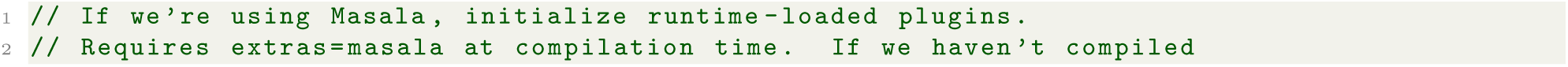
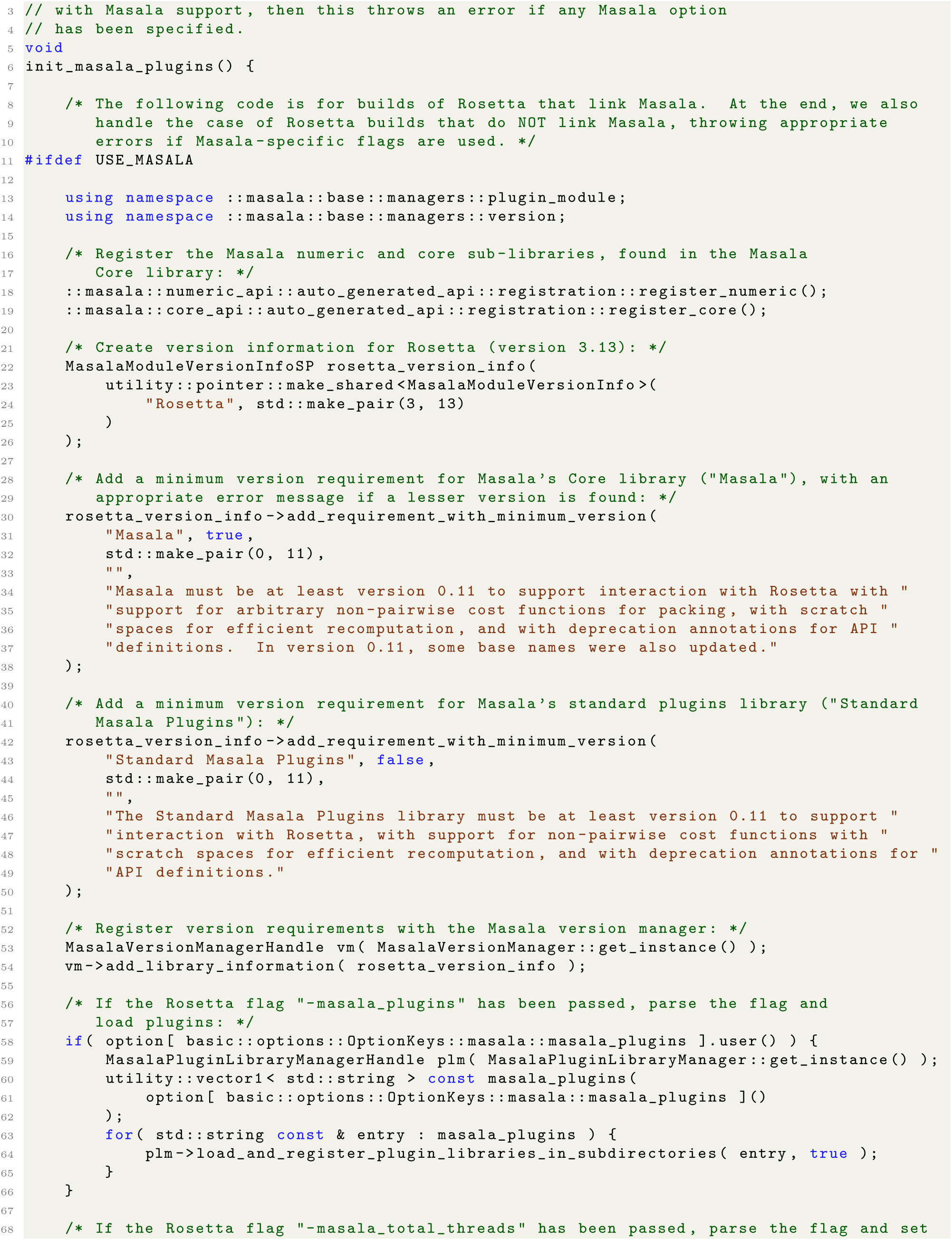
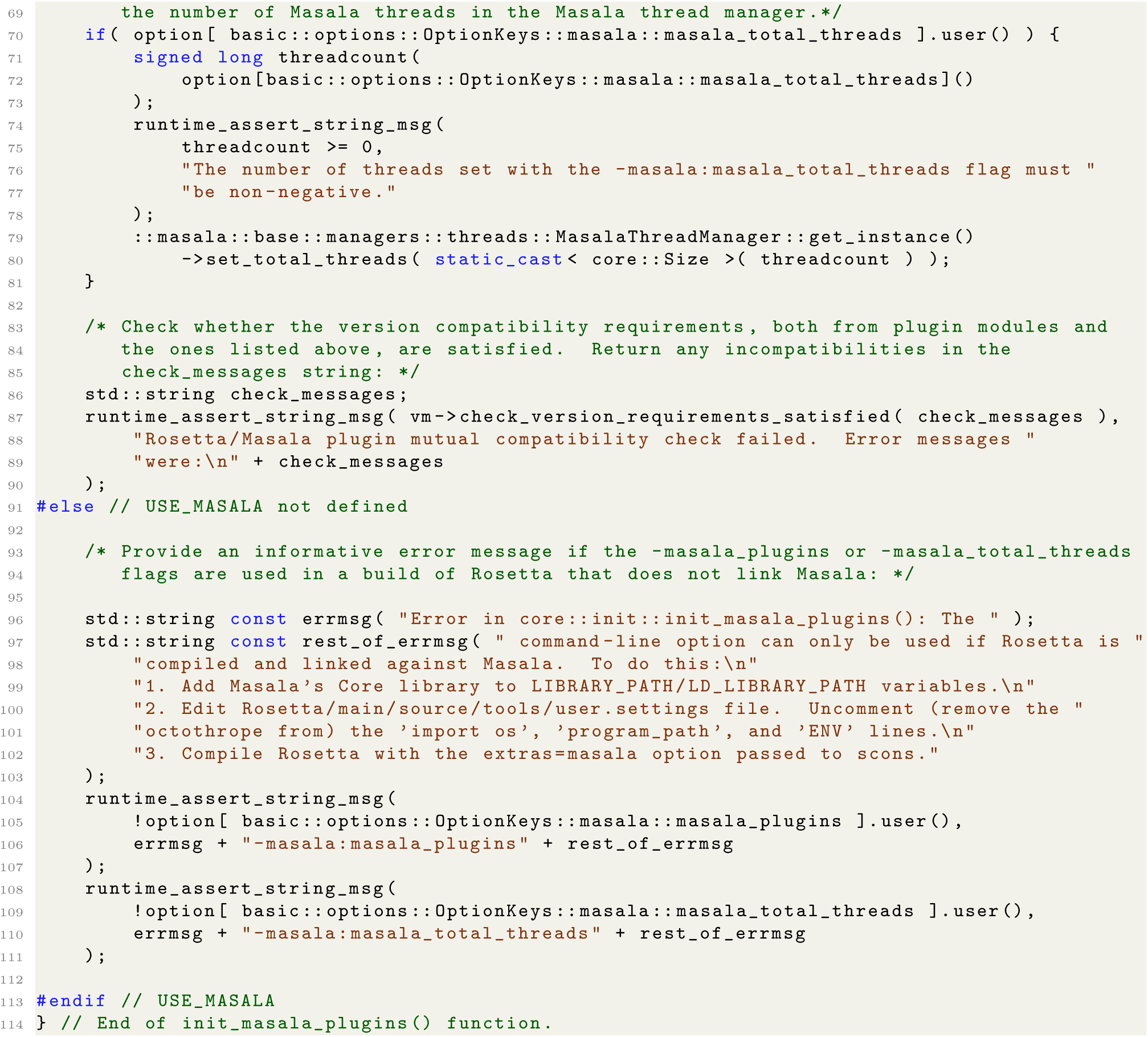

**Listing 16.** Code added to the patched version of Rosetta to permit initialization of Masala plugins from paths passed to Rosetta applications on the command-line with the `-masala_plugins` command-line flag. This function also sets the number of threads launched by the Masala thread manager, based on the `-masala_total_threads` flag passed on the command-line. The `init_masala-plugins`() function is found in `Rosetta/main/source/src/core/init.cc` in the patched version of Rosetta.

### 10.2 Adding Masala user interface elements to the RosettaScripts user interface

The first challenge in making it possible to use Masala modules in other software, such as Rosetta, is providing a user interface for modules unknown to the other software at compilation time. Since Masala’s API definition system provides a full description of all publicly-accessible functions in Masala plugin modules, including setter functions, this information can be used to generate user interfaces automatically. Here, we show how we did this to extend the RosettaScripts scripting language, an XML-style language used to configure Rosetta modules and to chain them together to build Rosetta protocols, giving RosettaScripts users the ability to configure Masala plugin modules that were loaded at runtime and not known to Rosetta at compilation time. The pattern followed here may be emulated in other software as well.

Extending RosettaScripts required two steps. First, it was necessary to extend the RosettaScripts XML schema definition (XSD), the machine-generated and machine-readable specification of the allowed inputs that can be recognized by RosettaScripts, to include the interface elements for Masala plugin modules loaded at runtime. Second, we had to add the ability to parse the new XML elements that were added to the XSD, and to configure the appropriate modules from user inputs provided in those elements. The patch for the Rosetta source code that we have provided adds a new file, `Rosetta/main/source/src/utility/xsd - utility/masala util.cc` (and a corresponding `masala_util.hh` header file), which contains utility functions both for modifying the RosettaScripts XSD and for parsing added elements from RosettaScripts XML.

Listing 17 shows a portion of the function `utility::xsd utility::generate xml schema for masala - plugin object tag(·)`. This function modifies the RosettaScripts XSD, adding to it definitions for a tag or sub-tag with user-configurable attributes with names matching zero- or one-input setter functions in a Masala plugin object. For clarity, only a portion of this function is shown. Note that this function supports nesting, with XML tags for outer Masala objects containing XML tags for contained Masala objects used by those outer Masala objects.

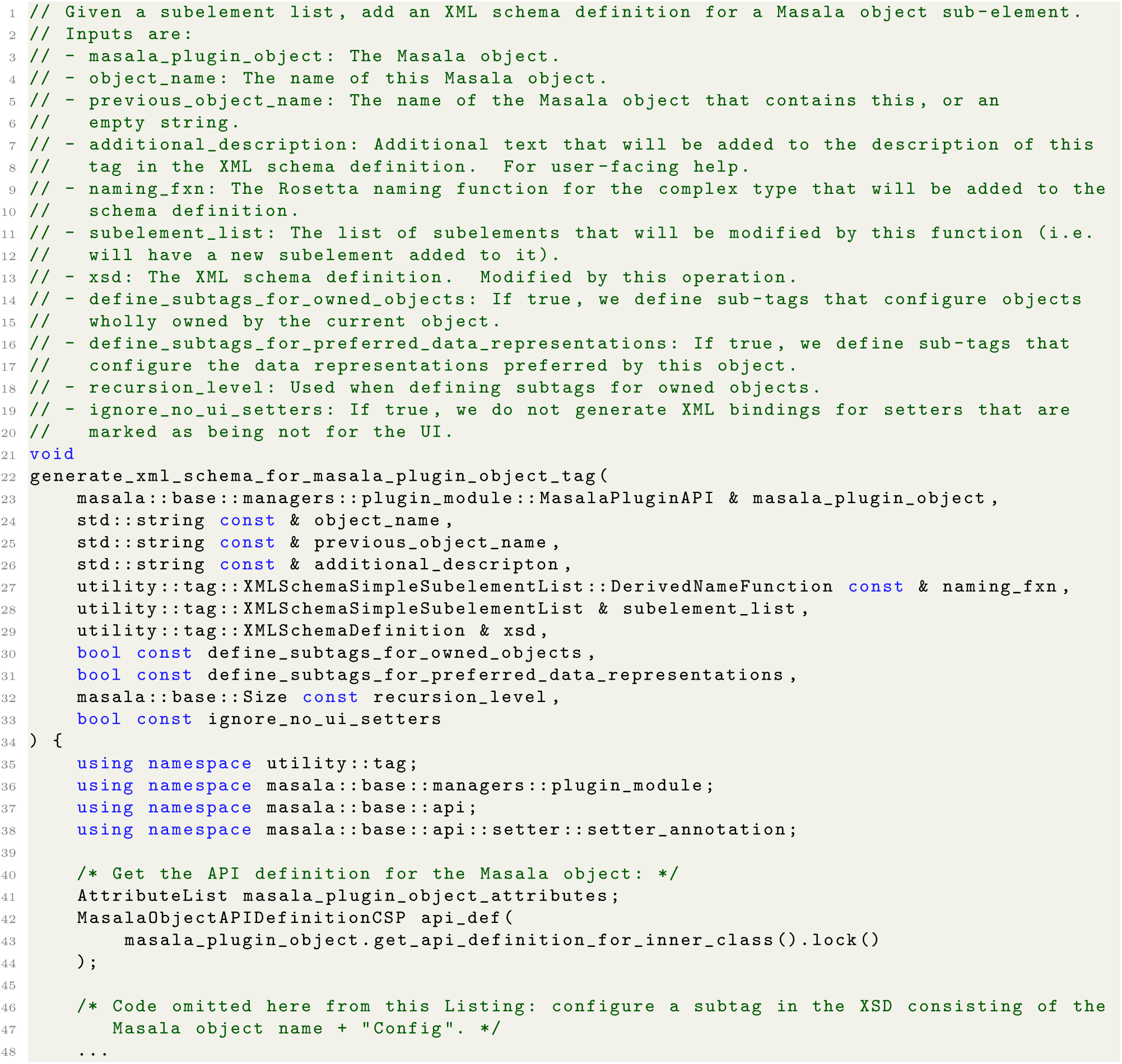
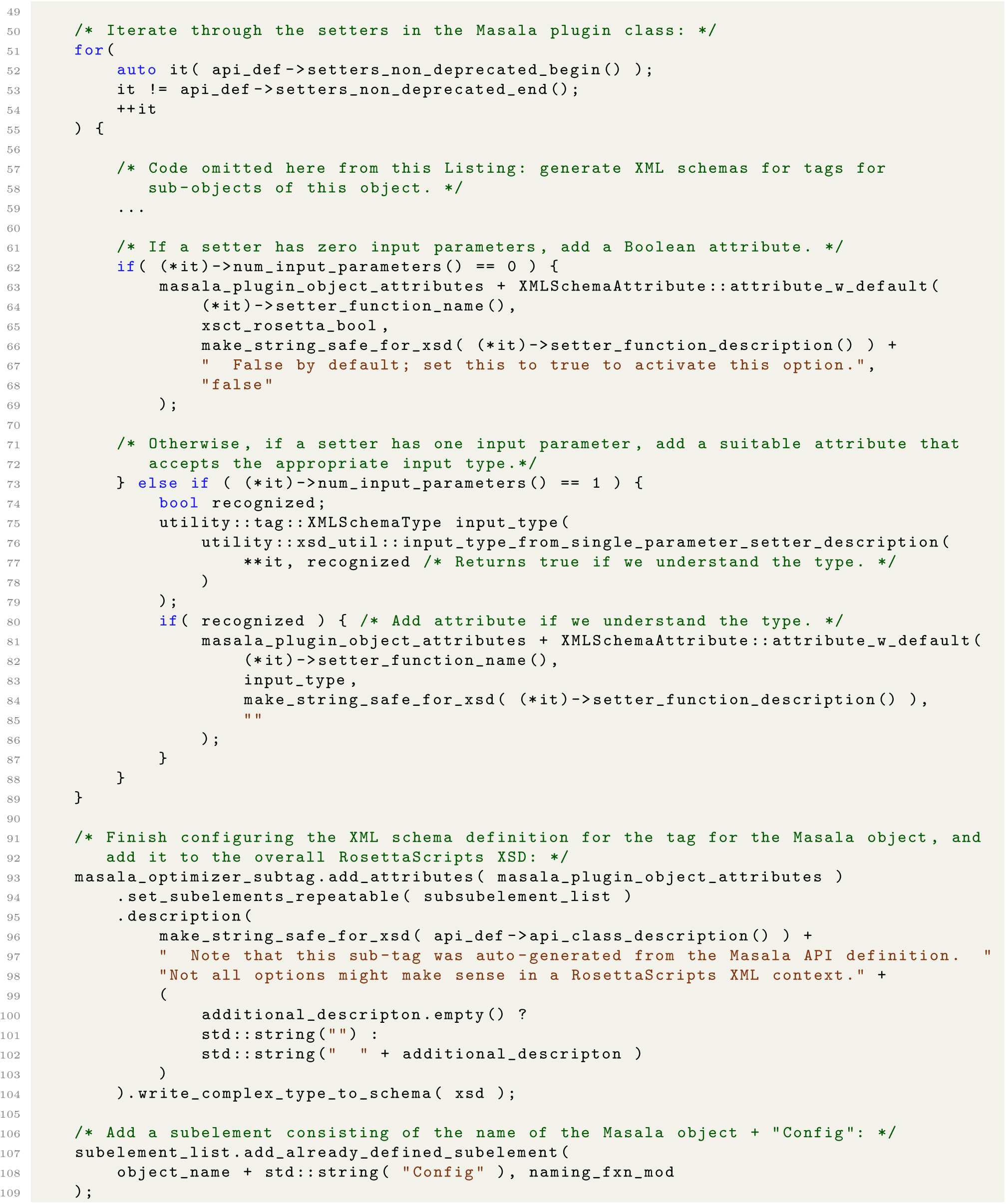
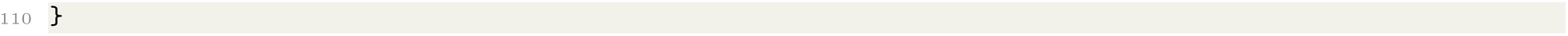

**Listing 17.** The function `utility::xsd_utility::generate xml_schema_for_masala_plugin_object_tag(·)` (abridged), found in `Rosetta/main/source/src/utility/xsd util.cc`. This function modifies the RosettaScripts XSD in order to add a tag with attributes that match zero- and one-input setter functions in a given Masala object’s API definition.

Listing 18 shows the `parse_masala_setters_function(·)` function, which, given a RosettaScripts XML tag and a Masala plugin object, parses subtags configuring that object, and passes the parsed values to corresponding setters in the Masala object through function definitions in the API definition that contain function pointers to the setters. It is the complement of the `generate_xml_schema_for_masala_plugin - object tag(·)` function shown in Listing 17.

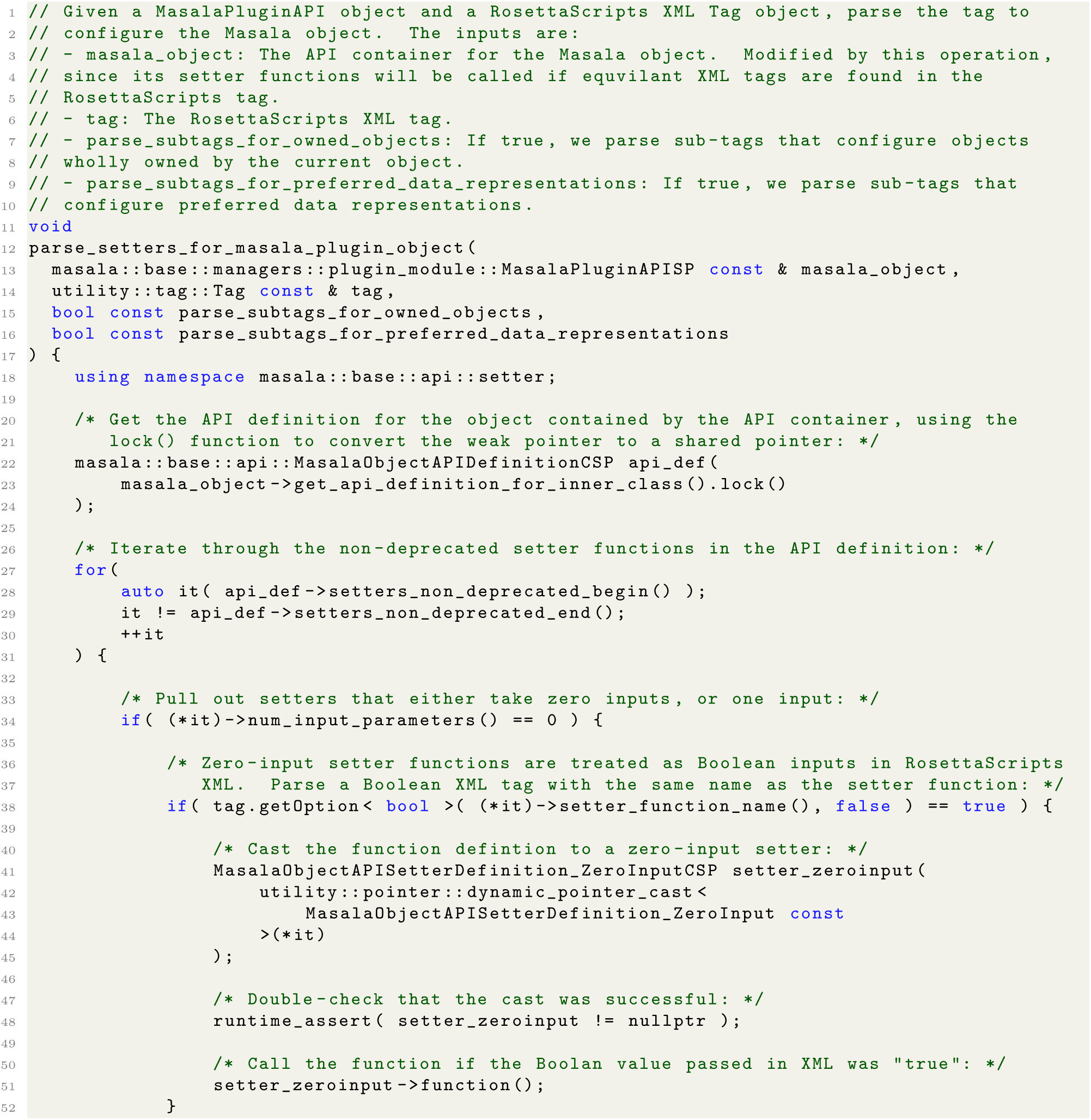
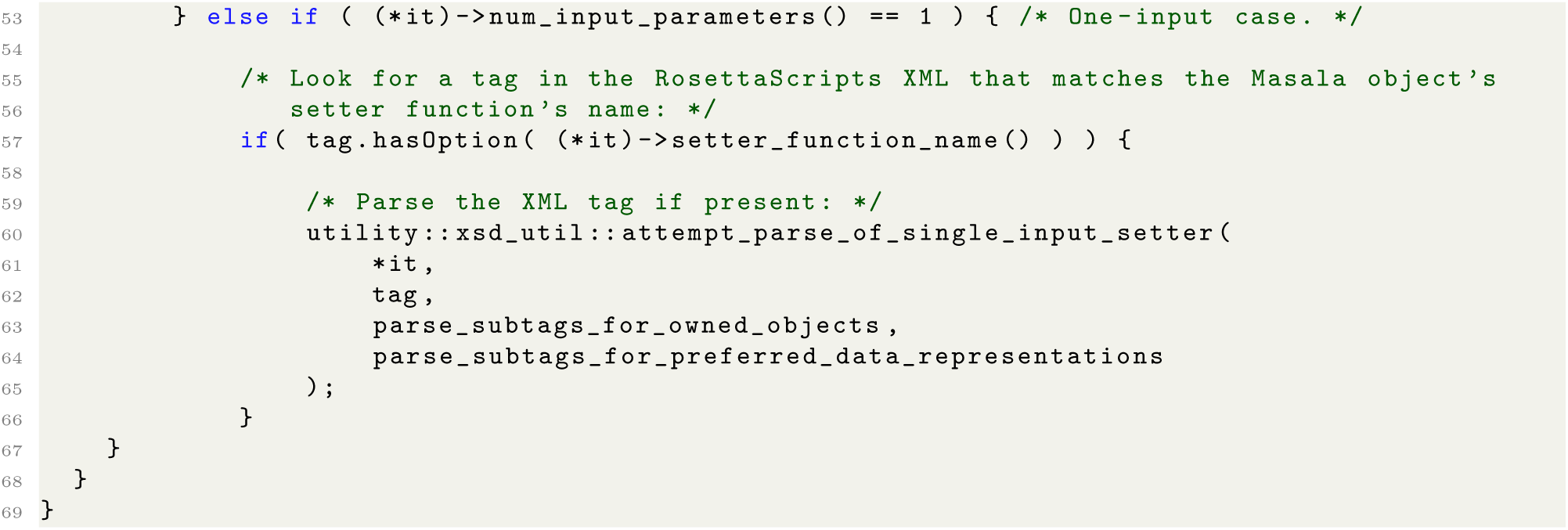

**Listing 18.** Rosetta function `utility::xsd_utility::parse_masala_setters_function(·)`, which parses an XML tag to provide inputs to the setters of a Masala object. This function is defined in the patched Rosetta codebase in file `Rosetta/main/source/src/utility/xsd_util/masala_util.cc`.

These two utility functions shown in Listings 17 and 18 are used by both the Rosetta `PackRotamersMover`, to configure Masala CFN solver engines to replace the Rosetta `packer` (section 10.3), and by the Rosetta `Minmover`, to configure Masala RVL solver engines to replace the Rosetta `minimizer` (section 10.7). They can also be recycled for any other Rosetta modules that may be adapted to to use Masala engines in the future.

### 10.3 Modifications to the Rosetta PackRotamersMover to use Masala CFN solver engines

We used the functions shown in Listings 17 and 18 to modify the `PackRotamersMover::provide_xml_- schema(·)` function, which provides an XML schema definition for the `PackRotamersMover`’s RosettaScripts user interface, and the `PackRotamersMover::parse_my_tag(·)` function, which parses an XML tag to configure the `PackRotamersMover`. These modifications extended the `PackRotamersMover’s` XML interface so that the user could configure a Masala CFN solver engine in the same block of XML code in which a `PackRotamersMover` instance is configured, with particular CFN engines loaded at runtime from Masala plugin libraries. Please refer to `Rosetta/main/source/src/minimization packing/PackRotamersMover.cc` in the patched version of Rosetta for details.

Rosetta `Mover` subclasses all implement an `apply(·)` function, which operates on a Rosetta `Pose` object (Rosetta’s representation of a molecular structure), altering it in some way. The `PackRotamersMover` accepts a set of Rosetta `TaskOperation` objects that define the allowed rotamers in a rotamer optimization problem. When `PackRotamersMover::apply(·)` is called, these are used to generate a set of rotamers; the Rosetta `packer` is then called to perform the actual rotamer optimization. The `PackRotamersMover` then copies the optimized rotamers back into the `Pose` object. We modified the `PackRotamersMover::apply(·)` function as shown in Listing 19 to permit a Masala CFN solver engine to be used in place of the Rosetta `packer`.

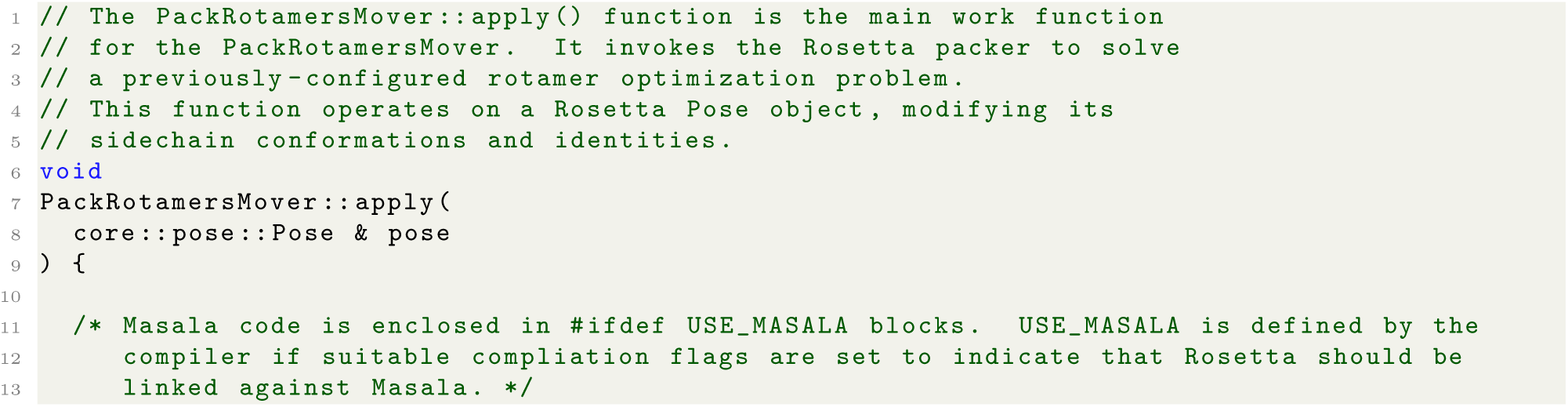
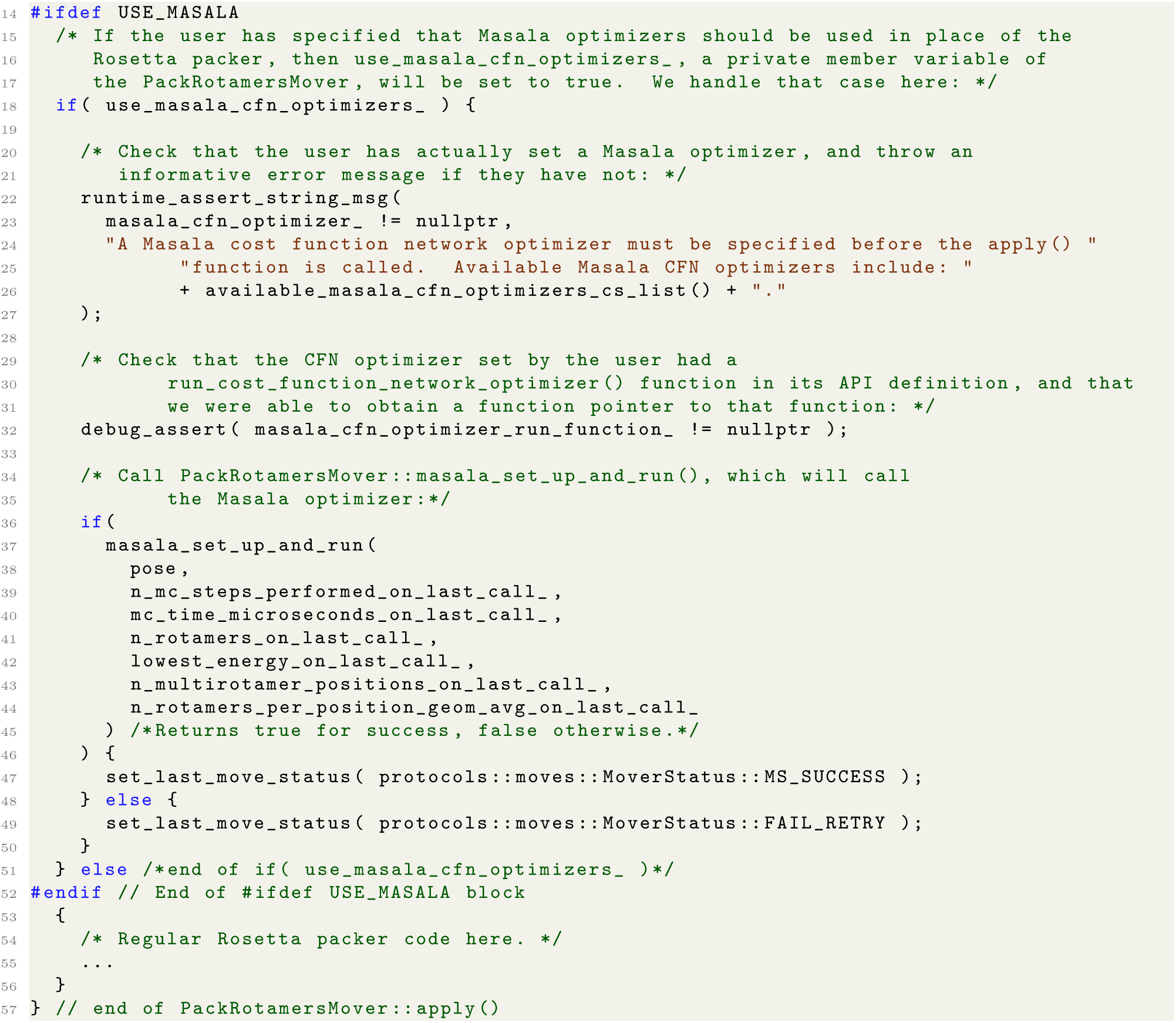

**Listing 19.** Modifications to Rosetta’s `PackRotamersMover::apply(·)` function that ensure that `PackRotamersMover::masala_set_up_and_run(·)` is called if the user has specified that a Masala optimizer be used in place of the Rosetta `packer`.

In the above, a comma-separated list (*i.e.* a string) of the available Masala cost function network optimizers is obtained by calling the `PackRotamersMover::available masala cfn optimizers cs list()` function, which interrogates the `MasalaPluginModuleManager` for any available CFN optimizers, as shown in Listing 20. (We also used this function in the `PackRotamersMover::provide_xml_schema(·)` function, which reports the XML interface for the `PackRotamersMover`; see `Rosetta/main/source/src/minimization - packing/PackRotamersMover.cc` for details.) Note that the output of this function will vary depending on what Masala plugin libraries have been loaded.

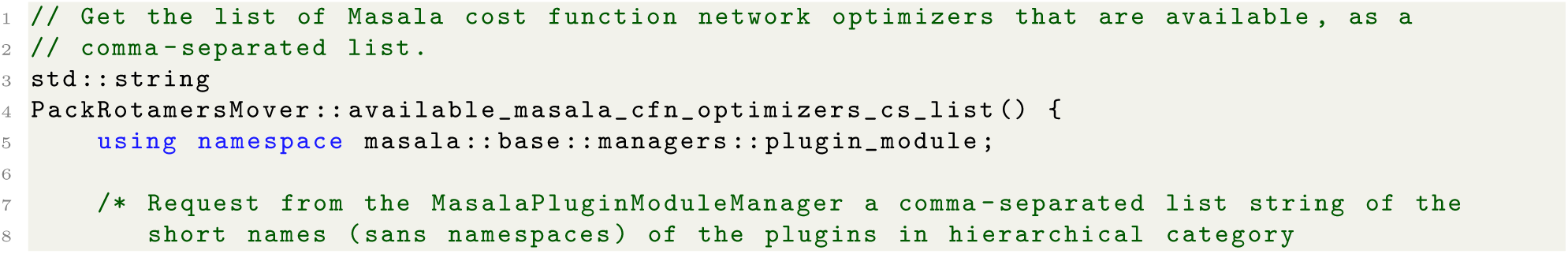
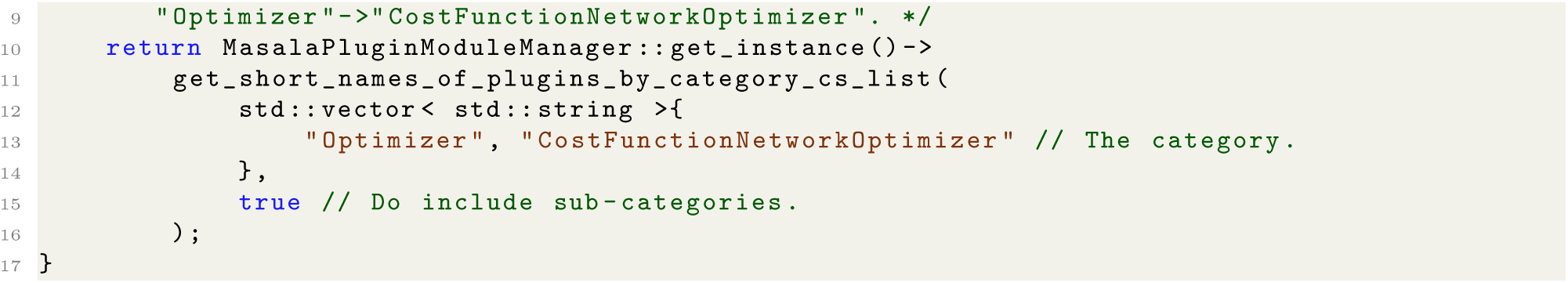

**Listing 20.** The `PackRotamersMover::available_masala_cfn optimizers_cs_list()` function, which provides a string consisting of a comma-separated list of the names of all available Masala CFN optimizers.

Shown in Listing 19, the `PackRotamersMover::masala_set_up_and_run(·)` function, which is called from `PackRotamersMover::apply(·)`, contains much of the code for populating the Masala CFN data representation and for using Masala CFN solver engines. Listing 21 shows a simplified version of this function, highlighting the key steps.

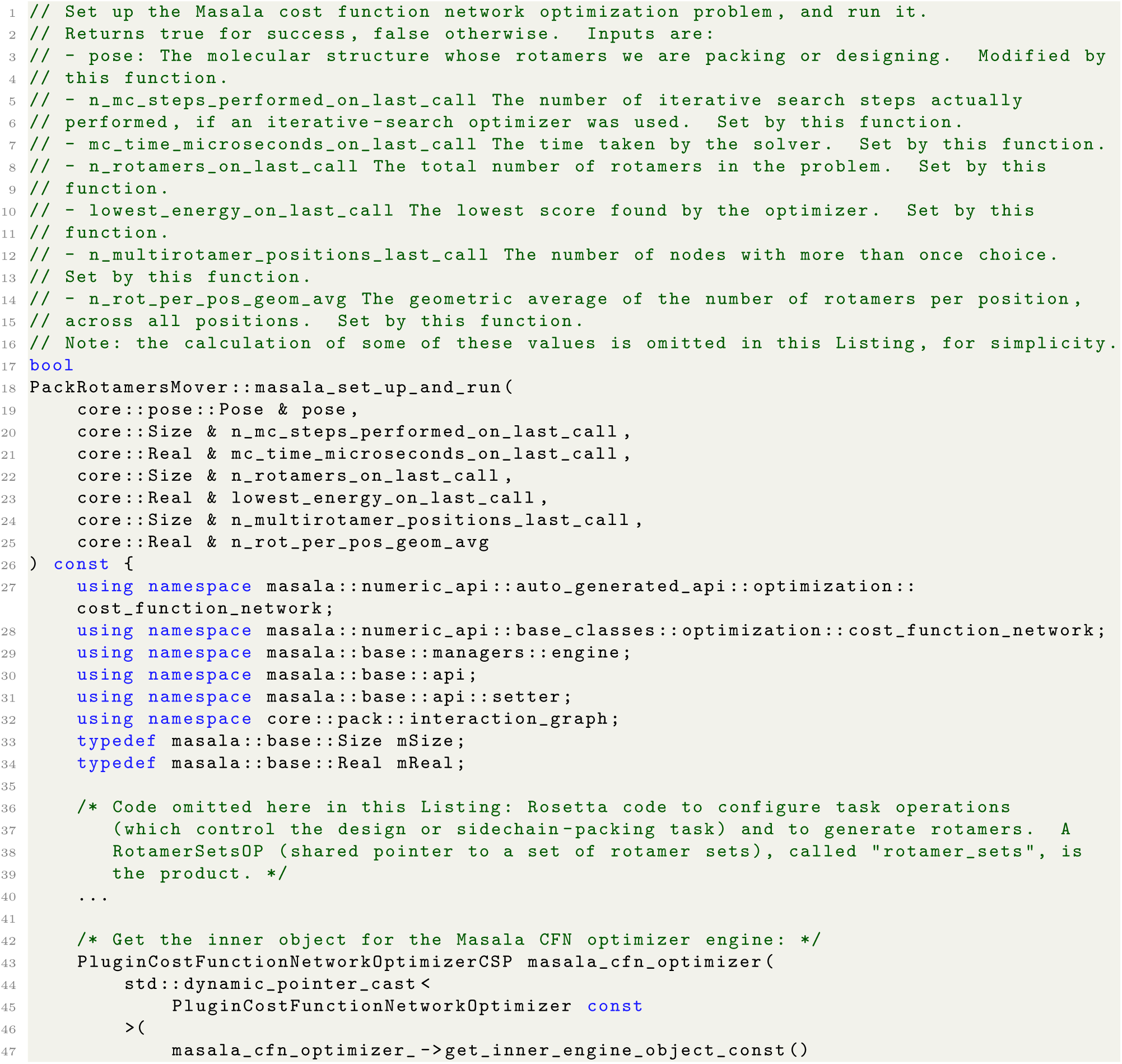
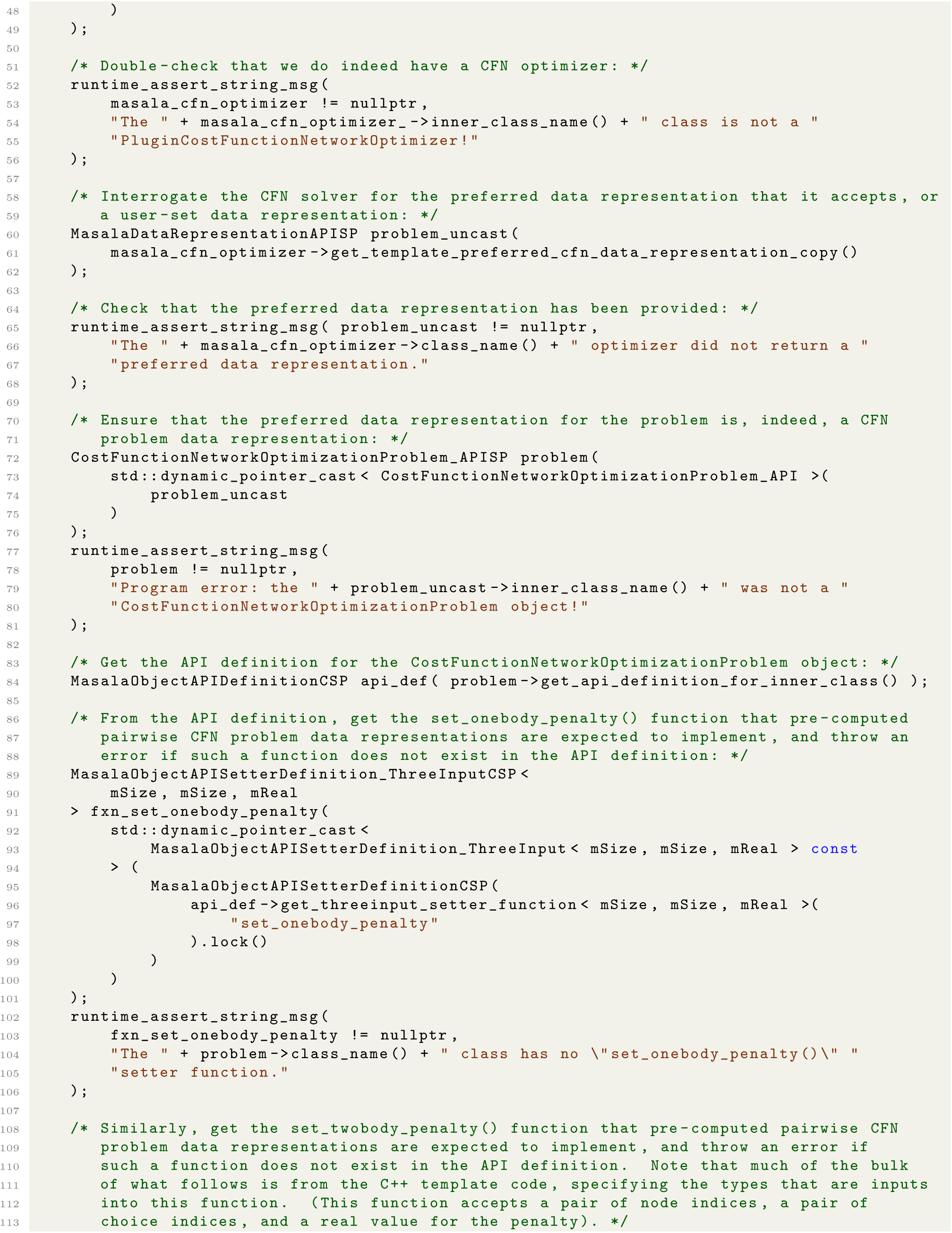
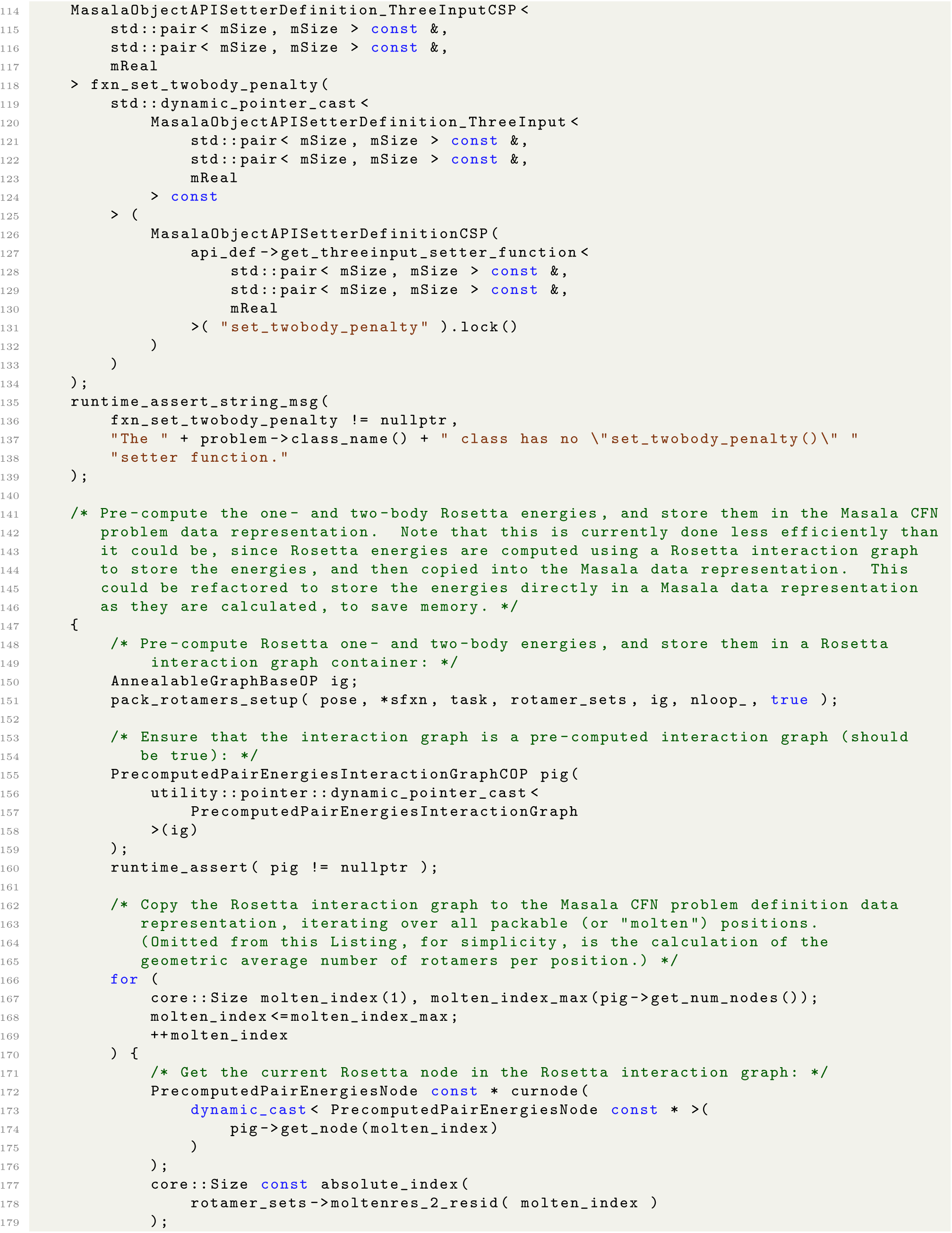
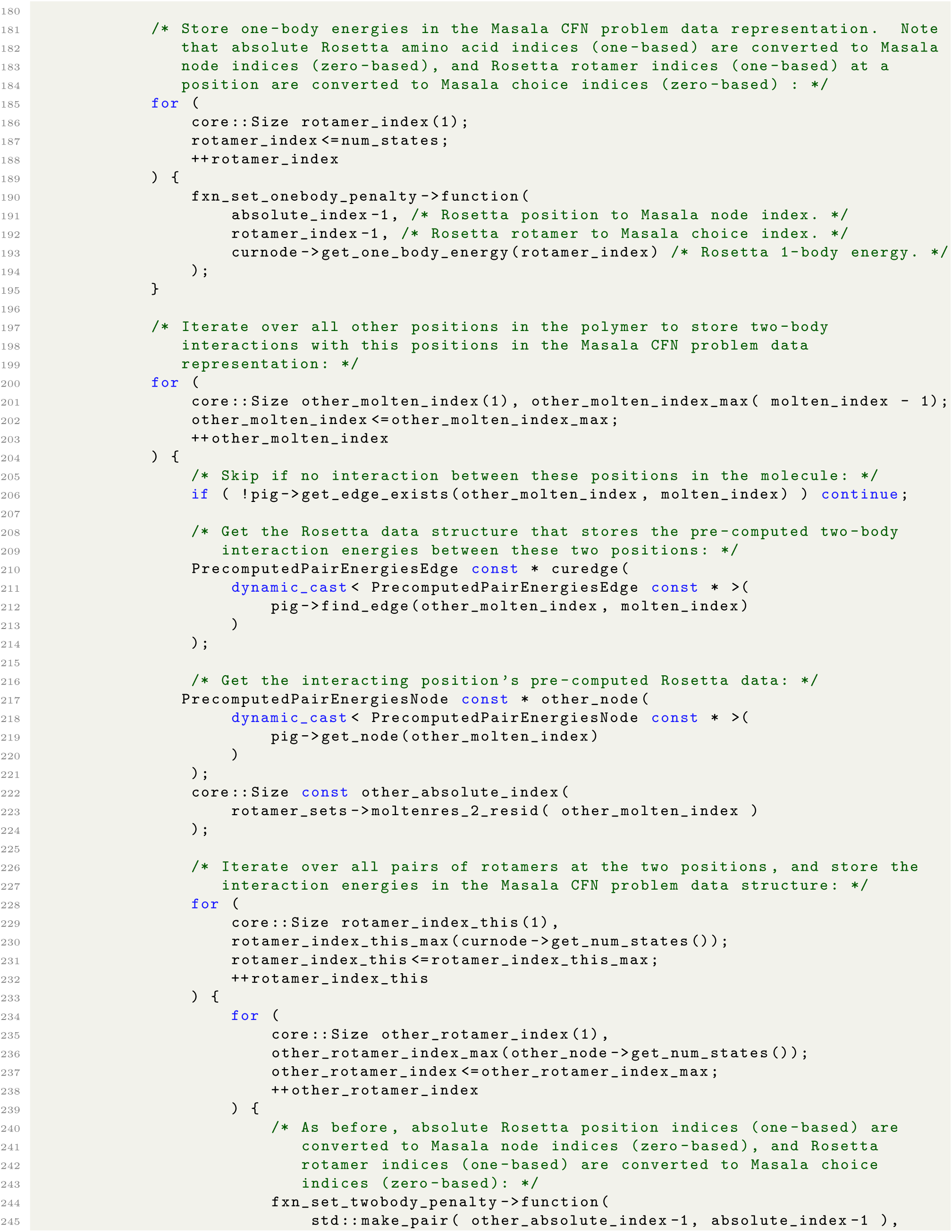
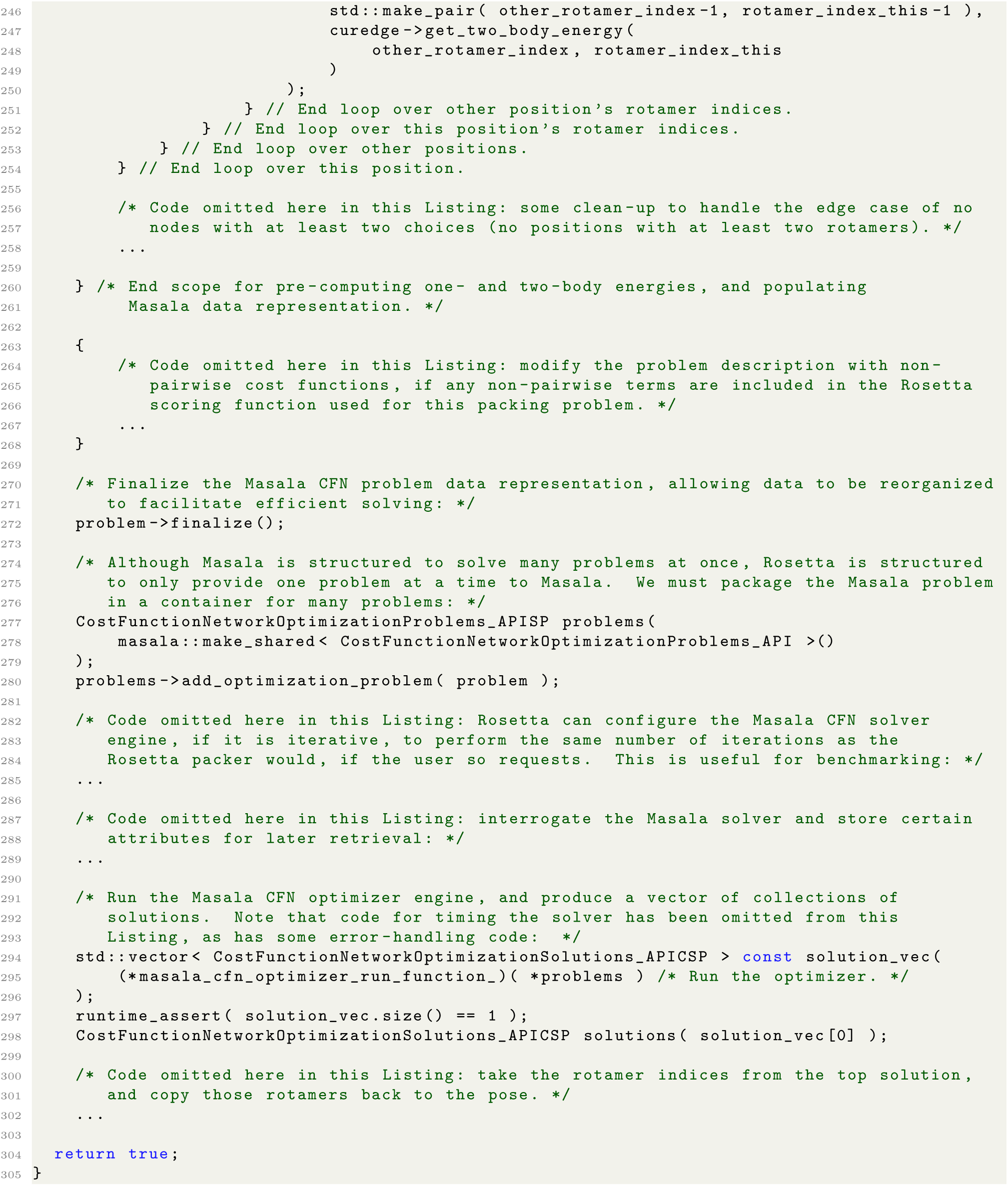

**Listing 21.** Abridged contents of the `PackRotamersMover::masala_set_up_and_run(·)` function, showing the main parts that set up Masala CFN data representations and run the Masala CFN solver engine. See `Rosetta/main/source/src/protocols/minimization_packing/PackRotamersMover.cc` in the version of Rosetta patched to use Masala for full detail.

Note that the problem setup includes some steps that involve interrogating the Masala API definition for the data representation, and dynamically casting Masala function definitions prior to using them, as well as some runtime checks. There is of course some computational cost associated with this. However, these are steps performed once at the start of each problem, likely costing a few microseconds of computing time; the optimizer will then run for tens of seconds. Additionally, the function definitions are extracted and dynamically cast once, then used many times to set one- and two-body penalties in the problem, avoiding unnecessarily repeated casts.

The modifications to `PackRotamersMover::provide_xml_schema(·)` and `PackRotamersMover::parse_-_my_tag(·)` allow the `PackRotamersMover` to be configured in a RosettaScripts XML script to use Masala CFN solvers. Since XML tags can be constructed in a PyRosetta script and the `PackRotamersMover::parse_my_-_tag(·)` called from Python code, this also provides a means of using Masala CFN solvers from PyRosetta, though future support for a more direct interface, with Python bindings generated directly from the Masala API definitions, is planned. But what about other Rosetta protocols that use the `PackRotamersMover`? How can we ensure that Masala CFN solvers can be configured in those contexts?

There are two relevant contexts. The first is other `Mover` subclasses that execute more complicated protocols, calling the `PackRotamersMover` internally. The `FastRelax` [87, 88] and `FastDesign` [18] `Mover` subclasses are examples. For these, we propagated the changes that we had made to the `PackRotamersMover` XML interface to these `Mover` subclasses, ensuring that they could be configured with Masala CFN solvers from RosettaScripts scripts using the same syntax. The second is Rosetta applications and protocols that are controlled at the command-line. For these, we added support for setting a default `PackRotamersMover`, used wherever the `packer` is invoked unless overridden by other commands, with the option to provide a RosettaScripts-format XML tag configuring a `PackRotamersMover` using the command-line flag `-default_ - masala_packer_configuration <PACKROTAMERSMOVER XML configuration file>`. This file provides the opportunity to configure a Masala CFN solver using the same RosettaScripts syntax again. The user interface is discussed further in section 3.4.

The net effect of these changes is that plugin Masala CFN solvers may be used in any existing Rosetta protocol in place of the Rosetta `packer`, an optimizer that accounts for a large fraction of Rosetta’s runtime in most design, validation, or other modelling protocols, without further alterations to Rosetta’s source code or any recompilation of Rosetta. This permits rapid cycles of developing new CFN solvers and testing them in real-world contexts in order to explore new methods of solving difficult rotamer optimization problems efficiently.

### 10.4 Adapting the **hbnet** design-centric guidance term to use Masala data representations

Design-centric guidance terms (reviewed in [42] and [43]) are additional terms added to the scoring function used during design, which help to impose additional user-defined constraints on the rotamer selection process to guide designs towards desired features or away from undesired features. Unlike the default energy terms, design-centric guidance terms need not be pairwise-decomposable. We adapted five of Rosetta’s design-centric guidance terms to be compatible with Masala’s solvers, in most cases achieving significant speedups through better data representations. In this section, we describe the modifications to the `hbnet` term in detail. The changes to `aa_composition`, `netcharge`, `buried_unsatisfied_penalty`, and `voids_penalty` are summarized more briefly in section 10.5.

#### 10.4.1 Rosetta’s original **hbnet** design-centric guidance term

Natural proteins fold and self-assemble spontaneously in water, driven primarily by the hydrophobic effect [89, 90] but further stabilized by hydrogen bond networks. Unlike hydrophobic interactions, which tend to be fairly nonspecific and create broad wells in the conformational free energy landscape [91], hydrogen bonds are short-range, highly directional interactions that only occur between particular donor and acceptor atoms. As a consequence, they tend to stabilize very particular conformations, creating deep but narrow wells in the energy landscape [92, 93]. When patterned hydrogen bond networks form, this tends to lend a great deal of stability and rigidity to a folded structure or complex, which is often desirable. This is particularly important for the rational design of cyclic peptides able to bind to polar surfaces on proteins, where hydrogen bond networks can be the key to both affinity and specificity.

The rational design of hydrogen bond networks has long been a challenge for designers of proteins and other folding heteropolymers using combinatorial optimization tools like the Rosetta `packer`, since hydrogen bond networks are *non-pairwise decomposable* features: recognizing one requires consideration of more than two amino acid residues at at time. In 2016, Boyken *et al.* introduced the Rosetta `HBNet` mover [94], a module that used a specialized optimization algorithm to introduce sidechain hydrogen bond networks onto a protein backbone; in a subsequent step, the Rosetta `packer` could be used to optimize the rest of the sequence. Maguire *et al.* later accelerated the hydrogen bond placement by replacing the deterministic algorithm with stochastic Monte Carlo methods [95], but these still relied on subsequent optimization of other sidechains using the `packer`. While useful, these approaches suffered from the two steps of optimizing for two different desired properties: the first step could produce hydrogen-bonded sidechain patterns that hindered optimization for favourable hydrophobic packing in the second step. More modern machine learning methods for protein design have not yet proven adept at optimizing this particular feature. To address this, in 2017 we added the `hbnet` design-centric guidance term to Rosetta.

The `hbnet` term is designed to provide a bonus, `H` (*s⃗*) for the formation of a hydrogen bond network given rotamer selection *s⃗*, where the magnitude of the bonus is dependent on the size of the network, and where the bonus can be rapidly updated at each step during the simulated annealing search of rotamer selections performed by the Rosetta `packer`. The term is multiplied by a user-set weight *λ*_hbnet_, and then added to the weighted sum of the energetic terms *E*_1_ . . . *E_n_* to yield the overall score 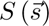, as shown in Eq. (9).

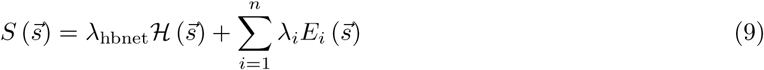

The `hbnet` term’s C*++* code was implemented in the Rosetta codebase in `src/core/pack/guidance_- scoreterms/hbnet_energy/HBNetEnergy.cc`, and in the corresponding `HBNetEnergy.hh` header file and `HBNetEnergy.fwd.hh` forward declarations file. Within the Rosetta source code, the `HBNetEnergy` class, defined in namespace `core::pack::guidance_scoreterms::hbnet_energy`, multiply inherits from classes `WholeStructureEnergy` (in namespace `core::scoring::methods`) and `ResidueArrayAnnealableEnergy` (in namespace `core::scoring::annealing`). The former defines it to be a non-pairwise decomposable scoring term that would ordinarily be incompatible with the Rosetta `packer`, but the latter ensures that it is of a special sub-category of non-pairwise scoring terms which, although not pairwise-decomposable, can be otherwise made fast to compute and fast to update during the `packer` simulated annealing search.

During `packer` setup, the ordinary pairwise-decomposable energy function is used to pre-compute one-body energies for all rotamers (an 𝒪(*ND*) calculation) and two-body energies for all interacting rotamer pairs (a typically 𝒪(*ND*^2^) calculation) (**Fig. 8A**). During this setup phase, `ResidueArrayAnnealableEnergy`-derived terms that implement a `set_up_residuearrayannealableenergy_for_packing(·)` function are permitted to cache data needed for the subsequent simulated annealing search. The `hbnet` term begins by pre-computing and caching backbone-backbone hydrogen bonds, which ordinarily cannot be altered by the `packer`. These are stored in a heap-allocated graph structure. Two hydrogen bonding graphs of *N* nodes each (where *N* is the number of positions in the structure or `Pose`), one for the current hydrogen bond network under consideration and another for the hydrogen bond network for the last accepted move, are also heap-allocated.

**Fig 8.**
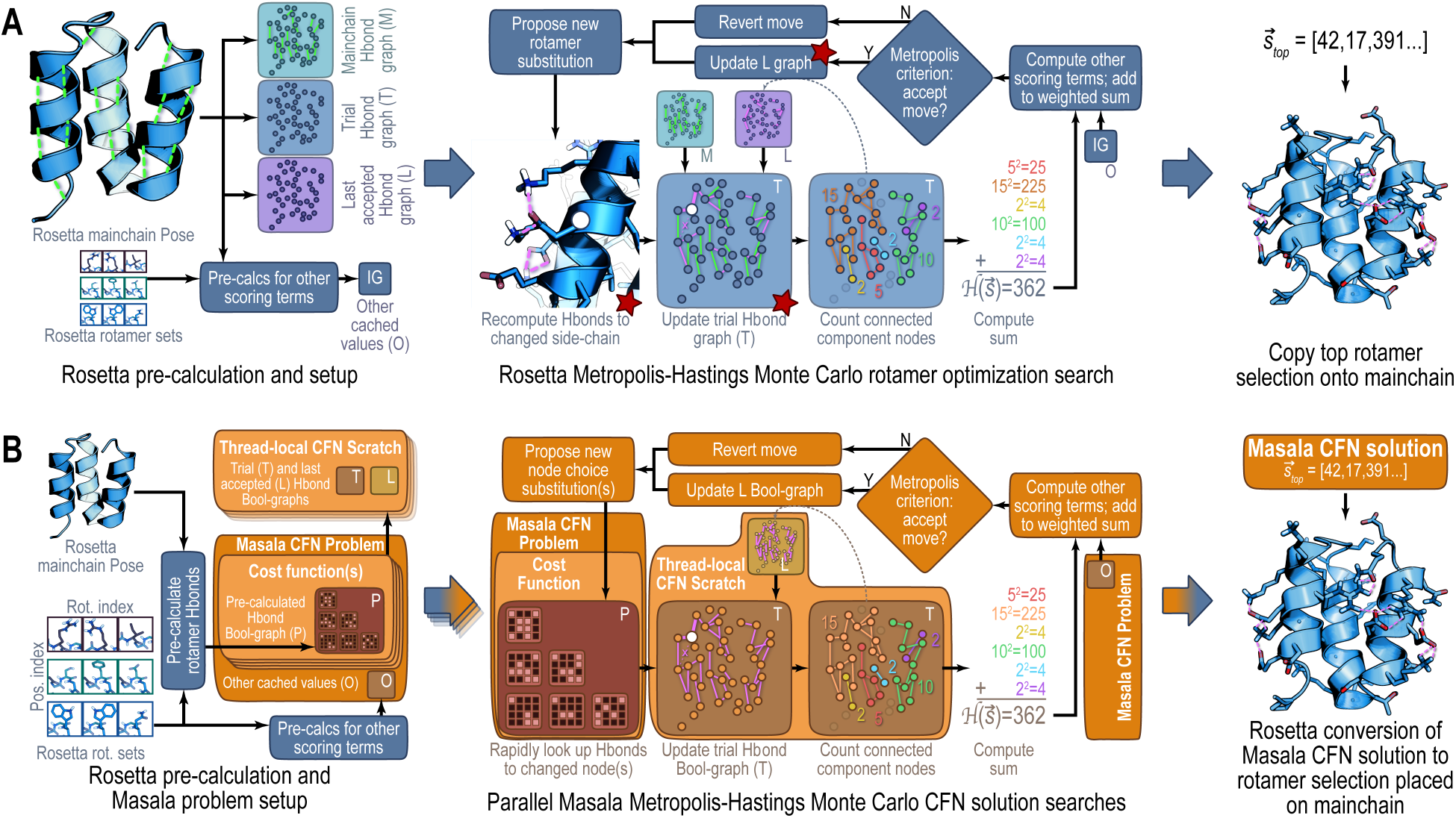
Schematic of original Rosetta and new Masala-enhanced `hbnet` design-centric guidance terms. (**A**) Original Rosetta `hbnet` guidance term. The pre-computation phase was used only to allocate graph structures for use during the search phase, and to identify mainchain-mainchain hydrogen bonds (green) that would not change during rotamer optimization. The search phase required rapid geometry-based identification of hydrogen bonds involving changed sidechain rotamers (white circle) in order to update graphs used for the connected component analysis to score hydrogen bond networks. Red stars mark steps that added to the computational cost, either due to the need to consider explicit molecular geometry in the midst of the Monte Carlo search, or the need to perform heap allocations when updating graph edges or copying graphs. (**B**) Refactored `hbnet` guidance term implemented in Masala and linked by Rosetta. Masala steps are shown in orange and brown, and Rosetta steps in blue. By replacing the search-phase hydrogen bond identification with a pre-computed Boolean graph, and by using a similar data structure for the cached last accepted graph, the search is greatly accelerated. Masala also permits parallelization of the search.

The `packer` then proceeds to its simulated annealing search, randomly substituting one rotamer at a time in the candidate solution vector 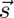, updating the solution score, and accepting or rejecting the move by the Metropolis criterion. At every move, the `hbnet` term updates its computed value of ℋ 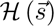 by constructing the graph of hydrogen-bonded positions, computing sidechain-sidechain and sidechain-backbone inter-rotamer hydrogen bonds for all selected rotamers on the fly, and looking up pre-computed backbone-backbone rotamers. The pre-allocated graphs are used as scratch space; however, heap allocations may occur at this step as edges are added or pruned. At the start of the search, or if jumps to new regions of rotamer selection space occur, the graph is computed from scratch; when single rotamer substitutions are performed, the graph is populated from the last accepted and only those edges involving the altered position are recomputed. When a rotamer substitution is accepted, the last accepted graph is updated to match the current graph.

Once the hydrogen bond graph is populated or updated for the current rotamer selection 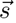, a depth-first search is used to compute a heap-allocated vector of the sizes of all “islands” (all connected components) in the graph. Islands of size 1 are discarded. Given r islands with at least two nodes, of node counts c_1_ . . . c*_r_*, the penalty function ℋ 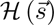 is computed by Eq. (10).

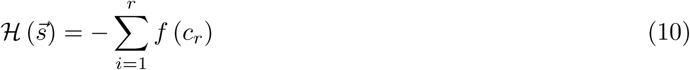

In Eq. (10), *f* (*·*) is selected by the user from a choice of a linear function, a quadratic, a square root function, or a logarithmic function. The quadratic function gives a ramping bonus heavily promoting very large hydrogen bond networks; the square root and logarithmic functions give diminishing returns permitting other terms to dominate once the hydrogen bond networks grow sufficiently large.

In summary, Rosetta’s `hbnet` design-centric guidance term augmented the energy function used for design with the Rosetta `packer`, turning the design optimization problem into a weighted, multi-objective optimization problem in which one objective was minimizing energy and the other was maximizing hydrogen bond networks. This required evaluating a non-pairwise decomposable term during a `packer` simulated annealing trajectory; however, much of the term’s computational cost was not from its non-pairwise nature, but from the need to compute hydrogen bonds (which individually are a pairwise-decomposable feature) on the fly at each simulated annealing step. Despite the significant cost, the `hbnet` design-centric guidance term has proven useful for real-world peptide design challenges, including the design of peptide macrocycle inhibitors of the New Delhi metallo-*β*-lactamase [24].

#### 9.4.2 Refactored **hbnet** with Rosetta linking Masala

As described in Section 2.2.2, the Masala software suite is built around a paradigm of *engines* and *data representations*. When Rosetta is linked against Masala, Rosetta’s `PackRotamersMover`, a module which ordinarily configures a rotamer optimization problem and calls the Rosetta `packer`, can instead request a Masala CFN optimizer engine and corresponding data representation, as described in section 10.3. To refactor the Rosetta `hbnet` design-centric guidance term to work with Masala CFN solvers, it was only necessary to modify the `hbnet` source code in the Rosetta codebase to request and configure a Masala data representation, and to implement a suitable data representation compatible with some subset of Masala’s CFN solver engines.

Rosetta’s `PackRotamersMover` pre-computes one- and two-body terms of the objective function used for optimization. During this phase, the `PackRotamersMover` also iterates over those terms in the scoring function which inherit from `ResidueArrayAnnealableEnergy`, and which are not represented in the pairwise-decomposed pre-computed scores, providing these non-pairwise terms with the opportunity to perform pre-computations of their own by calling each term’s `set_up_residuearrayannealableenergy_for_packing(·)` function override. The Rosetta `hbnet` guidance term, for instance, uses this phase to pre-compute and cache mainchain-mainchain hydrogen bonds, which remain constant across all rotamer selections, so that these need not be calculated on the fly. Masala-compatible `ResidueArrayAnnealableEnergy`-derived classes provide a `generate_masala_cost_functions(·)` function, which produces a vector of one or more Masala `CostFunction` data representation objects which the `PackRotamersMover` encapsulates in the problem description. These are specialized data representations used to compute scoring functions that are non-pairwise decomposable but which can be computed or updated rapidly during a simulated annealing search. The `PackRotamersMover` then passes the data representation describing the problem to the selected Masala CFN solver, runs it, and obtains solution vectors which it converts back to rotamer selections.

The `HBNetEnergy::generate_masala_cost_functions(·)` function returns one of several subclasses of the `CostFunction` class intended for rapidly computing functions of connected component (or “graph island”) node counts. Different subclasses (`LinearGraphIslandCountCostFunction`, `SquareOfGraphIs-landCountCostFunction`, `SquareRootOfGraphIslandCountCostFunction`, and `LogOfGraphIslandCount-CostFunction`) are implemented in namespace `standard_masala_plugins::optimizers::cost function_ - network::cost function::graph island based` of the Standard Masala Plugins library. These provide graph island-based cost functions that calculate the sum of the connected component node counts or a function thereof. The subclass chosen depends both on the user’s choice of Masala optimizer and the user’s choice of *f* (*·*), the function applied to the connected component counts.

The class inheritance for the versions of these cost functions implemented for use with the CPU-based CFN solver engines in the Standard Masala Plugins library, as well as for cost functions discussed in section 10.5, is shown in **Fig. 9**. These versions all use an efficient Boolean matrix representation to cache pre-computed data indicating whether a pair of selected choices at a pair of nodes would be connected by an edge (in this case, representing a hydrogen bond between a pair of rotamers) or not. The particular data structure used is deliberately kept opaque to external code, however, allowing alternative data representations to be swapped in seamlessly. This allows the hydrogen bond identification step, not only for mainchain-mainchain hydrogen bonds but for all hydrogen bonds, to be moved from the search phase to the pre-computation phase, enormously accelerating the Masala CFN optimizer engine’s search for an optimal choice vector 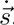. (As discussed in section 4.1, we measured speedups of roughly threefold.) Equivalents for other hardware or other CFN solvers (for instance, GPU-based CFN solvers) will be implemented in other plugin Masala libraries in the future, and will be usable with Rosetta without modification to the Rosetta source code.

**Fig 9.**
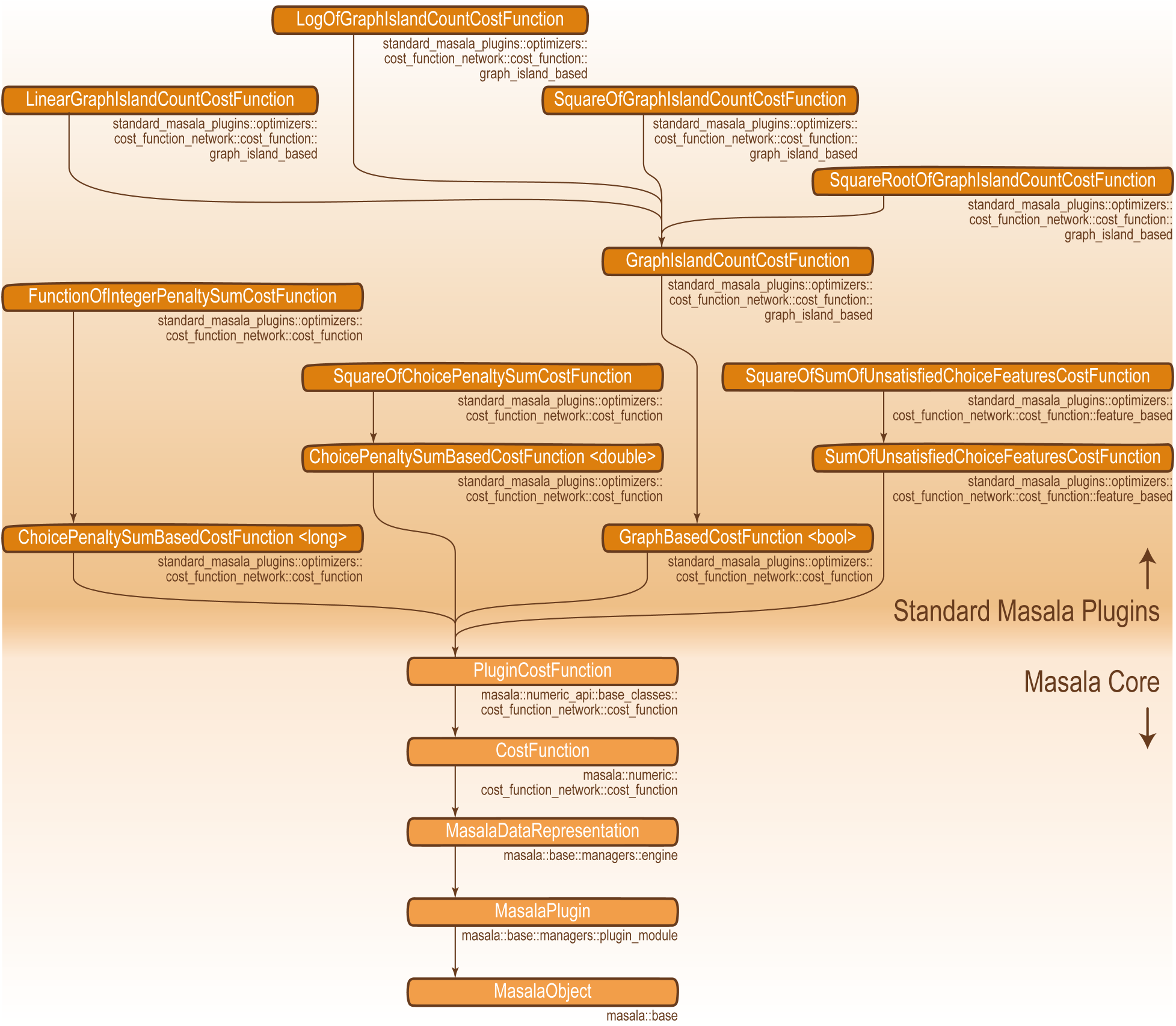
Class inheritance of Standard Masala Plugins cost functions implemented for the `hbnet` term (section 10.4) and other design-centric guidance terms (sec 10.5), for use with CPU-based Masala cost function network optimizer engines. Equivalent cost functions compatible with other solver engines (for instance, GPU solvers), will be implemented in other Masala plugin libraries, and will also inherit from `PluginCostFunction`. prior knowledge or known experimental constraints to be applied during the design process. For instance, it may be desirable to have at least one D- or L-tryptophan or tyrosine residue, or other amino acid type with absorbance around 280 nm, in a cyclic peptide, to facilitate absorbance-based concentration determination after synthesis. Or one may wish to specify that a peptide’s net charge be less than some limit, say +2, to avoid extreme levels of positive charge that can cause a peptide not merely to pass through the cell membrane, but to disrupt it [96]. These constraints are added to the total score weighted by Lagrange multipliers λ_aacomp_ and λ_netcharge_, transforming the CFN optimization problem into a multi-objective optimization problem, as shown in Eq. (11).

Listing 22 shows code added within the Rosetta `HBNetEnergy` class that requests a suitable data representation from the `MasalaDataRepresentationManager`. Given the graph island-based cost function data representation, during the pre-computation phase, the `HBNetEnergy` class iterates through all pairs of interacting residue positions in the current `Pose`, and through all pairs of rotamers for each pair of positions. It determines whether each rotamer pair forms at least one hydrogen bond, and marks those pairs that do in the graph island-based cost function. This cost function is then returned and ultimately used in any search performed by the Masala CFN optimizer engine, including the simulated annealing search of the `MonteCarloCostFunctionNetworkOptimizer` or the greedy trajectory of the `GreedyCostFunctionNetworkOptimizer` (see section 9.1.1 of Appendix C).

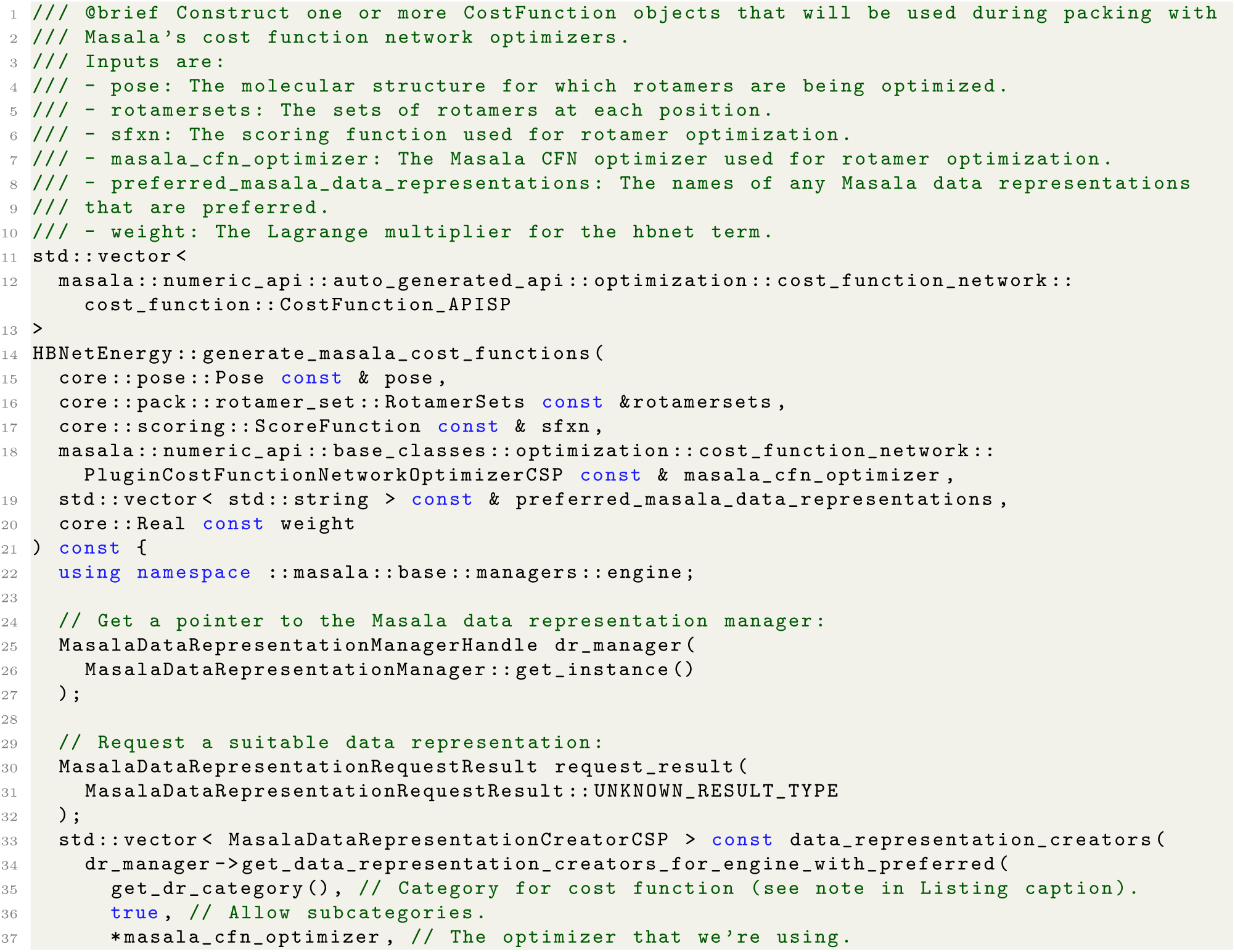
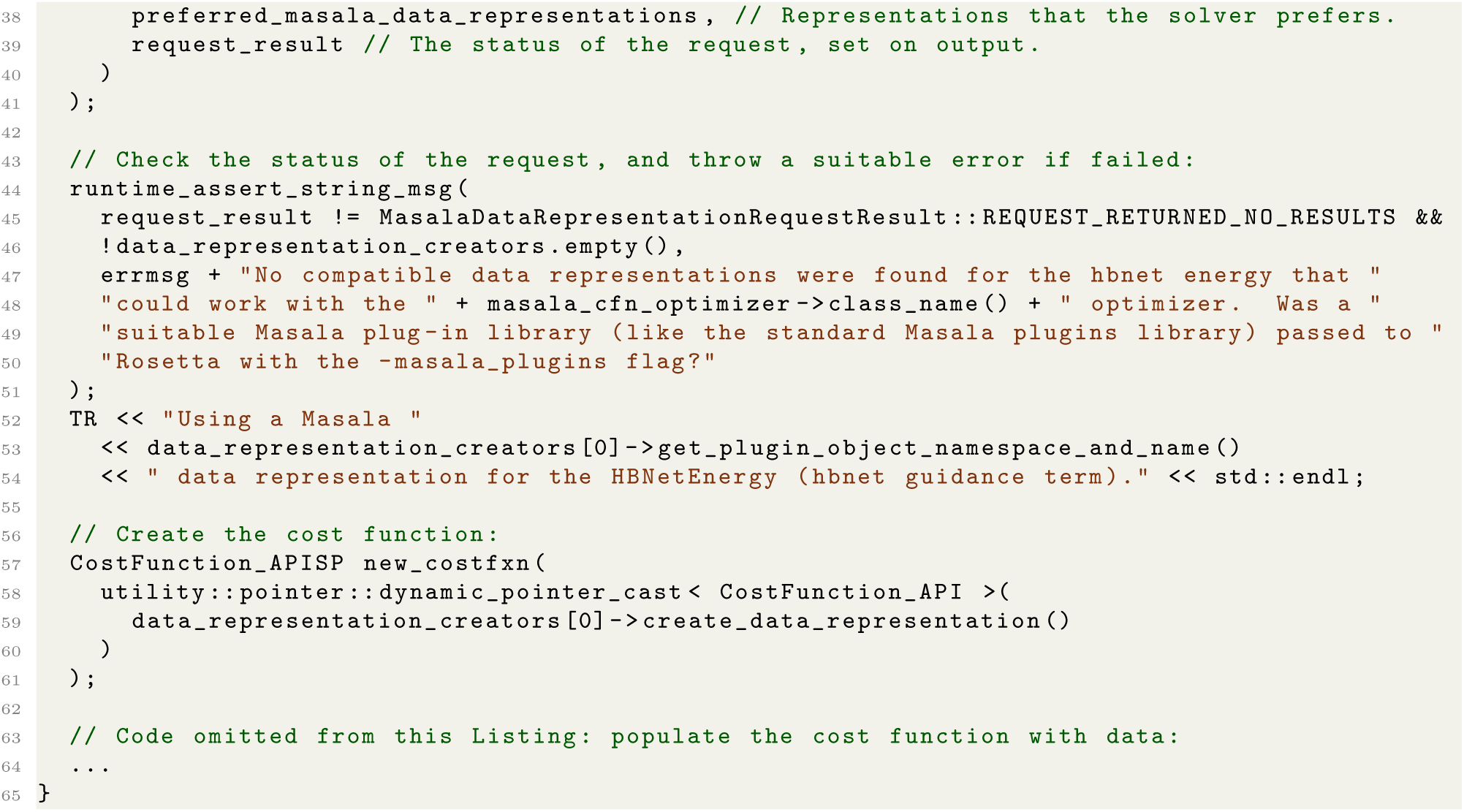

**Listing 22.** Abridged contents of the `HBNetEnergy::generate_masala_cost_functions(·)` function, showing how the code requests a data representation for a Masala cost function that is compatible with the selected Masala CFN solver engine. See `Rosetta/main/source/src/core/pack/guidance scoreterms/hbnet energy/HBNetEnergy.cc` in the version of Rosetta patched to use Masala for full detail. Note that the `get_dr_category()` function returns the hierarchical category for the cost function data representation as a vector of strings. For instance, for a quadratic penalty, the category is “CostFunction” *→* “GraphBasedCostFunction” → “GraphIslandCountCostFunction” → “SquareOfGraphIslandCountCostFunction”.

### 10.5 Adapting other design-centric guidance terms to use Masala data representations

As was the case for the `hbnet` design-centric guidance term, we adapted other design-centric guidance terms to be compatible with Masala CFN optimizers, by adding suitable Masala data representations to the Standard Masala Plugins library that could be configured by the terms when they were included in a rotamer optimization problem. These guidance terms include the `aa_composition`, `netcharge`, `buried - unsatisfied_penalty`, and `voids_penalty` terms (reviewed in [42] and [43]). When all of these terms are used in conjunction, the objective function minimized by a CFN solver is given by Eq. (11):

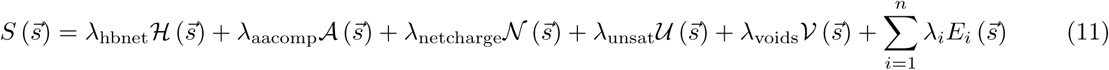

As in Eq. (9), 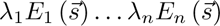 represent the other, physics-based energy terms in the scoring function, each weighted by its own Lagrange multiplier. 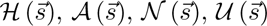 and 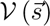 are defined in Eq. (10), above, and in Eqs. (12), (13), (14), and (15), below.

#### 10.5.1 The aa_composition and netcharge design-centric guidance terms

The `aa_composition` term and the `netcharge` term impose a user-defined penalty for deviation from a desired amino acid composition or from a desired net charge, respectively, as shown in Eqs. (12) and (13). This allows

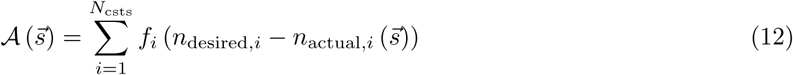

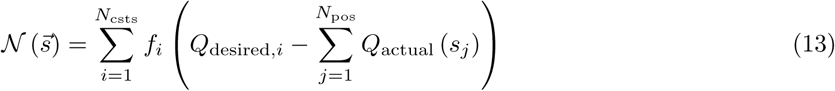

In the above, *N*_csts_ is the number of amino acid composition constraints or net charge constraints (applied globally or to regions), *N*_pos_ is the number of amino acid positions in the molecule, and *f*_1_ (*·*) *. . . f_N_*_csts_ (*·*) are user-defined functions applied to the count differences. In Eq. (12), *n*_desired*,i*_ is the desired number of amino acids of a given type or with a given set of properties for the *i^th^* amino acid composition constraint, and *n_actual,i_* 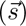 is the actual count given the current selection, 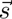. Similarly, in Eq. (13), *Q*_desired*,i*_ is the desired net formal charge for the *i^th^* region of a molecule, and *Q*_actual_ (*s_j_*) is the actual net formal charge for the rotamer selected at the *j^th^* position given the current rotamer selection 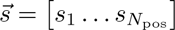.

To make these two guidance terms compatible with Masala CFN solver engines, we added a Masala cost function type called a `FunctionOfIntegerPenaltySumCostFunction`. The version compatible with the `MonteCarloCostFunctionNetworkOptimizer`, the `GreedyCostFunctionNetworkOptimizer`, the `RandomCost-FunctionNetworkOptimizer`, and other Standard Masala Plugins CFN optimizers, is defined in namespace `standard_masala_plugins::optimizers::cost_function_network::cost_function`. The `FunctionOfIntegerPenaltySumCostFunction`, like any Masala plugin or data representation, has a hierarchical category associated with it; in this case, it is “CostFunction” → “ChoicePenaltySumBasedCostFunction” *→* “IntegerPenaltySumBasedCostFunction” *→* “FunctionOfIntegerPenaltySumCostFunction”. Future equivalents of the `FunctionOfIntegerPenaltySumCostFunction`, intended to be compatible with other solvers (*e.g.* for GPU or QPU), will be implemented in the same category, though possibly in different libraries or namespaces, and certainly with different class names. By querying the `MasalaDataRepresentationManager` for a data representation in this category compatible with the chosen Masala CFN optimizer engine, the `aa_composition` and `netcharge` terms can always either prepare a suitable data representation, or throw an informative error message telling the user that these guidance functions cannot be used with the chosen optimizer engine (for want of a suitable data representation).

The `FunctionOfIntegerPenaltySumCostFunction` class implemented in the Standard Masala Plugins library permits a signed integer penalty to be assigned to each choice on each node. At scoring time, it sums the integers assigned to the selected choices across all nodes, then applies a user-defined function to the sum. In rotamer optimization problems, choices correspond to rotamers, and nodes to positions in a peptide or protein. For the `aa_composition` scoring term, the value assigned to each choice is 0 or 1, depending on whether it is a rotamer of the desired type to be counted or not; for the `netcharge` term, the value is the net formal charge of the rotamer multiplied by 0 or 1, depending on whether the rotamer is at a position in the set of positions selected for a given net charge constraint. Setters allow a series of penalty values to be provided for a range of possible values of the integer sum for the selected choices, as well as a selection of the behaviour (repeating a constant final value, extrapolating the last two values linearly, or extrapolating the last three values quadratically) outside of the range provided by the user.

Because the search performed at optimization time need only sum penalty values, `aa_composition` is greatly accelerated even on a single thread, since there is no longer any comparison of amino acid name strings or property strings during the search (see section 4.1). We suspect that the `netcharge` is the sole guidance function that didn’t show a major speed improvement because it was already summing integer formal charges, so this doesn’t greatly alter its function or single-threaded performance. Nevertheless, the ability to easily implement equivalent data representations on GPU will greatly facilitate GPU-based massively parallel peptide and protein design, and even on multi-core CPUs, this introduces the ability to parallelize the search over rotamer combinations in multiple threads.

#### 10.5.2 The buried unsatisfied penalty design-centric guidance term

Rosetta’s `buried_unsatisfied_penalty` imposes a penalty during rotamer optimization for buried hydrogen bond donors or acceptors that are not involved in hydrogen bonds. These are highly destabilizing features in the cores of folded peptides or proteins, since these chemical groups can almost certainly be satisfied in the unfolded state by hydrogen bonding with water [97]; as such they should be discouraged during design. Rosetta’s hard-coded implementation uses a graph structure, the Rosetta `BuriedUnsatPenaltyGraph` (defined in Rosetta namespace `core::pack::guidance_scoreterms::buried_unsat_penalty`) to pre-compute hydrogen bonds between all donor and acceptor groups in all candidate rotamers in the problem, then uses this graph to look up hydrogen bonding interactions during the rotamer search. At each step in the search, a penalty is computed as defined in Eq. (14). This penalty is added to the overall score, weighted by a Lagrange multiplier *λ*_unsat_, as shown in Eq. (11). Unfortunately, the `BuriedUnsatPenaltyGraph` data structure is less efficient than it could be, and its complexity hinders attempts to experiment with better ways to set up the problem for faster solving.

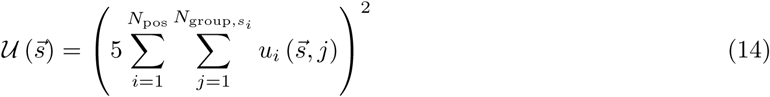

In Eq. (14), *N*_pos_ is the number of positions in the peptide or protein, *N*_group*,s*_*_i_* is the number of hydrogen-bonding groups (donors or acceptors) in the rotamer *s_i_* currently selected at the *i^th^* position, and 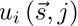 is 1 if the *j^th^* hydrogen-bonding group in the rotamer currently selected at the *i^th^* position is unsaturated or over-saturated by the overall selection 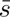, and 0 otherwise. Note that computation of 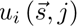 necessitates an inner loop over selected rotamers at all positions to count the number of other choices satisfying the *j^th^* hydrogen-bonding group in the rotamer selected at the *i^th^* position, meaning that efficient data representations are crucial to keep this from being extraordinarily slow.

Masala introduces “CostFunction” *→* “SumOfUnsatisfiedChoiceFeaturesCostFunction” *→* “SquareOfSumOfUnsatisfiedChoiceFeaturesCostFunction” as a general hierarchical category for data representations that define one or more “features” per choice at each node (an abstraction of the chemical groups for each rotamer at each position in a peptide or protein) and count the number of such “features” that fail to interact with one another given a particular selection of choices, squaring the count to compute the final penalty. In the Standard Masala Plugins library, the version of this cost function that is compatible with the CPU-based CFN solvers is also called the `SquareOfSumOfUnsatisfiedChoiceFeaturesCostFunction`, and is located in `standard - masala plugins::optimizers::cost function network::cost function::feature based`. This implementation uses `std::unordered map` for fast hash-based lookups of features for a given choice selection, resulting in a speedup of up to two orders of magnitude relative to Rosetta’s implementation (see section 4.1). However, as before, additional data representations may be implemented with other names; provided they are in the same data representation category, they will be seamlessly interoperable with existing design protocols allowing future efficiency enhancements. As before, additional data representations will be written for GPU- or QPU-based solvers.

#### 10.5.3 The voids penalty design-centric guidance term

The Rosetta `voids_penalty` term, described in detail in [42], discourages voids or holes in the packed core of a peptide or protein. It does this by pre-computing and caching both the volume to be filled (*V* ), and the volume of the portion of each rotamer (indexed as *j*) at each position (indexed as *i*) that lies in the volume to be filled (*v_i,j_*). Volumetric grids are used only for this pre-computation; at runtime, a penalty given by Eq. (15), based on the pre-computed and cached values, is calculated. This penalty is minimized when the total volume of the currently-selected rotamers matches the volume to be filled, implying no holes. As before, this is added to the total score, weighted by a Lagrange multiplier *λ*_voids_, as shown in Eq. (11).

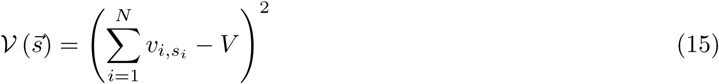

To make the `voids_penalty` guidance term compatible with Masala CFN optimizers, we added the “CostFunction” *→* “ChoicePenaltySumBasedCostFunction” *→* “SquareOfChoicePenaltySumCostFunction” category of cost function data representations. These cost functions allow a protocol-defined double-precision floating-point penalty to be assigned to each choice at each node (corresponding here to the volumes *v_i,j_* assigned to the *j^th^* rotamer at the *i^th^* position), as well as a protocol-defined constant (corresponding here to the total volume *V* to be filled). When the cost function value is computed for a choice selection *s⃗*, the penalties of the currently-selected choices are summed, the constant is subtracted, and the result is squared. For compatibility with the Masala CFN optimizers in the Standard Masala Plugins library, we added the `SquareOfChoicePenaltySumCostFunction` class in namespace `standard masala plugins::optimizers::cost function network::cost function`. This uses a `std::unordered map` for fast hash-based lookup of node choice penalty values, and permits an approximately threefold speedup over the Rosetta implementation (see section 4.1). Once again, additional data representations may be added in the future, in the same data representation category but with unique class names, to permit alternative internal structures offering greater speedups on CPU or to permit compatibility with alternative approaches on GPU, QPU, or other hardware.

### 10.6 Template preferred data representations

The platform described here has already accelerated methods development. In separate work to develop methods for designing peptides and proteins on quantum computers (see preprints [72] and [73]), we realized that, in addition to having different data representations for different types of solvers, it can be useful to have different data representations for the *same* type of solver, in order to experiment with how new ways of representing a problem can alter the ease with which a solver may find a solution. However, if there are multiple data representations compatible with a particular solver, there needs to be an easy way for a user to specify a preferred one; otherwise, the `MasalaDataRepresentationManager` could return *any* of the representations compatible with the given solver. To this end, we added a special annotator for API setter functions called a `PreferredTemplateDataRepresentationSetterAnnotation` (defined in `masala::base::api::setter::setter_annotation`). This annotation indicates that the setter accepts a preconfigured data representation that should be used as a template for all subsequent data representation requests. Rosetta has been modified to be aware of this annotation, allowing configuration of a data representation (which may have setter functions configuring options of its own) from within the XML setup tags of a Masala engine.

### 10.7 Modifications to the Rosetta MinMover to use Masala RVL solver engines

The Rosetta `MinMover` calls the Rosetta `minimizer` to perform gradient-descent minimization in order to relax molecular structures, moving geometry to find a nearby local minimum in the energy landscape [1]. It does this typically using the L-BFGS quasi-Newtonian method to approximate the inverse of the Hessian matrix, with line searches using the Armijo-Goldstein convergence criterion as a sub-algorithm [81, 85]. As was the case for the `PackRotamersMover`, our Rosetta patch modifies the `MinMover` to provide the option of using Masala real-valued local optimization (RVL) solver engines, which can be provided in runtime-linked Masala plugins, in place of the internal Rosetta `minimizer`. As before, the default behaviour of the `MinMover` remains to use the Rosetta `minimizer`.

The modifications follow the same pattern as those made for the Rosetta `PackRotamersMover`, and make use of the `generate_xml_schema_for_masala_plugin_object tag(·)` function shown in Listing 17 and the `parse_masala_setters_function(·)` shown in Listing 18 to modify the XML schema definition describing the MinMover’s RosettaScripts interface and to parse user inputs to configure the MinMover, respectively.

The `FastRelax` and `FastDesign` movers, which perform alternating calls to the `PackRotamersMover` and the `MinMover`, have also been modified so that the user may specify that Masala RVL solvers should be used during continuous minimization steps, just as they were modified to allow users to specify that Masala CFN solvers should be used during discrete rotamer optimization steps (described in section 10.3). Please refer to the source code in `Rosetta/main/source/src/protocols/minimization packing/MinMover.cc` and in `Rosetta/main/source/src/protocols/relas/FastRelax.cc` in the patched version of Rosetta for the changes. Additionally, users may provide a default `MinMover` setup script (in RosettaScripts XML format) by passing the `-default_masala_minmover_configuration <CONFIGURATION file name>` command-line flag to any Rosetta application, allowing a Masala RVL optimizer to be specified for use in any Rosetta protocol that would otherwise use the `minimizer`.

### 9.8 Adding new functionality through Masala plugin modules

As alluded to in the previous sections, the modifications to Rosetta (or similar modifications that can be made to any other software package) permit easy development of new solver engines or new data representations for more efficiently solving problems, with immediate deployment in production protocols without the need to refactor or even recompile Rosetta (or other calling code). Through the hierarchical category definitions, it is possible to classify plugin modules so that particular types can be requested. The slight disadvantage of this system is that a developer loses the usual strong guarantees in C++ about the type and inheritance of each class, which would result in errors thrown by the compiler if any assumptions were violated. This necessitates some runtime checks of object types after dynamic casting. The advantage, though, is versatility. Adding a new plugin Masala module is as simple as the following:

1. Create header (`.hh`), forward declaration (`.fwd.hh`) and implementation (`.cc`) files for the new class in some directory in an existing sub-library of an existing Masala plugin library. (Alternatively, new libraries can be created.)
2. Create the new class, having it inherit from `MasalaPlugin` or one of its sub-classes. If the class is an engine, it should inherit from `MasalaEngine` or one of its sub-classes (such as `PluginCost-FunctionNetworkOptimizer` or `PluginRealValuedFunctionLocalOptimizer`). If the class is a data representation, it should inherit from `MasalaDataRepresentation` or one of its sub-classes (such as `PluginPairwisePrecomputedCostFunctionNetworkOptimizationProblem`).
3. Implement the functionality for the new class, in one or more work functions. Note that some parent classes may have work functions for you to define, such as `PluginCostFunctionNetworkOptimizer::run cost function network optimizer(·)`.

(a) If this new class is a `MasalaPlugin`-derived class (which it must be, in order to work with the Masala plugin system), ensure that plugin category and keyword functions (`get categories()` and `get keywords()`) are implemented.
(b) If this new class is a `MasalaEngine`-derived class, ensure that engine category and keyword functions (`get engine categories()` and `get engine keywords()`) are implemented as well. Note that since `MasalaEngine` derives from `MasalaPlugin`, the `get engine categories()` and `get engine - keywords()` functions could simply call `get categories()` and `get keywords()` (which must also be implemented) if the categories and keywords are the same.
(c) If this new class is a `MasalaDataRepresentation`-derived class, ensure that data representation category and keyword functions (`get data representation categories()` and `get data representation keywords()`) are implemented. Since `MasalaDataRepresentation` derives from `MasalaPlugin`, these functions could call `get categories()` and `get keywords()` (which must also be implemented) if the categories and keywords are the same. In addition, ensure that `get - compatible masala engines()`, `protected empty()`, `protected reset()`, `protected clear()`, and `protected_make_independent()` are all implemented.
(d) If this new class is an `OptimizationProblem`, in addition to ensuring that all `MasalaPlugin` and `MasalaDataRepresentation` functions are implemented, ensure that `protected_finalize()` is implemented. This function permits optional reorganization of internal data after all data have been loaded into the class.
4. Add suitable setters and getters so that your class may be configured.
5. Write the `get_api_definition()` function for your class, and ensure that any setters and work functions that you want to include in the public API are included.
6. Add your new class to the `generate_api_classes()` function in the `api/` subdirectory of the library in which you have added your class.
7. Add unit tests to the `tests` directory. (Optional but strongly recommended.)
8. Build your Masala plugin library using the `buildme.sh` script.
9. Ensure that, at runtime, the Masala plugin library’s path is passed to Rosetta (or other external software that has linked Masala’s Core library at compilation time). API definitions will automatically be used to add user interface elements (see section 3.4) for your new plugin module, permitting it to be used in existing protocols with just a few lines added to the protocol script.

## 12 Acknowledgements

TZ is funded by the Natural Sciences and Engineering Research Council of Canada (NSERC), through a Canada Graduate Scholarship (Doctoral), Discovery Grant, and Collaborative Research and Training Experience (CREATE) Quantum Computing Doctoral Fellowship. NA and PH are funded by NIH grant DP2GM146249. QZ, SMBAT, PDR, and VKM are wholly funded by the Simons Foundation. We are grateful to Alex Ford for implementing the `ResidueArrayAnnealableEnergy` in Rosetta, and to Chris Edelmaier, Géraud Krawezik, Benjamin P. Brown, and Bryce Palmer for stimulating discussion and advice on algorithms and software development. Thanks are also due to Ekaterina Maximova, Allon Goldberg, Ella King, Mariah Culpepper, and Jacqueline Seal for using Masala for design as it was developed, and for diligently reporting bugs. We also thank the Flatiron Institute’s Scientific Computing Core for ongoing support. The computations reported in this paper were performed using resources made available by the Flatiron Institute. The Flatiron Institute is a division of the Simons Foundation. Additional benchmarking was performed on the Aurora supercomputer, for which an award of computer time was provided by the INCITE program. This research used resources of the Argonne Leadership Computing Facility, which is a DOE Office of Science User Facility supported under Contract DE-AC02-06CH11357.

